# Neural circuitry of dialects through social learning in Drosophila

**DOI:** 10.1101/511857

**Authors:** Balint Z Kacsoh, Julianna Bozler, Sassan Hodge, Giovanni Bosco

## Abstract

Drosophila species communicate the presence of parasitoid wasps to naïve individuals. This observation suggests a rudimentary Drosophila social structure. Communication between closely related species is efficient, while more distantly related species exhibit a dampened, partial communication. Partial communication between some species is enhanced following a period of cohabitation, suggesting that species-specific variations in communication “dialects” can be learned through social interactions. However, it remains unclear as to how the behavioral acquisition and how learning dialects is facilitated by distinct brain regions. In this study, we have identified six regions of the Drosophila brain essential for dialect learning, including the odorant receptor Or69a. Furthermore, we pinpoint subgroups of neurons such as motion detecting neurons in the optic lobe, layer 5 of the fan-shaped body, and the D glomerulus in the antennal lobe, where activation of each are necessary for dialect learning. These results demonstrate that Drosophila can display complex social behaviors with inputs to multiple regions of the Drosophila brain and unique subsets of neurons that must integrate olfactory, visual and motion cues.

## Introduction

The ability to decipher, unravel, and react to environmental information is a phenomenon found throughout all life forms. Organisms benefit from receiving information regarding stressors and/or advantageous environmental conditions. From bacteria to plants and to mammals, information transfer can occur within and between species. Social communication is most extensively documented in more complex organisms such as mammals and birds. However, insects can also display a broad range of sociality. When one observes insect groups in nature, their behavior readily reveals distinct planes of cooperation, communication, and social structure (Wilson, 1997). Eusocial insects provide insight to a high level of complex social structure. For example, one can observe honeybees gathering nectar from a flowerbed. While this might seem a rudimentary and relatively simple behavior, this forager was guided to this particular flower by nest mates’ dance, which contains symbolic information about distance, direction, and food source quality (Beekman et al, 2006, Breed et al, 2004, Chittka & Geiger, 1995, Crist, 2004, Gould, 1974). Ants also demonstrate a highly complex social structure comprised of multiple castes and accompanying behavior (Hölldobler & Wilson, 2009, Wilson, 1983, Wilson, 1971). In some species, when a queen has recently been lost, worker ants begin to antenna-duel for the prize of becoming the new queen. The winner converts into a queen, acquires an extended lifespan, and becomes reproductively active by means of epigenetic changes(Yan et al, 2017). This queen contest is dependent on many social cues and interactions including winning the antenna duel, but also acceptance of winners and losers. Clearly, multiple signals must be shared among individual ants for a queen to suppress workers from becoming a queen, while, in the absence of a queen, different inputs allow for only one worker to become a queen.

*Drosophila melanogaster*, in conjunction with other Drosophila species, have provided insights into a wide array of biological processes, including the mechanisms of learning, memory, and other multifaceted behaviors (Bier, 2005, Helfand & Rogina, 2003). The once canonically “asocial” fruit fly can be utilized to study rudimentary social interactions, including social learning where information transfer occurs between two individuals (Battesti et al, 2012). Another fundamental study of sociality in Drosophila is the behavior and associated neural circuitry of mating (Lebreton et al, 2017a). Mating requires recognition of a receptive individual by visual and olfactory cues. Machine learning has been employed to demonstrate the ability of Drosophila to use the visually distinct features of other individuals to determine identity within a group setting— an important finding supporting the possibility that a rudimentary social structure exists in Drosophila (Schneider et al, 2018). Oviposition site and food selection are also driven by social exchange mediated by olfactory cues and neural processing (Lin et al, 2015). Recent work has also demonstrated the importance of the communal aspect of the Drosophila lifecycle involving tumor-genesis, where Drosophila are able to discriminate between individuals at different stages of tumor progression and prefer social environments of flies without tumors (Dawson et al, 2018). It has also been suggested that social learning in Drosophila can lead to persistent mate-choice reminiscent of cultural transmission, and that such choices can in turn be inherited through social learning from one generation to the next (Danchin et al, 2018, Danchin et al, 2010). Thus, there is mounting experimental evidence that Drosophila could be useful as models for understanding genetic and neurophysiological mechanisms of social learning as well as evolution of social structures.

Adult Drosophila across the genus have been in an arms race against predators and have subsequently evolved complex immune and behavioral changes to protect their offspring from endoparasitoid wasps. These wasps can prey on immature stages of larvae of certain Drosophilid species (Bozler et al, 2017, Driessen et al, 1989, Edmondson & Wyburn, 1963, Fleury et al, 2004, Kacsoh & Schlenke, 2012, Kacsoh et al, 2014a, LaSalle, 1993, Mortimer et al, 2012, Mortimer et al, 2013, Rizki, 1957, RIZKI & RIZKI, 1959, Schlenke et al, 2007a, Wigglesworth, 1959). These behavioral changes include, but are not limited to, an altered food preference and reduced oviposition (egg-laying) (Kacsoh et al, 2018a, Kacsoh et al, 2018b, Kacsoh et al, 2013, Kacsoh et al, 2015a, Kacsoh et al, 2015c, Kacsoh et al, 2017, Lefevre et al, 2012). The wasp predator threat triggers social transference of information between experienced and naïve individuals. Wasp-exposed teacher flies communicate the wasp threat to naïve flies through visual cues (Kacsoh et al, 2015c), followed by naïve student flies then depressing their own egg-laying rate. This communication can occur within a species, but it also occurs between related species. Closely related species demonstrate an efficient level of communication. A dampened, or partial communicative ability is observed when more distantly related species are paired (Kacsoh et al, 2018b). Very distantly related species lack the ability to communicate. Remarkably, the observed partial communication between some species is alleviated following a cohabitation period where an exchange of visual and olfactory signals is enabled. This observation is highly suggestive of natural variations in modes of communication, similar to the evolution of linguistic variations between species, termed “dialects” (Kacsoh et al, 2018b, Manak, 2018).

Each of the behaviors outlined above is achieved with a brain the size of a grain of sand, yet, these Drosophila demonstrate a remarkable level of neuronal plasticity influenced by life history and socialization (Avarguès-Weber et al, 2011, Chittka & Niven, 2009, Dyer, 2012, Dyer, 1996, Giurfa, 2003, Greggers & Menzel, 1993, Srinivasan, 2010, Zhang, 2012, Zhang et al, 2007). In ants, bees, and even Drosophila, we are privileged to not only see how complex societies evolve independently of humans, but also to dissect, with ever increasing clarity, the relationship between advanced social order, the forces of natural selection that shaped them, and the underlying neuro-genetic, epigenetic, and genetic mechanisms that guide these behaviors.

In this study we focus on neural circuitry of dialect learning in the Drosophila system: Interspecies dialect learning, unlike intraspecies social learning, is multi-modal, requiring, at minimum, olfactory, visual, sex-specific, temporal, neuronal, and ionotropic cues for successful dialect acquisition (Kacsoh et al, 2018b, Kacsoh et al, 2015c). Considering the ecological niches of Drosophila, where these insects do not exist in a monoculture, brings to the forefront the highly social nature of these insects(Markow, 2015). Using this fly-fly social learning paradigm, we asked: (1) what neuronal groups are required for interspecies dialect learning; (2) what is the subset of neurons in a given neuronal structure that is necessary for the behavior; and (3) what is the subset of neurons in a given neuronal structure that is sufficient, when activated, to drive dialect learning.

## Results

### Dialect learning is governed by multiple brain regions

In order to ascertain communication ability, we utilized the fly duplex. The fly duplex is an apparatus with two transparent acrylic compartments allowing one to test whether *D. ananassae* responds to seeing predators (acute response) and if exposed “teacher” female flies can communicate this threat to naïve unexposed *D. melanogaster* “student” female flies (Kacsoh et al, 2015c). The fly duplex allows flies to see other flies or wasps in the adjacent compartment, without direct contact, making communication exclusively visual (Figure 1 A). We place ten female and two male *D. ananassae* into one duplex compartment, with an adjacent compartment containing (or not for control) twenty female wasps. Following a 24-hour exposure, wasps are removed and acute response is measured by counting the number of eggs laid in the first 24-hour period in a blinded manner. Flies are shifted to a new duplex, with ten female and two male *D. melanogaster* naïve student flies in the adjacent compartment (Figure 1 A, see methods). Subsequent to a second 24-hour period, all flies are removed and the response of both teacher and student is measured by counting the number of eggs laid in a blinded manner (see methods). The 24-48-hour period measures memory of teachers having seen the wasps and students having learned from the teachers. As previously shown, using wild-type *D. anananassae*, we find both an acute response and a memory response to the wasp in teacher flies and a partial learned response in naïve *D. melanogaster* student flies (Supplementary Figure 1, Supplementary File 1 for all raw egg counts and p values) (Kacsoh et al, 2018b, Kacsoh et al, 2015c, Lefevre et al, 2012, Lynch et al, 2016). We define partial communication as oviposition depression of students paired with wasp-exposed teachers being ~50-65% of unexposed flies and statistically non-significant from the ~50% oviposition rate. Next, we recapitulated a previous finding demonstrating the ability of *D. melanogaster* to enhance its communication ability with *D. ananassae* following a weeklong cohabitation termed “dialect learning” (Supplementary Figure 1) (Kacsoh et al, 2018b). We define this enhanced communication ability as egg-laying by students paired with wasp-exposed teachers being ~10-30% compared to unexposed and statistically different from ~50% oviposition rate. Using this paradigm, we wished to elucidate the neural circuitry that governs this complex behavior by individually deactivating defined subsets of neurons in an inducible fashion. Understanding dialect learning, storage, and retrieval requires a fundamental knowledge of the underlying neuronal circuits, which is currently unknown.

**Figure 1.**
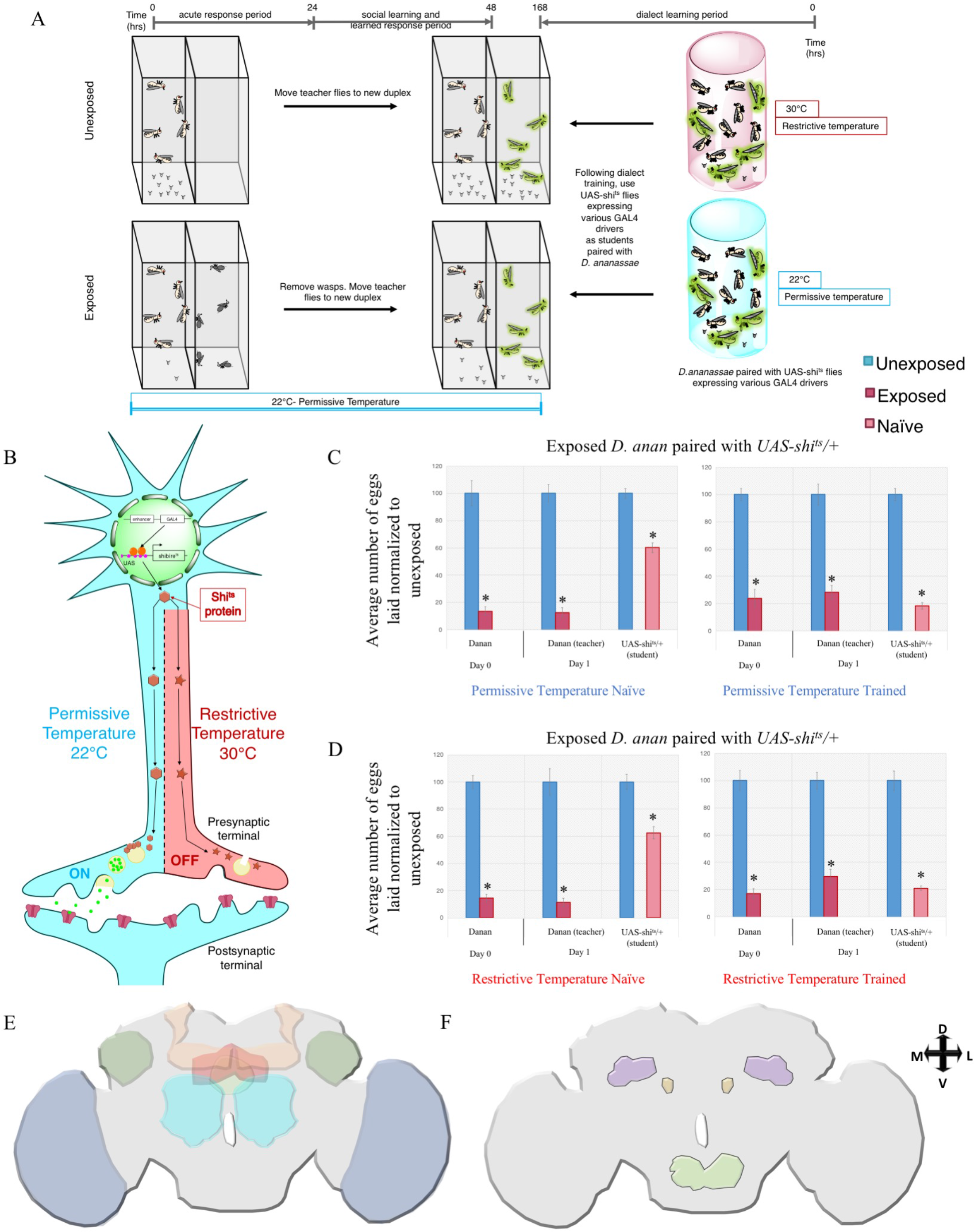
Dialect learning is governed by multiple brain regions revealed by UAS-shits. (A) Standard experimental design using UAS-shibire (UAS-shits) in conjunction with various GAL4 drivers. Dialect learning is performed at either the permissive (22°C) or restrictive (30°C) temperature, while the wasp exposure and social learning period is performed exclusively at the permissive temperature. (B) Schematic of the UAS-shits expression in neurons, where the restrictive temperature turns off the neurons of interest and the permissive temperature does not affect these neurons. (C) Percentage of eggs laid by exposed flies normalized to eggs laid by unexposed flies is shown. UAS-shits outcrossed to *Canton S* trained by *D. ananassae* at either the permissive temperature (C) or the restrictive temperature (D) show wild-type dialect acquisition in both the naïve and trained states. Error bars represent standard error (n = 12 biological replicates) (*p < 0.05). (E) Brain regions identified as involved in dialect learning—the optic lobe (blue), mushroom body (orange), antennal lobe (teal), lateral horn (green), fan-shaped body (red), and ellipsoid body (light green). (F) Brain regions identified as being dispensable for dialect learning— the prow (light green), bulb (yellow), and superior clamp (purple).

In order to identify brain regions that govern dialect learning, we utilized the FlyLight database, from which we selected GAL4 drivers that marked singular structures both strongly and ubiquitously (Jenett et al, 2012a). The FlyLight GAL4 lines were generated by insertion of defined fragments of genomic DNA that serve as a transcriptional enhancer. We selected 9 brain structures to analyze: the optic lobe, mushroom body, antennal lobe, lateral horn, fan-shaped body, ellipsoid body, prow, superior clamp, and the bulb (Couto et al, 2005a, Hsu & Bhandawat, 2016, Yu et al, 2013). If available, we obtained two unique driver lines matching our selection criteria. Using mCD8-GFP as a reporter, immunofluorescence reveals GFP expression constrained to these regions of interest, justifying these lines for further testing (Supplementary Figures 2–15 A-B).

Following identification of GAL4-driver lines marking unique brain structures, we utilized Gal4-mediated expression of the temperature sensitive *shibire*, UAS-shibire^ts^ (UAS-shi^ts^) in order to inducibly deactivate defined subsets of motion sensing neurons during dialect training (Kitamoto, 2001). Dialect training was performed at either the restrictive (30ºC) or permissive (22ºC) temperatures, after which trained or naïve *D. melanogaster* were paired with either unexposed or wasp-exposed *D. ananassae* at the permissive temperature only (Figure 1 A). This experimental setup means that only during the dialect training period is there an activation of the UAS-shi^ts^ construct, and subsequent inactivation of a defined region of interest (Figure 1 B). Wild-type flies demonstrate normal dialect learning ability and untrained states when incubated at either 22ºC or 30ºC for one week, indicating temperature shifts alone during the cohabitation period do not adversely affect dialect learning (Supplementary Figure 1). Additionally, outcrossed UAS-shi^ts^ (UAS-shi^ts^/+) demonstrate wild-type untrained and dialect-trained states at both the restrictive and permissive temperatures (Figure 1 C-D).

Previous results have demonstrated that visual system and mushroom body (MB) are required for dialect learning (Kacsoh et al, 2018b). This was demonstrated by dialect training (cohabitation) in the dark, resulting in no dialect learning. In addition, the GAL4 Gene-Switch system to transiently express tetanus toxin light chain (UAS-TeTx) specifically in the MB of *D. melanogaster* (to inhibit synaptic transmission during dialect training) also resulted in no dialect learning (Kacsoh et al, 2018b, Mao et al, 2004). Recent work has also highlighted direct neural pathways, which convey visual information from the optic lobe to the MB (Vogt et al, 2016). As a proof of concept of our experimental design, we tested two optic lobe and MB drivers in conjunction with UAS-shi^ts^. Since previous observations indicated both MB and visual system function were essential for dialect learning, the expectation was that temperature induced inactivation of both brain regions should recapitulate observations acquired with UAS-TeTx. When dialect training is performed at the restrictive temperature, both optic lobe and MB Gal4 lines driving expression of UAS-shi^ts^ result in perturbed dialect learning (Supplementary Figures 2–5). This was true for two unique Gal4 driver lines for each MB and optic lobe regions, while at the permissive temperature flies were exhibited wild-type dialect learning (Supplementary Figures 2–5). These results validate our experimental approach, further suggesting that dialect learning is in part governed by visual inputs and MB-dependent learning and memory circuitry (Aso et al, 2014, Takemura et al, 2017).

Using the approach where we utilize Gal4-mediated expression of UAS-shi^ts^, we identify four additional brain regions that govern dialect learning, where we see perturbed dialect learning at the restrictive temperature: the antennal lobe (Supplementary Figure 6–7), lateral horn (Supplementary Figure 8), fan-shaped body (Supplementary Figure 9–10), and ellipsoid body (Supplementary Figure 11–12). We find three regions, the bulb (Supplementary Figure 13), prow (Supplementary Figure 14), and superior clamp (Supplementary Figure 15), whose activation is dispensable for dialect learning. Collectively, we identify 6 regions of the brain that require activation during dialect learning (Figure 1 E), and 3 regions that do not (Figure 1 F). The finding that 3 structures could be inactivated and still result in dialect learning suggests that inactivation of any brain structure is not sufficient to perturb dialect learning. This further suggests that the regions we did identify are important and are appropriate candidates for further circuit dissection.

### Dialect learning is partially mediated by Or69a

Following the observation that the antennal lobe is required for dialect learning (Supplementary Figures 6–7), we wished to further dissect this brain region and identify a more precise series of neurons and possible pheromone candidates involved in dialect learning. Previous data has demonstrated a role for *Orco* in dialect learning—specifically that *Orco* function is a necessary during dialect training (Kacsoh et al, 2018b). The majority of olfactory receptors in Drosophila require a co-receptor for wild-type function, including *Orco* (Or83b) for odorant receptors (Larsson et al, 2004). In nature, Drosophila can utilize volatiles to transmit information across both long and short distances (Greenfield, 1981, Wicker-Thomas, 2007, Wyatt, 2014, Wyatt, 2010). Given this observation, we wished to identify a more specific odorant receptor that is necessary for dialect learning and the accompanying glomerulus that is enervated.

The Drosophila olfactory system is comprised of four sensillum classes—antennal trichoids, antennal basiconics, antennal coeloconics, and palp basiconics. These detectors are present on the third antennal segment and the maxillary palp (Figure 2 A). Housed within these sensillar classes are olfactory sensory neurons (OSNs), of which there are approximately 1300, that project from sensilla to innervate the glomeruli of the antennal lobe (Figure 2 A) (Vosshall et al, 1999, Vosshall et al, 2000, Vosshall, 2001, Vosshall & Hansson, 2011). Olfactory glomeruli are morphologically conserved, spherical compartments of the olfactory system located in the antennal lobe. These 52 glomeruli are distinguishable by their chemosensory repertoire, position, and volume. Glomeruli receive information regarding olfactory detecting from responses originating from OSNs in the sensilla (Grabe et al, 2016). Individual OSNs express only one receptor gene in conjunction with the *Orco* co-receptor, providing specificity. Detected information is then subsequently relayed to other regions of the brain, i.e. to the mushroom body signaling to the lateral horn by projection neurons. OSNs are housed in each of the four sensilla classes (Fishilevich & Vosshall, 2005).

**Figure 2.**
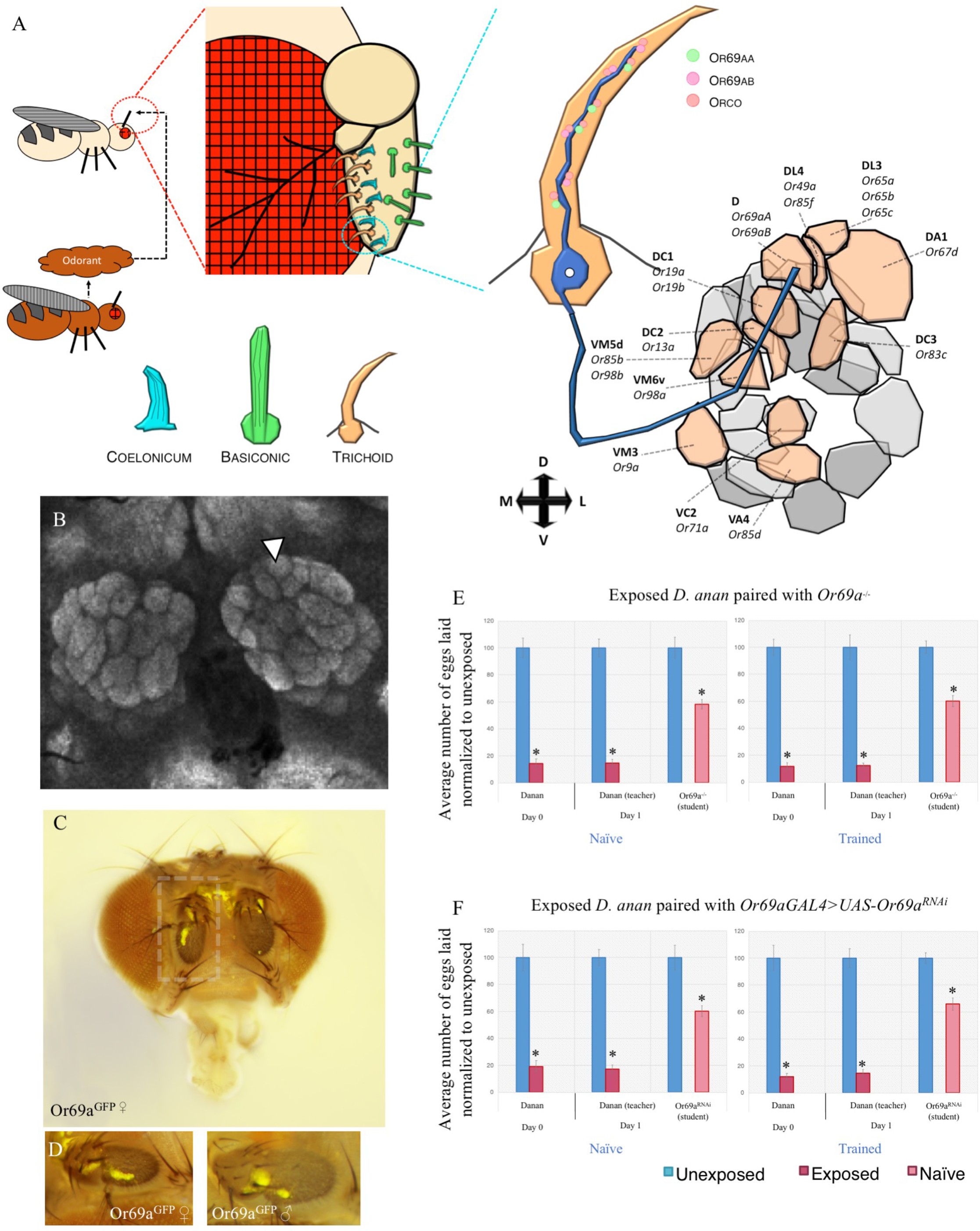
Dialect learning mediated by the odorant receptor Or69a. (A) Schematic of Drosophila antenna which has three various olfactory detecting centers: coelonicum, basiconic, and trichoid. Or69a is highlighted in the closeup and its enervation of the D glomerulus in the antenna lobe is highlighted. Glomeruli in the same focal plane are colored orange, while glomeruli in a different focal plane are colored shades of grey. (B) The Drosophila antennal lobe with the D glomerulus indicated with an arrowhead. Nc82 staining shown in grey. (C-D) Expression of Or69a using the Or69a^GFP^ construct highlighting expression in the antenna. Represented in (C) is a female fly head, while (D) represents magnified antennae (shown in box) of both female and male flies expressing Or69a^GFP^. (E) Percentage of eggs laid by exposed flies normalized to eggs laid by unexposed flies is shown. Or69a^−/−^ show wild-type naïve behavior, but are unable to learn the dialect from *D. ananassae* following training. (F) Or69a^GAL4^ driving Or69a^RNAi^ shows wild-type naïve behavior, but is unable to learn the dialect from *D. ananassae* following training. Error bars represent standard error (n = 12 biological replicates) (*p < 0.05).

We utilized the current knowledge of the Drosophila olfactory system to identify potential olfactory receptor candidates that might be involved in dialect learning. In particular, we identify Or47b, Or65a, and Or69a as potential candidates, which have been demonstrated to play a role in other social interaction based assays (Lebreton et al, 2017b, Lone & Sharma, 2012, van Naters, Wynand van der Goes & Carlson, 2007). The glomeruli enervated by each of the Ors respective OSNs are also marked in both antennal lobe drivers we utilized previously, making them suitable candidates (Supplementary Figure 6–7).

The sensillum that houses the Or47b receptor enervates the VA1lm glomerulus of the Drosophila antennal lobe (Supplementary Figure 16 D, F) (Fishilevich & Vosshall, 2005). Previous studies have demonstrated that Or47b receptor and receptor neurons are necessary for social-sexual interactions, such as locomotor activity and nocturnal sex drive (Lone & Sharma, 2012). Or47b in males drives responses to virgin females, and in females drives responses to male cuticular extract (van Naters, Wynand van der Goes & Carlson, 2007). Given that both males and females are required in dialect learning (Kacsoh et al, 2018b), we hypothesized that Or47b would be a suitable candidate for testing. We find that flies mutant in Or47b (Or47b^−/−^) and RNAi (Or47b^RNAi^) expressing *D. melanogaster* targeting Or47b using an Or47b-GAL4 results in wild-type dialect learning, suggesting that dialect learning is not directly mediated by Or47b (Supplementary Figure 16 E, G).

The sensillum that houses the Or65a receptor enervates the DL3 glomerulus of the Drosophila antennal lobe (Figure 2 A, Supplementary Figure 16 H) (Fishilevich & Vosshall, 2005). Previous studies have demonstrated that the Or65a receptor and receptor neuron is involved in mate recognition. In particular, Or65a responds to 11-cis-Vaccenyl acetate (cVA), a male-specific lipid that is present on male genital material (van Naters, Wynand van der Goes & Carlson, 2007). cVA is utilized as both an attractant pheromone to an oviposition site as well as a means to dampen female mating receptivity following copulation (Datta et al, 2008, Kurtovic et al, 2007, Laturney & Billeter, 2016). We hypothesized that his complex interaction between males and females could be co-opted in dialect learning, including testing the possibility of involvement of the *fruitless* neural circuitry. We find that RNAi (Or65a^RNAi^) expressing *D. melanogaster* targeting Or65a using an Or65a-GAL4 results in wild-type dialect learning, suggesting that dialect learning is not directly mediated by Or65a, and most likely not primarily mediated by the fruitless circuit (Supplementary Figure 16 I).

Or69a is expressed in the third antennal segment (Figure 2 C-D, Supplementary Figure 16 A). Or69a has been recently deorphanized and identified to be involved in long-range, species-specific pheromone detection as well as food detection (Lebreton et al, 2017b). This dual function likely evolved as there are two isoforms of Or69a—Or69aA and Or69aB (Lebreton et al, 2017b). The sensillum that houses the Or69a receptor enervates the D glomerulus of the Drosophila antennal lobe via the ab9A OSN (Figure 2 A-B) (Couto et al, 2005a, Fishilevich & Vosshall, 2005). The compounds detected by this receptor is the cuticular hydrocarbon *Z4-11A1* that acts as a species identifier in both sexes (i.e. *D. melanogaster* is attracted but *D. simulans* is repelled). Additionally, this receptor binds kairomonal terpenoids, such as linalool or terpineol, which are found in both fruit and yeast headspace, suggesting food and oviposition site detection (Lebreton et al, 2017b). We find that flies mutant in Or69a (Or69a^−/−^) and RNAi (Or69a^RNAi^) expressing *D. melanogaster* targeting Or69a using an Or69a-GAL4 results in perturbed dialect training (Figure 2 E-F). Outcrossed UAS and GAL4 lines demonstrate wild-type dialect learning (Supplementary Figure 16 B-C). This suggests that Or69a is an essential mediator of dialect learning.

Collectively, our more in-depth analysis into the role of antennal lobe neural circuitry and its associated olfactory receptors in dialect learning reveals the role of Or69a as an olfactory receptor involved in dialect learning during communal living. Or69a has been previously demonstrated to have a dual affinity for both sex and food odorants, in a species-specific manner. Our data provide further evidence of complex integration of pheromonal cues in Drosophila that involve integration of social and habitat derived cues. We also rule out Or47b and Or65a receptors, both previously demonstrated to be involved in other social interactions. We also identify the OSN that enervates the D glomerulus in the antennal lobe as part of the neural circuit governing dialect learning.

### Dialect learning is mediated by motion-detecting circuitry

Given our finding that the optic lobe is necessary for dialect learning, we wished to further dissect this brain region and identify a more precise neuron subset. In Drosophila, motion detection requires synaptic outputs of the R1-R6 photoreceptors. These photoreceptors project their axons into the first optic neuropil, known as the lamina. This projection forms a retinotopic map of visual space (Heisenberg & Buchner, 1977, Heisenberg & Wolf, 1979, Wardill et al, 2012, Yamaguchi et al, 2008). This map is comprised of 800 columnar elements, within which R1-R6 make synaptic connections with three projection neurons. These three neurons are the lamina monopolar neurons L1, L2, and L3, as well as a local interneuron, known as amc (Figure 3 A) (Meinertzhagen & O’neil, 1991, Rivera-Alba et al, 2011). Each of these neurons has been shown to respond to different types of inputs. The L1 neuron has been shown to provide input to a pathway that detects moving light edges, while the L2 neurons provides input into a pathway that detects moving dark edges(Clark et al, 2011, Joesch et al, 2010, Joesch et al, 2013). The L3 neuron is thought to inform landmark orientation and spectral preference (Rister et al, 2007a). L1-L3 neurons represent all of the direct second order relays from R1-R6 into the medulla. Interestingly, L2 makes synaptic contacts with a third order monopolar cell, L4, which has been proposed to be involved in motion sensing, given its morphology and connectivity, though not definitively shown (Figure 3 A) (Braitenberg, 1970, Meinertzhagen & O’neil, 1991, Silies et al, 2013, Strausfeld & Campos-Ortega, 1973, Strausfeld & Campos-Ortega, 1977, Takemura et al, 2017, Zhu et al, 2009).

**Figure 3.**
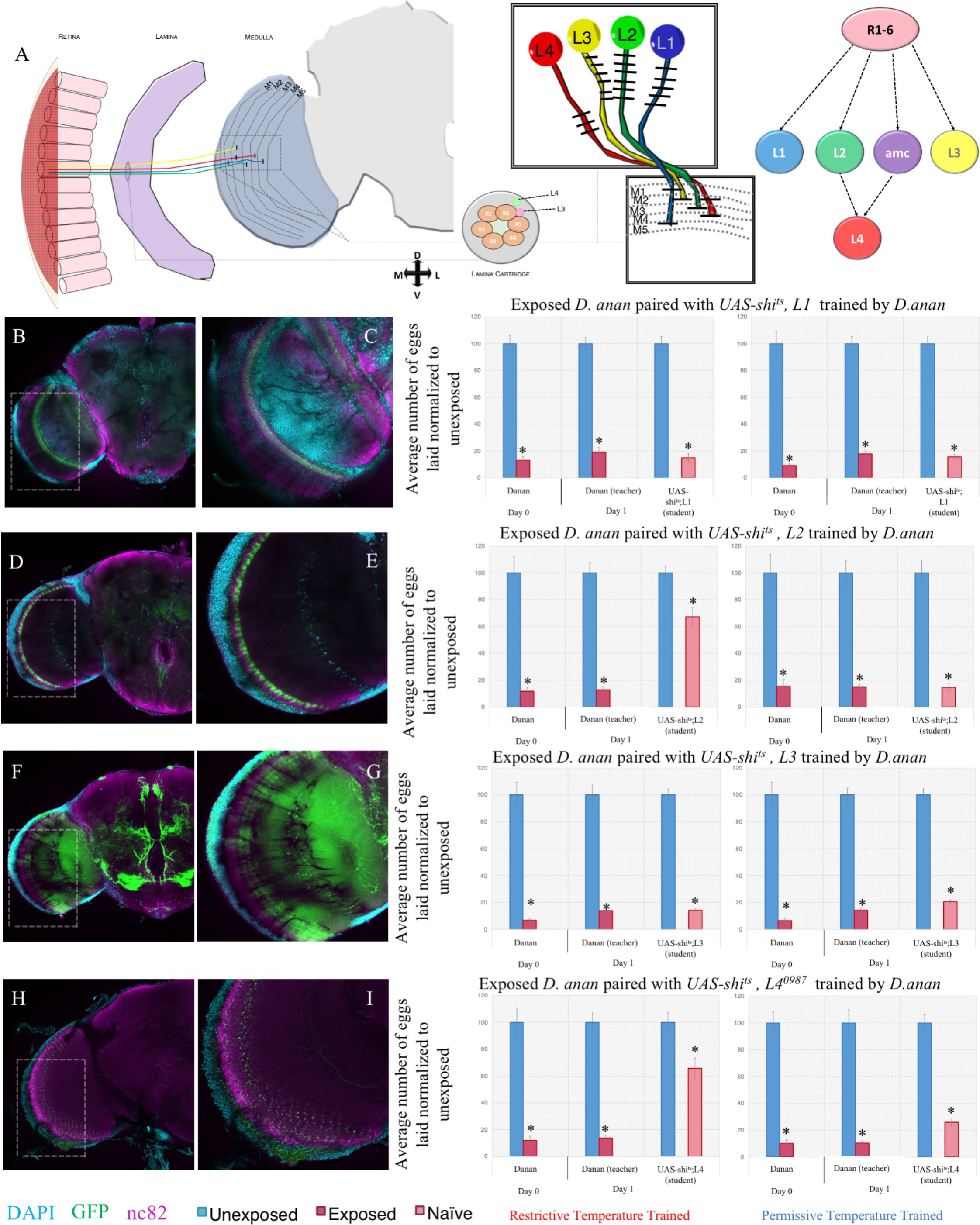
Motion detecting neurons in the optic lobe are required for dialect learning. (A) Schematic of Drosophila visual system. The optic lobe is shown highlighting the L1, L2, L3, and L4 neurons. One lamina cartridge and one medulla column are magnified to show the dendritic and axonal arborization patterns. The photoreceptors R1-R6 make synaptic connections with the L1, L2, L3, and the amc interneurons. L4 receives most inputs from L2 and amc. Dialect learning is performed at either the permissive (22°C) or restrictive (30°C) temperature, while the wasp exposure and social learning period is performed exclusively at the permissive temperature. Naïve states are shown in figure S 19. (B) Confocal image of adult brain where L1-GAL4 is driving UAS-CD8-GFP, stained with nc82 (magenta) and DAPI (teal). A magnification is shown of boxed area. See also, figure S 17. (C) UAS-shits crossed to L1-GAL4 trained by *D. ananassae* at the permissive temperature shows wild-type trained state and at the restrictive temperature shows wild-type acquisition in the trained state. (D) Confocal image of adult brain where L2-GAL4 is driving UAS-CD8-GFP, stained with nc82 (magenta) and DAPI (teal). A magnification is shown of boxed area. See also, figure S 18. (E) UAS-shits crossed to L2-GAL4 trained by *D. ananassae* at the permissive temperature shows wild-type trained state, but at the restrictive temperature shows defective acquisition in the trained state. (F) Confocal image of adult brain where L3-GAL4 is driving UAS-CD8-GFP, stained with nc82 (magenta) and DAPI (teal). A magnification is shown of boxed area. See also, figure S 19. (G) UAS-shits crossed to L3-GAL4 trained by *D. ananassae* at the permissive temperature shows wild-type trained state and at the restrictive temperature shows wild-type acquisition in the trained state. (H) Confocal image of adult brain where L4^0987^-GAL4 is driving UAS-CD8-GFP, stained with nc82 (magenta) and DAPI (teal). A magnification is shown of boxed area. See also, figure S 20. (I) UAS-shits crossed to L4^0987^-GAL4 trained by *D. ananassae* at the permissive temperature shows wild-type trained state, but at the restrictive temperature shows defective acquisition in the trained state. Error bars represent standard error (n = 12 biological replicates) (*p < 0.05).

Previous data suggests that a moving visual cue, provided by wing movement, is detected by Drosophila in the dialect-training period (Kacsoh et al, 2018b). Given this observation, we wondered whether motion-detecting neurons in the optic lobe could be responsible for acquisition of visual information dialect learning. Therefore, we utilized Gal4-mediated expression of the temperature sensitive *shibire*, UAS-shibire^ts^ (UAS-shi^ts^) in order to inducibly deactivate defined subsets of motion sensing neurons in the optic lobe during dialect training (Kitamoto, 2001). Paired controls were performed where dialect training was conducted at the permissive temperature. We utilized L1-, L2-, L3-, and L4-GAL4 lines with strong, but constrained, expression patterns (Figure 3 B, D, F, H and Supplementary Figures 17–21) (Fisher et al, 2015, Silies et al, 2013). Silencing of the L1 motion detecting neuron resulted in wild-type dialect learning, suggesting that activation of the L1 neurons are dispensable for this behavior (Figure 3 C). Silencing of the L2 motion detecting neuron resulted in the inability of *D. melanogaster* to learn the dialect, suggesting that L2 neurons are necessary for this behavior (Figure 3 E). Silencing of the L3 motion detecting neuron resulted in wild-type dialect learning, suggesting that activation of the L3 neurons are dispensable for this behavior (Figure 3 G). L4 receives most of its synaptic inputs from L2, but is also interconnected with neighboring dorso- and ventroposterior cartridges (Meinertzhagen & O’neil, 1991, Rivera-Alba et al, 2011, Takemura et al, 2017). When we silenced the L4 motion detecting neurons, we find that the ability of *D. melanogaster* to learn the dialect is perturbed, suggesting that L4 neurons are necessary for this behavior (Figure 3 I). Following this result, we tested a *splitL4-GAL4* line (L4^0980^-VP16AD, L4^0987^-GAL4DBD) (Supplementary Figure 21, Supplementary Figure 23 A). When using the *splitL4* line in conjunction with UAS-shi^ts^, we find that the ability of *D. melanogaster* to learn the dialect is perturbed, further solidifying the observation that L4 neuronal activation is necessary for dialect learning (Supplementary Figure 23 A). This finding uncovers a novel role for the L4 neurons, which have been suggested to be involved in motion detection in spatial summation by anatomical experimentation, but L4 neurons have never been implicated in a specific behavioral paradigm(Silies et al, 2013). Untrained animals at the control and restrictive temperatures behave as wild-type (Supplementary Figure 22–23).

Given that the L2 and L4 neurons are necessary for dialect learning, we wondered whether they are also sufficient. To ask this question, we expressed a temperature sensitive transient receptor potential A1 (TRPA1)(Kang et al, 2012, Lamaze et al, 2017) channel via a Gal4-mediated expression, confining expression to either the L2 or L4 neurons. Using this construct, neurons can be activated at 30ºC, while the neurons are in their steady state at 22ºC (Supplementary Figure 24 A-B). Outcrossed TRPA1/+ behave as wild-type when trained at either 30ºC or 22ºC (Lamaze et al, 2017)(Supplementary Figure 24 C-D). We activated motion-detecting neurons during the dialect-training period while keeping flies in the dark as a means to test sufficiency (Figure 4 A). Flies were kept in the dark during the training period, but all other sensory stimuli are presumably engaged (i.e. olfactory and mechanosensory interactions). At the thermally inactive temperature (basal activity level), *D. melanogaster* are unable to learn the dialect in the dark, even though the other senses are still engaged (Figure 4 C,G,E,I), recapitulating previous experiments demonstrating the need for full spectrum light (Kacsoh et al, 2018b). Activation of the either the L1, L2, or L3 motion detecting neuron results in no dialect training, suggesting that activation of these neurons is not sufficient to overcome the lack of visual cues in the dark to elicit dialect learning (Figure 4 B-G). Surprisingly, we find that activation of the L4 motion detecting neurons, using both driver lines, results in the ability of *D. melanogaster* to learn the dialect even when in the dark (Figure 4 H-I, Supplementary Figure 26 A). Untrained flies behave as wild-type, with no dialect acquisition, when kept at either temperature in the dark (Supplementary Figure 25, Supplementary Figure 26 B). Given that senses other than visual cues are stimulated when training is performed in the dark, and that activation of the L4 neurons results in dialect learning in the dark experimental setup, we conclude that the activation of L4 neurons is sufficient for dialect learning, but only with respect to the visual circuit.

**Figure 4.**
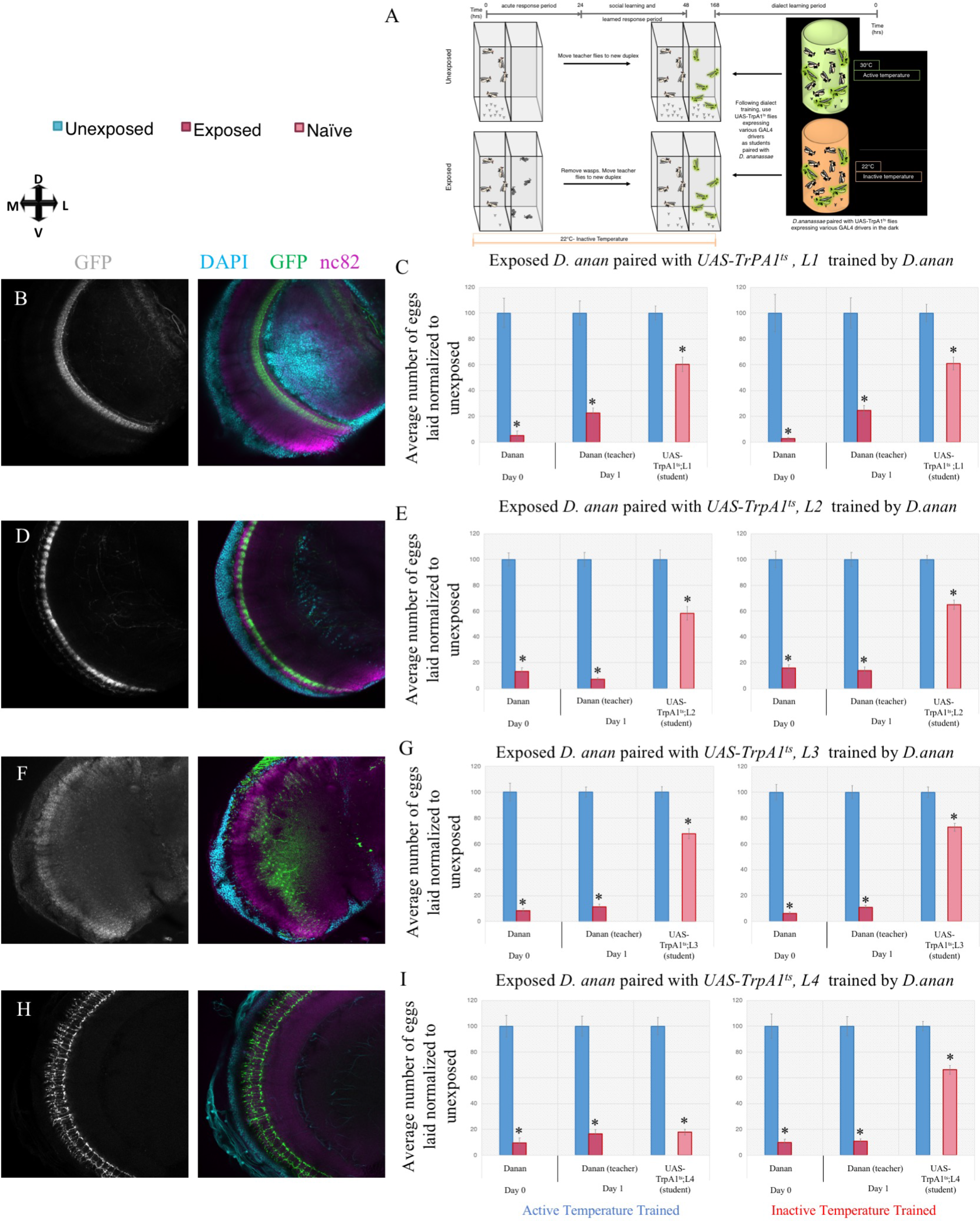
Motion detecting neurons in the optic lobe are sufficient for dialect learning. (A) Experimental design using UAS-TrpA1^ts^ in conjunction with various GAL4 drivers to test the role of motion detecting neurons in the optic lobe. Dialect learning is performed at either the active (30°C) or inactive (22°C) temperature exclusively in the dark, while the wasp exposure and social learning period is performed exclusively at the restrictive temperature. Naïve states are shown in figure S 21. (B) Confocal image of adult brain where L1-GAL4 is driving UAS-CD8-GFP, shown in white. A merged color image, showing staining with nc82 (magenta) and DAPI (teal) is also shown. (C) UAS-TrpA1^ts^ crossed to *L1-GAL4* trained by *D. ananassae* at the activation temperature shows no acquisition of the dialect and no acquisition at the inactive temperature, which demonstrates the need for full spectrum light and that L1 neurons are not sufficient to drive the behavior. (D) Confocal image of adult brain where L2-GAL4 is driving UAS-CD8-GFP, shown in white. A merged color image, showing staining with nc82 (magenta) and DAPI (teal) is also shown. (E) UAS-TrpA1^ts^ crossed to *L2-GAL4* trained by *D. ananassae* at the activation temperature shows no acquisition of the dialect and no acquisition at the inactive temperature, which demonstrates the need for full spectrum light and that L2 neurons are not sufficient to drive the behavior. (F) Confocal image of adult brain where L3-GAL4 is driving UAS-CD8-GFP, shown in white. A merged color image, showing staining with nc82 (magenta) and DAPI (teal) is also shown. (G) UAS-TrpA1ts crossed to L3-GAL4 trained by D. ananassae at the activation temperature shows no acquisition of the dialect, which demonstrates the need for full spectrum light and that L3 neurons are not sufficient to drive the behavior. (H) Confocal image of adult brain where L4^0987^-GAL4 is driving UAS-CD8-GFP, shown in white. A merged color image, showing staining with nc82 (magenta) and DAPI (teal) is also shown. (I) UAS-TrpA1^ts^ crossed to *L4^0987^-GAL4* trained by *D. ananassae* at the activation temperature shows acquisition of the dialect, but not at the inactive temperature, which demonstrates the need for full spectrum light. Error bars represent standard error (n = 12 biological replicates) (*p < 0.05).

Collectively, our genetic analysis of the optic lobe neurocircuitry and its involvement in dialect learning reveals that the visual stimulus that is detected during dialect learning is primarily detected by the dark edge detecting L2 circuit that enervates the L4 neuron, both of which are necessary for dialect learning (Figures 3 and 4). When dialect training is performed in the dark, activation of the L4 neuron is sufficient for dialect learning (Figure 4).

### Dialect learning is mediated by region 5 of the fan-shaped body

Given our observation that the fan-shaped body is necessary for dialect learning, we wished to further dissect this brain region and identify a more precise neuron subset. The fan-shaped body (FB) is located at a centralized region of the fly brain that is organized into multiple layers. As a member of the central complex(Hanesch et al, 1989), the FB is reported to be important for multiple functions, including control of locomotion(Strauss, 2002), visual feature recognition (Liu et al, 2006) and visual information processing (Weir & Dickinson, 2015), courtship maintenance (Sakai & Kitamoto, 2006), and sleep regulation(Berry et al, 2015, Donlea et al, 2011, Ueno et al, 2012). The FB has been also implicated in the regulation of electric- and heat-shock-induced innate avoidance and conditioned avoidance(Hu et al, 2018). The FB is composed of at least 9 layers(Hu et al, 2018, Wolff et al, 2015). Given that one can probe distinct layers of the FB, we wondered whether the entire FB or distinct FB regions are involved in dialect learning.

We first focused on the entire FB, followed by subsets of FB neurons, by selecting three GAL4 lines from the FlyLight database (Jenett et al, 2012a). The lines were chosen based on covering either the entirety (R75G12) or a subset (R38E07, R89E07, R49H02) of the FB in addition to labeling few to no other regions of the brain. Using mCD8-GFP as a reporter, immunofluorescence reveals that all FB regions are labeled by R75G12 (Figure 5 B, Supplementary Figure 27); regions 5, 8, 9 labeled by R38E07 (Figure 5 D, Supplementary Figure 28); regions 2, 8, 9 by R89E07 (Figure 5 F, Supplementary Figure 29); and regions 1, 4, 6 by R49H02 (Figure 5 H, Supplementary Figure 30). We individually blocked the output of these neurons by expression of UAS-shibire^ts^ (UAS-shi^ts^) at the restrictive temperature. We find that inactivation of the entire FB, or the sub-regions 5, 8, and 9 with the R38E07 driver line, resulted in perturbed dialect learning at the restrictive temperature, but wild-type dialect learning at the permissive temperature (Figure 5 C, E). The untrained state is wild-type at both restrictive and permissive temperatures (Supplementary Figure 31 A-D). Inactivation of regions 2, 8 and 9 with the R89E07driver line and inactivation of regions 1, 4, and 6 with the R49H02 driver line resulted in wild-type dialect learning in both the restrictive and permissive temperatures, and wild-type untrained states (Figure 5 G, I, Supplementary Figure 31 E-H). Taken together, we conclude that region 5 of the FB is necessary for dialect learning, whereas regions 1,2, 4, 6, 8, and 9 are dispensable.

**Figure 5.**
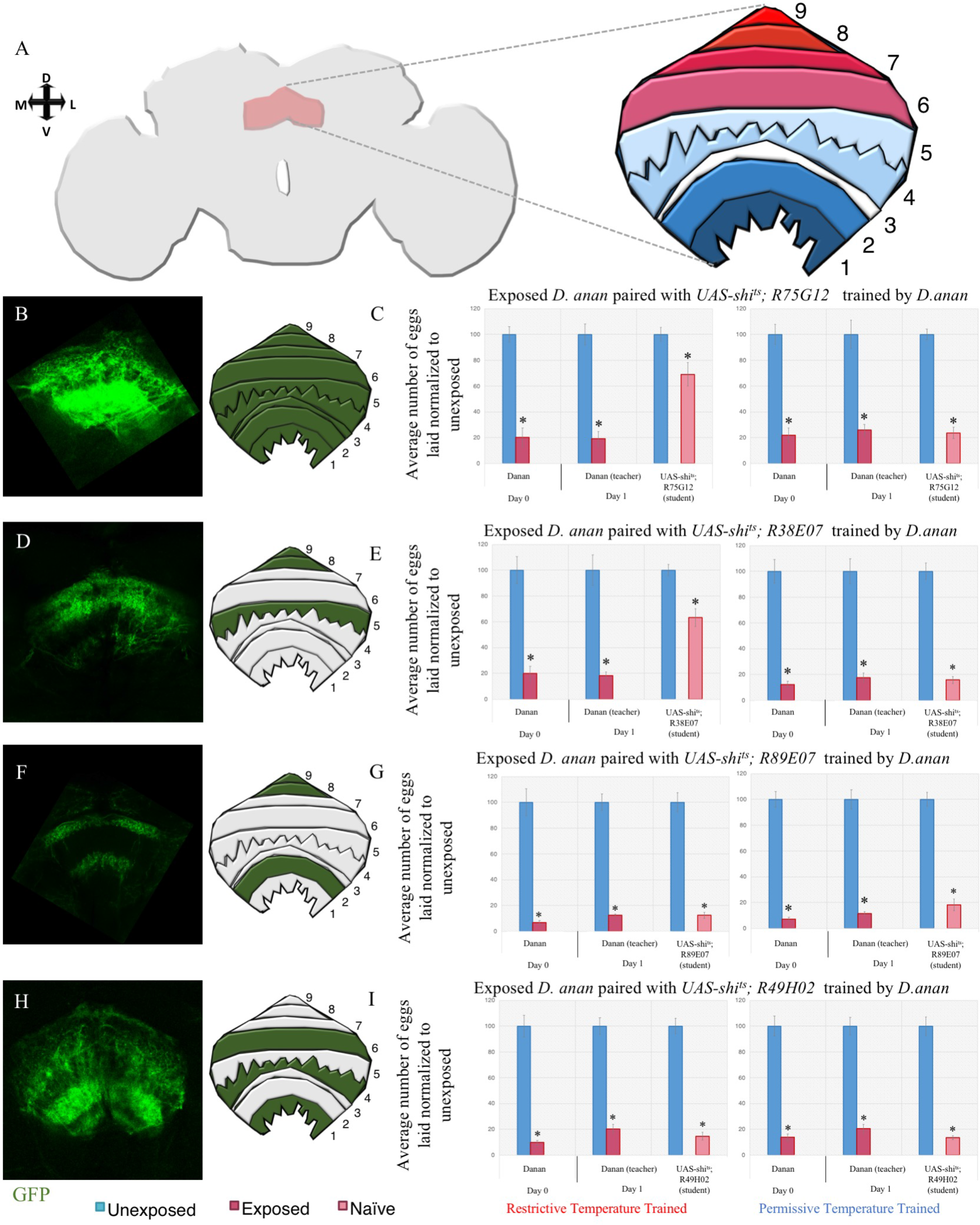
Region 5 of the fan-shaped body is necessary for dialect learning. (A) Schematic of the Drosophila fan-shaped body (FSB) with the various regions (1-9) highlighted. (B) Expression pattern and schematic of the R75G12 FSB driver shows pan-FSB marking. See also, figure S 27. Percentage of eggs laid by exposed flies normalized to eggs laid by unexposed flies is shown at the permissive (22°C) or restrictive (30°C) temperature. (C) UAS-shi^ts^ crossed to R75G12 trained by *D. ananassae* at the permissive temperature shows wild-type trained state, but at the restrictive temperature shows defective acquisition in the trained state. (D) Expression pattern and schematic of the R38E07 FSB driver shows FSB marking in regions 5,8, and 9. See also, figure S 28. (E) UAS-shi^ts^ crossed to R38E07 trained by *D. ananassae* at the permissive temperature shows wild-type trained state, but at the restrictive temperature shows defective acquisition in the trained state. (F) Expression pattern and schematic of the R89E07 FSB driver shows FSB marking in regions 2,8, and 9. See also, figure S 29. (G) UAS-shi^ts^ crossed to R89E07 trained by *D. ananassae* at the permissive and the restrictive temperatures shows wild-type trained state. (H) Expression pattern and schematic of the R49H02 FSB driver shows FSB marking in regions 2,8, and 9. See also, figure S 30. (I) UAS-shi^ts^ crossed to R49H02 trained by *D. ananassae* at the permissive and the restrictive temperatures shows wild-type trained state. All untrained states are shown in figure S 31. Error bars represent standard error (n = 12 biological replicates) (*p < 0.05).

Given that the FSB is necessary for dialect learning, specifically neurons in region 5, we wondered whether they are also sufficient. To ask this question, we expressed a temperature sensitive transient receptor potential A1 (TRPA1)(Kang et al, 2012, Lamaze et al, 2017) channel via a Gal4-mediated expression. To ascertain sufficiency of the FB, we altered the dialect learning setup—we altered the function of time whereby dialect training was performed for either 24 hours or 72 hours (Figure 6 A). When using wild-type *D. melanogaster*, we find that 24 hours is not a sufficient length of training for dialect acquisition, while 72 hours is sufficient (Figure 6 B).

**Figure 6.**
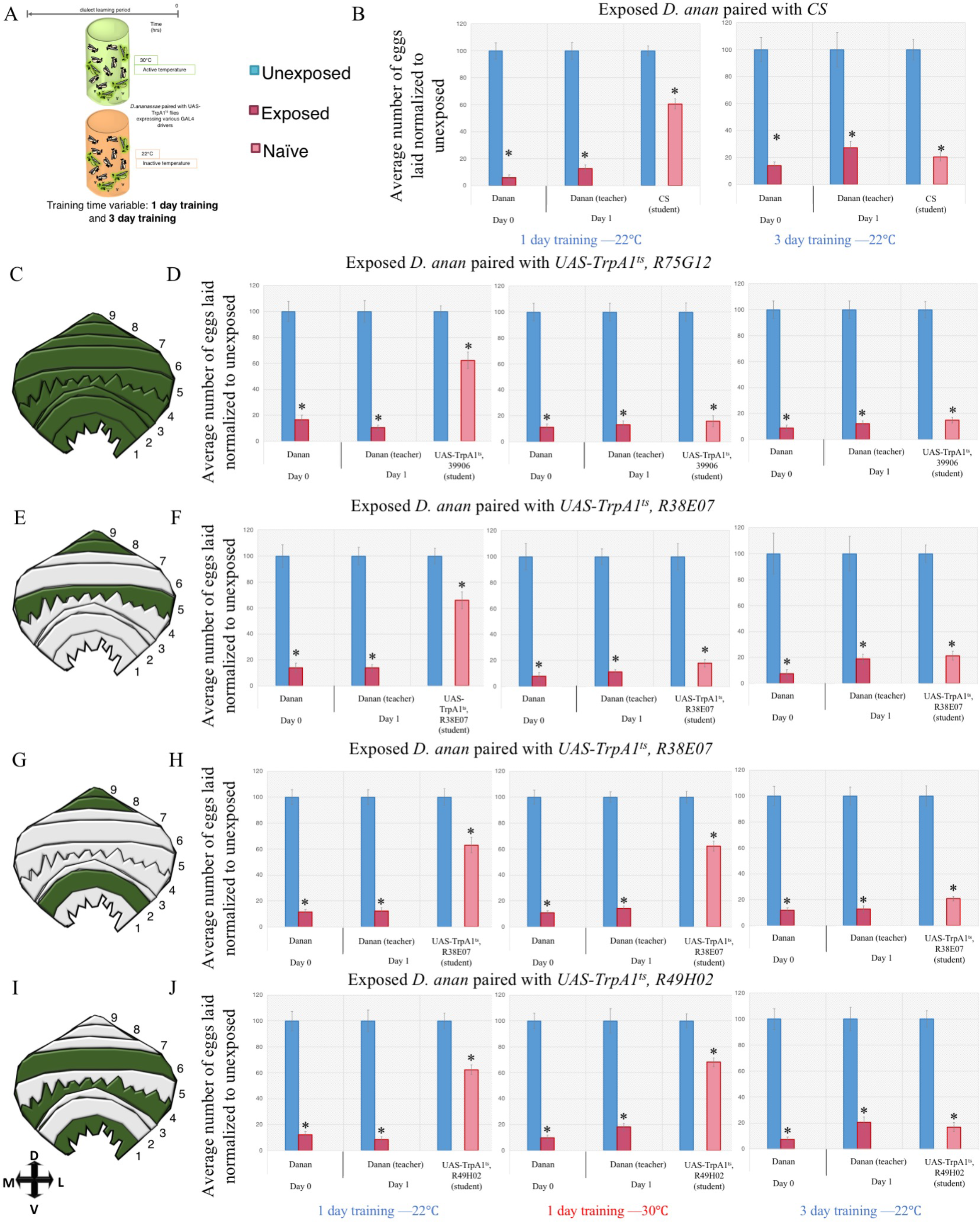
Region 5 of the fan-shaped body is sufficient for dialect learning. (A) Schematic of experimental variant introduced to standard dialect training setup—training is either 1 or 3 days at either the active (30°C) or inactive (22°C) temperature. Percentage of eggs laid by exposed flies normalized to eggs laid by unexposed flies is shown. (B) *Wild-type* flies are unable to learn the dialect from *D. ananassae* after 1 day of training but are able to learn after 3 days of training at 22°C. (C) Schematic of the R75G12 fan-shaped body (FSB) driver shows FSB marking in all regions. (D) UAS-TrpA1^ts^ crossed to R75G12 trained by *D. ananassae* at the inactive temperature shows wild-type phenotype. 1 day of training at the active temperature is sufficient to allow for dialect acquisition suggesting that this region of the FSB is sufficient to drive dialect learning. 3 days of training at the inactive temperature shows wild-type dialect acquisition. (E) Schematic of the R38E07 FSB driver shows FSB marking in regions 5,8, and 9. (F) UAS-TrpA1^ts^ crossed to R38E07 trained by *D. ananassae* at the inactive temperature shows wild-type phenotype. 1 day of training at the active temperature is sufficient to allow for dialect acquisition suggesting that this region of the FSB is sufficient to drive dialect learning. 3 days of training at the inactive temperature shows wild-type dialect acquisition. (G) Schematic of the R89E07 FSB driver shows FSB marking in regions 2,8, and 9. (H) UAS-TrpA1^ts^ crossed to R89E07 trained by *D. ananassae* at the inactive temperature shows wild-type phenotype. 1 day of training at the active temperature is not sufficient to allow for dialect acquisition suggesting that this region of the FSB is not sufficient to drive dialect learning. 3 days of training at the inactive temperature shows wild-type dialect acquisition. (I) Schematic of the R49H02 FSB driver shows FSB marking in regions 2,8, and 9. (J) UAS-TrpA1^ts^ crossed to R49H02 trained by *D. ananassae* at the permissive and the restrictive temperatures shows wild-type trained state. 3-day training period at active (30°C) is shown in figure S 32 for each genotype. Error bars represent standard error (n = 12 biological replicates) (*p < 0.05).

We utilized the temporal requirement of dialect learning to elucidate whether activation of FB neurons is sufficient to enhance dialect learning. Activation of the FB was achieved using UAS-TRPA1 for either 1 or 3 days, paired with thermally inactive/basal control, in order to see if activation can change the temporal requirement of dialect learning. We find that pan FB activation accelerates dialect learning; we observe that 1 day of training is sufficient at the activated temperature (Figure 6 D). At both the thermally inactive/basal and active temperatures for the TRPA1 protein, we observe wild-type dialect acquisition following a 3-day training regime (Figure 6 D, Supplementary Figure 32 A-B). When testing sub-regions, we find that activation of regions 5, 8, 9 produces accelerated dialect learning at the activated temperature following a 1-day regime, but activation of regions 2, 8, 9 and regions 1, 4, 6 do not produce an accelerated dialect learning output (Figure 6 E-J). All lines show wild-type dialect learning ability at both temperatures following a 3-day training regime (Figure 6 E-G, Supplementary Figure 32 C-H). Taken together, we conclude that activation of region 5 of the FB in the presence of *D. ananassae* via TRPA1 can enhance dialect learning by shortening the time required for dialect training.

## Discussion

In this study, we have elucidated the fundamental neural circuitry that governs dialect learning in Drosophila. When naïve *D. melanogaster* students are placed next to wasp-exposed *D. ananassae*, *D. melanogaster* students exhibit only partial communication ability. *D. melanogaster* can learn the dialect of *D. ananassae* after a period of cohabitation (dialect training), yielding inter-species communication enhanced to levels normally observed among conspecifics. Using the FlyLight library, in conjunction with UAS-shi^ts^ to inducibly deactivate defined subsets of neurons during dialect training, we identify 6 distinct regions of the fly brain that are involved in dialect learning: the optic lobe, mushroom body, antennal lobe, ellipsoid body, fan-shaped body, and the lateral horn (Jenett et al, 2012a). Following identification of these regions, we further elucidated a neuronal subset in three distinct regions—the antennal lobe, optic lobe, and the fan-shaped body—that are necessary for dialect learning.

The current study has identified a novel role for an olfactory channel, Or69a, where twin Ors are simultaneously expressed in the same OSN population that sense species-specific pheromones and food odorants (Lebreton et al, 2017a)(Figure 2). The OSN enervates the D glomerulus in the antennal lobe, which we hypothesize signals to the lateral horn and MB through interneurons. A cluster of lateral horn neurons, termed P1, has been identified to collect olfactory and contact chemosensory signals and subsequently elicits male courtship, and might be involved (Pavlou & Goodwin, 2013). Projection neurons from the DA1 glomerulus respond to the male produced olfactory signal, cVA (Kohl et al, 2013). The question that arises for future work is how Or69a and projection neurons from its associated D glomerulus contribute excitatory input to the sexually dimorphic circuitry of the lateral horn (Couto et al, 2005b).

Previous studies have demonstrated the role of L1 and L2 motion detecting neurons as specialized for the detection of moving light and dark edges, while L3 are involved in landmark detection (Clark et al, 2011, Joesch et al, 2010). Additional work has identified the L4 neuron as involved in the connectome of motion recognition (Silies et al, 2013). The L4 neuron receives its main input from L2, but also receives functionally significant inputs from other neurons (Silies et al, 2013). Using genetic agents restricted to L2 and L4 neurons, we observe independent silencing of either neuron results in perturbed dialect acquisition (Figure 3). When using activating agents, we find a role for L4 where activation of L4 in the absence of visual stimuli in *D. melanogaster* is sufficient for successful dialect training (Figure 4, Supplementary Figure 26). Our results argue that L4 might have a specific role in motion detecting of specific Drosophila movements. This finding is consistent with previous studies that suggest that L4 neurons may play a role in spatial summation and pooling information about contrast changes in motion detection (Rister et al, 2007b, Rivera-Alba et al, 2011, Takemura et al, 2008, Takemura et al, 2011, Takemura et al, 2017). L2 and L4 neurons make a diverse range of connections in the medulla. Knowing this, our data raise the possibility that downstream motion computations are distributed among many different neuron types. These neurons may then further converge in deeper layers of the visual system. This additional connectivity may tune neurons to particular features (De Vries & Clandinin, 2012, Egelhaaf & Borst, 1992, Hausen, 1982, Krapp et al, 1998, Mu et al, 2012). These more downstream and specialized neurons could then inform specific outputs appropriate to the visual stimulus during dialect training. It will be interesting to investigate the connectivity of L2 and L4 to these downstream neurons and extract what feature information is being detected (Supplementary Movies 1-2).

The current work has also investigated the roles of the well-defined fly brain region, the FB, and several groups of FB neurons. Genetic manipulations were specifically targeted to FB neurons by using GAL4 lines R75G12, R38E07, R89E07, and R49H02 (Jenett et al, 2012b, Li et al, 2009). The results presented demonstrate that FB neurons are capable of mediating social interactions. We identified a subgroup of large-field FB neurons contained in ventral layer 5 of the FB. Outputs of layer 5 neurons are necessary for dialect learning (Supplementary Movies 3-4). Silencing this particular layer or the entire FB demonstrates perturbed dialect acquisition (Figure 5). Activating layer 5 or the entire FB results in an alteration of the temporal dynamics of dialect learning such that flies can learn the dialect faster (Figure 6). As one of the distinct parts of the central complex in the fly brain (Hanesch et al, 1989), the FB is important for multiple functions, including locomotor control (Strauss, 2002), visual feature recognition and associated processing (Liu et al, 2006), courtship (Sakai & Kitamoto, 2006), sleep (Berry et al, 2015, Donlea et al, 2011, Ueno et al, 2012), and avoidance behaviors (Hu et al, 2018). Visual feature recognition and sleep primarily rely on dorsal FB layers, while avoidance behaviors largely rely on ventral and middle layers of FB. We identify a ventral layer of the FB involved in mediating social interactions during dialect learning. The current state of FB research suggests that different FB layers serve as hubs for distinct behavioral information.

Collectively, we present a model of dialect learning where inhibiting and activating synaptic transmission in certain brain regions results in perturbed or enhanced dialect learning, demonstrating a foundational neural circuit map of dialect learning (Figure 7, Supplementary Figure 33). Given the need for multiple sensory inputs, dialect learning is fundamentally different from the previously described teacher-student paradigms that utilizer only visual cue exchange as a means of information transfer (Kacsoh et al, 2015c). Additionally, we suggest that this study also points to previously unappreciated functions of the Drosophila MB in integrating information from olfactory and visual inputs, perhaps through the L4 motion detecting neuron in the optic lobe and the D glomerulus of the antennal lobe (Takemura et al, 2017).

**Figure 7.**
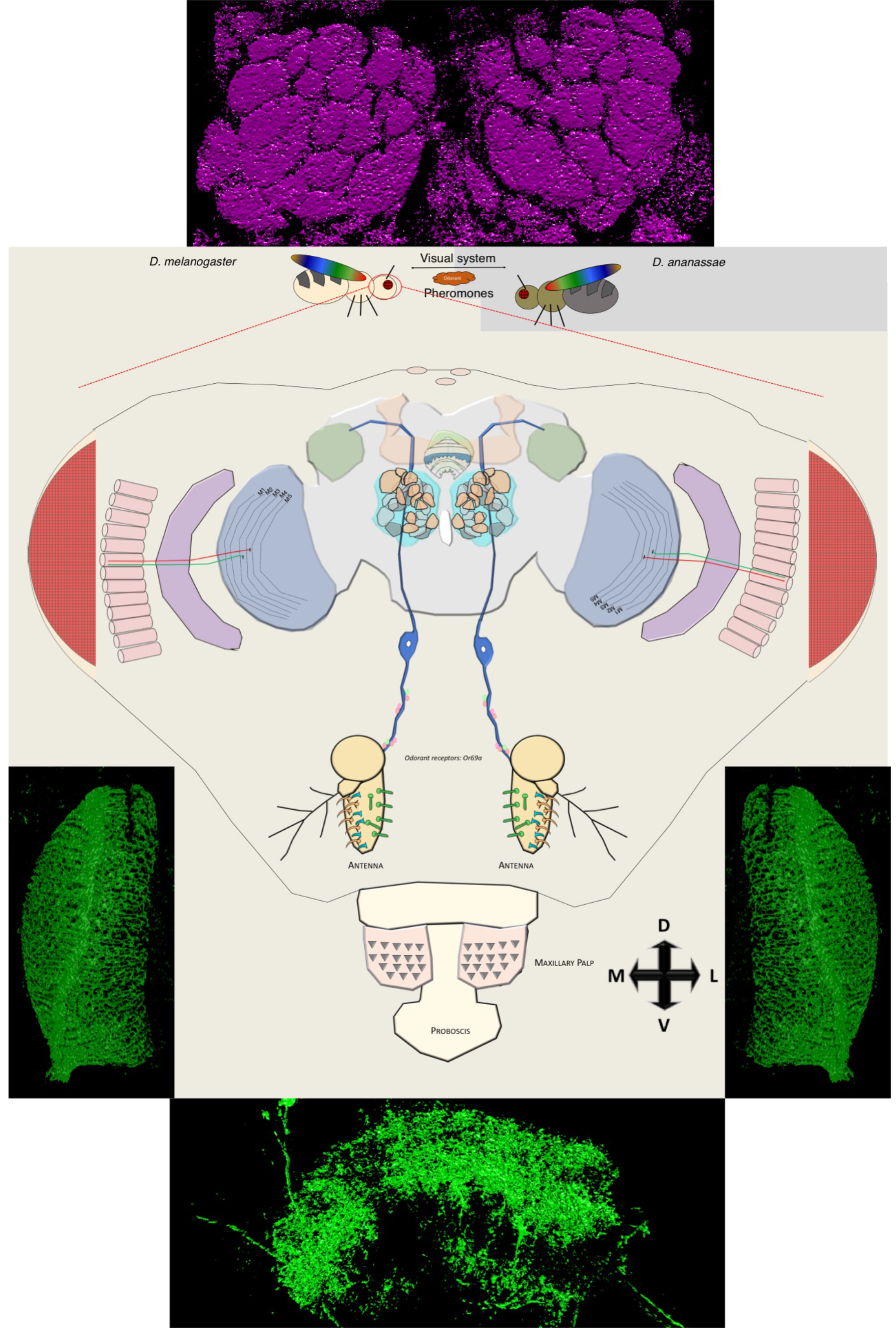
Model for the neural circuitry of dialect learning in Drosophila. Interspecies communication and dialect learning is dependent on the presence of multiple cues between *D. melanogaster* and *D. ananassae*. These cues are, in part, olfactory, visual, and ionotropic in nature. In *D. melanogaster*, we have identified multiple brain regions and neurons that contribute to the dialect learning. Regions we identified to be involved in dialect learning— the optic lobe (blue), mushroom body (orange), antennal lobe (teal), lateral horn (green), fan-shaped body (green/white), and ellipsoid body (light green). For the antennal lobe, we identify the D glomerulus, innervated by *Or69a* olfactory receptor neuron (ORN). For the fan-shaped body, we identify region 5 as necessary and sufficient for dialect learning. For the optic lobe we identify the motion detecting circuit as necessary by isolation of the L4 neurons. Flanking the model, super resolution microscopy with a 3D render is provided of the key regions of the fly brain that were investigated—region 5 of the fan-shaped body, the L4 neurons, and the antennal lobe. Super resolution images without the 3D render are located in Supplementary Figure 33.

Dialect learning between *D. melanogaster* and *D. ananassae* results in the ability of each species to more efficiently receive information about a common predator. Cognitive plasticity in conjunction with multiple circuits acts in unison to allow for dialect learning. This finding hints that adult behaviors may emerge only as a result of previous social experiences. Such experiences take place in nature, where relevant ecological pressures are ever present and multiple species co-exist and compete for resources while simultaneously cooperate against common foes. Cognitive plasticity wields a real fitness benefit to the individual and the collective, where sharing of information directly, or by coincident bystanders, could result in behavioral immunity to pan-specific threats. Our observations also raise the interesting possibility that different species share information not just about common predators, and perhaps even rudimentary social learning neuronal circuitry can be co-opted to receive and integrate useful information from a variety of close and distantly related species.

We have only just arrived at the boundary between insect behavioral neuro-genetics, evolution and ecology, and sociobiology. The current study integrates the vast tool-box and the life history of the Drosophila model in an effort to dissect the neuro-circuitry governing the observation that the direct fitness of the *individual* not being the primary determinant of fitness in a natural setting, but rather the maximization of inclusive fitness. The sociobiology trait of a group is not determined by a group member’s genetic fitness, but by the summed effects of all of the survival indices of the group. Collective wisdom in the group arises from poorly informed members, whose interactions with informed members are shaped by natural selection promoting cognitive plasticity and rudimentary sociality.

## Materials and Methods

### Insect Species/Strains

The *D. melanogaster* strain Canton-S (CS) was used as the wild-type strain and used for outcrosses. The Drosophila species *D. ananassase* and *D. virilis* were acquired from the Drosophila Species Stock Center (DSSC) at the University of California, San Diego, stock numbers 14024-0371.13 and 15010-1051.87, respectively. *L1, L2, L4^0987^*, and *splitL4* GAL4 lines were kindly provided by Marion Silies (European Science Institute, Germany). The CD8-GFP line was kindly provided by Mani Ramaswami (Trinity College, Dublin Ireland). All stocks used in experiments are listed in Table S1 with stock numbers shown (when applicable).

Flies aged 3-6 days post-eclosion on fresh Drosophila media were used in all experiments. All flies were maintained at 22ºC (the permissive temperature of experiments described later) with approximately 30-45% humidity, with a 12:12 light:dark cycle at light intensity 16_7_ with 30-55% humidity dependent on weather. Light intensity was measured using a Sekonic L-308DC light meter. The light meter utilized measures incident light and is set at shutter speed 120, sensitivity at iso8000, with a 1/10 step measurement value (f-stop). These are conditions used in previous studies (Kacsoh et al, 2018b, Kacsoh et al, 2015b, Kacsoh et al, 2015c, Kacsoh et al, 2017). All species and strains used were maintained in fly bottles (Genesse catalog number 32-130) containing 50 mL of standard Drosophila media. Bottles were supplemented with 3 Kimwipes rolled together and placed into the center of the food to promote pupation. Drosophila media was also scored to promote oviposition. Fly species stocks were kept separate to account for visual cues that could be conferred if the stocks were kept side-by-side.

We utilized the generalist Figitid larval endoparasitoid *Leptopilina heterotoma* (strain Lh14), that is known to infect a wide array of Drosophilids (Kacsoh & Schlenke, 2012, Kacsoh et al, 2014b, Schlenke et al, 2007b). *L. heterotoma* strain Lh14 originated from a single female collected in Winters, California in 2002. To propagate wasp stocks, we used adult *D. virilis*, which were placed in batches of 40 females and 15 males per vial (Genesse catalog number 32-116). The strain we used has been maintained on *D. virilis* since 2013. Adult flies are allowed to lay eggs in these standard Drosophila vials that contain 5 mL standard Drosophila media supplemented with live yeast (approximately 25 granules) for 4-6 days. Flies were then replaced by adult wasps—15 female and 6 male wasps—for infections. Infection timing gives the wasps access to the L2 stage of *D. virilis* larvae. Vials that contain wasps are supplemented with approximately 500 µL of a 50% honey/water solution that is applied to the inside of the cotton vial plugs. The honey used was organic, raw and unfiltered. Wasps aged 3-7 days post eclosion were used for all infections and experiments. Wasps were never reused for experiments, nor were they used for stock propagation if used for experiments.

### Fly Duplexes

We utilized the previously described fly duplex to examine teaching ability following dialect training (Kacsoh et al, 2018b, Kacsoh et al, 2015c). The fly duplexes were constructed (Desco, Norfolk, MA) by using three 25mm × 75mm pieces of acrylic that were adhered between two 75mm × 50mm × 3mm pieces of acrylic via clear acrylic sealant. This yields two compartments separated by one 3mm thick acrylic piece. Following sealant curing, each duplex is soaked in water and Sparkleen detergent (Fisherbrand™ catalog number 04-320-4) overnight, then soaked in distilled water overnight and finally air-dried.

For experiments using Fly Duplexes (teacher-student interaction), bead boxes (6 slot jewelers bead storage box watch part organizer sold by FindingKing) were used to accommodate 12 replicates of each treatment group. Compartments measure 32 × 114 mm with the tray in total measuring 21 × 12 × 3.5 mm. Each compartment holds 2 duplexes, and the tray holds a total of 12 duplexes (accounting for the 12 replicates). When setting up experiments, empty duplexes are placed into the bead box compartments. 50 mL standard Drosophila media in a standard Drosophila bottle (Genesse catalog number 32-130) is heated (Panasonic brand microwave) for 54 seconds. This heated media is then allowed to cool for 2 minutes on ice before being dispensed into the duplexes. Each duplex unit is filled with approximately 5 mL of the media and further allowed to cool until solidification. Food was scored to promote oviposition. We find that *D. ananassae* oviposition rate is extremely low in control conditions at 22ºC without scoring. The open end of the Fly Duplex is plugged with a cotton plug (Genesse catalog number 51-102B) to prevent insect escape. 10 female flies and 2 male flies are placed into one chamber of the Fly Duplex in the control, while 20 female Lh14 wasps are placed into the neighboring chamber in the experimental setting for 24 hours. After the 24-hour exposure, flies and wasps were removed by anesthetizing flies and wasps in the Fly Duplexes. Control flies undergo the same anesthetization protocol. Wasps are removed and replaced with 10 female and two male “student” flies. All flies are placed into new, clean duplexes for the second 24-hour period, containing 5 mL Drosophila media in a new bead box, prepared in the same manner as described for day 0 (wasp exposure period). Plugs used to keep insects in the duplexes are replaced every 24 hours to prevent odorant deposition on plugs that could influence behavior. The oviposition bead box from each treatment is also replaced 24 hours after the start of the experiment, and the second bead box is removed 48 hours after the start of the experiment. Fly egg counts from each bead box were made at the 0-24 and 24-48-hour time points in a blinded manner, where coding/decoding and counting are performed by two separate individuals. Used bead boxes and fly duplexes are soaked overnight in a 4% Sparkleen detergent solution (Fisherbrand™ catalog number 04-320-4), rinsed with distilled water, and allowed to air dry before subsequent use.

All experimental treatments as described above were run at 22ºC with a 12:12 light:dark cycle at light intensity 16_7_, using twelve replicates at 40% humidity unless otherwise noted. Fly duplexes and bead boxes soaked with distilled water mixed with Sparkleen after every use for 4 hours at minimum and subsequently rinsed with distilled water and air-dried. All egg plates were coded and scoring was blind as the individual counting eggs was not aware of treatments or genotypes. Genotypes were numerically coded and not revealed until the conclusion of the experiment.

### Dialect Exposure

Drosophila species were cohabitated during dialect training in standard Drosophila bottles (Genesee catalog number 32-130) containing 50 mL standard Drosophila media. Three Kimwipes were rolled together and placed into the center of the food. Three bottles were prepared per treatment. *D. melanogaster* and *D. ananassae* are incubated in each bottle with 100 female and 20 males of each species per bottle. Every two days, flies are placed into new bottles that prepared in the identical manner described above. Flies were cohabitation for approximately 168 hours (7 days), except where otherwise noted. This cohabitation takes place at either 30ºC or 22ºC depending on treatment parameters. Following cohabitation, flies are anesthetized and the two species are separated. *D. melanogaster* are then used as students to wasp or mock exposed *D. ananassae* teachers. This portion of the experiment always took place at 22ºC. For example, we cohabitated *D. melanogaster* and *D. ananassae* for one week at either 22ºC or 30ºC. Following the weeklong cohabitation, we separated the dialect-trained flies. Trained *D. melanogaster* were placed in duplexes next to *D. ananassae* either mock or wasp exposed at 22ºC.

For the one-day and three-day cohabitation experiments, batches of 3 bottles with 100 female and 20 males of each species were placed at 22ºC or 30ºC for 24 hours or 72 hours. Following the 24- or 72-hour cohabitation, flies were anesthetized and the two species were separated. *D. melanogaster* are then used as students to wasp or mock exposed *D. ananassae* teachers.

All experimental, dialect training treatments were run either at 30ºC or 22ºC (as indicated in figures) with a 12:12 light:dark cycle at light intensity 16_7_, using twelve replicates at 40% humidity unless otherwise noted. Light intensity was measured using a Sekonic L-308DC light meter. The light meter measures incident light and was set at shutter speed 120, sensitivity at iso8000, with a 1/10 step measurement value (f-stop). Additionally, all treatments were coded and scoring was blind as the individual counting eggs was not aware of treatments or genotypes. Coding and decoding was performed by a single individual, while counting was performed by another individual. All raw egg counts and corresponding p-values are provided in supplementary file 1.

### Cross set-up

For experiments utilizing either UAS-shi^ts^ or UAS-TRPA1^TS^, male flies containing tissue specific drivers were crossed to virgin female flies containing the UAS construct of interest. 20 female UAS virgin flies were mated to 10 males containing the selected hairpin. Crosses were performed in standard Drosophila fly bottles (Genesse catalog number 32-130) containing 50 mL of standard Drosophila media, supplemented with 10 granules of activated yeast. Crosses were kept at approximately 20ºC, with a 12:12 light:dark cycle at light intensity 16_7_ with 30-45% humidity dependent on weather. Crosses were moved to new bottles every two days until oviposition rates declined, at which point the adults were disposed of. F1’s were harvested upon eclosion and stored in standard Drosophila vials (Genesse catalog number 32-116) containing 5 mL Drosophila media with approximately 20 female and 5 male flies until utilized for experimentation.

### Immunofluorescence

Drosophila brains containing genetically distinct GAL4 constructs driving a mCD8-GFP were dissected in a manner previously described(Kacsoh et al, 2015b, Kacsoh et al, 2015c). Flies expressing GAL4 in a tissue specific manner driving mCD8-GFP were placed in batches into standard vials (Genesee catalog number 32-116) of 20 females, 2 males. Three vials were prepared to produce three replicates to account for batch effects of expression. We observed no expression batch effects. Brains that were prepared for immunofluorescence were fixed in 4% methanol-free formaldehyde in PBS with 0.001% Triton-X for approximately five minutes. The samples were then washed in PBS with 0.1% Triton-X and placed into block solution for 2 hours (0.001% Triton-X solution 5% normal goat serum). Samples were then placed into a 1:10 dilution of nc82 antibody (Developmental Studies at Hybridoma Bank, University of Iowa, Registry ID AB 2314866) to 0.001% Triton-X overnight at 4ºC. Following the overnight primary stain, samples were washed in PBS with 0.1% Triton-X three times. Secondary Cy3 antibody (Jackson Immunoresearch) was then placed on samples in a 1:200 ratio in 0.001% Triton-X. Following secondary staining, samples were washed in PBS with 0.1% Triton-X three times. This was followed by a 10-minute nuclear stain with 4′, 6-diamidino-2-phenylindole (DAPI) with three additional washes in PBS with 0.1% Triton-X before being mounted in vecta shield (vector laboratories catalog number: H-1000). Samples were then immediately imaged.

### Imaging

A Zeiss LSM 880 with Airyscan Confocal microscope was used for brain imaging for all samples with the exception of Supplementary figure 8, where a Leica SP8 confocal microscope was used with Leica lighting image deconvolution. Image averaging of 16x during image capture was used for all images. Whole head images of antennal GFP expression were taken using Nikon E800 Epifluorescence microscope with Olympus DP software. For Supplementary Movies 1-4 and Supplementary Figure 30, a spectral confocal/super-resolution microscope LSM 880 with Airyscan was used.

### Statistical analysis

Statistical tests on exposed v unexposed/teacher v student interactions were performed in Microsoft Excel. Welch’s two-tailed t-tests were performed for data. P-values reported were calculated for comparisons between paired treatment-group and unexposed and are included in supplemental file 1.

## Acknowledgements

We thank Mani Ramaswami, Marion Silies, FlyBase, the Vienna Drosophila Resource Center, and the Bloomington Drosophila Stock Center, for stocks. We acknowledge grants from Geisel School of Medicine at Dartmouth, the National Institute of Health Pioneer grant 1DP1MH110234 (GB), and the Defense Advanced Research Projects Agency grant HR0011-15-1-0002 (GB).

## SUPPLEMENTARY FIGURE LEGENDS

**Supplementary Figure 1.**
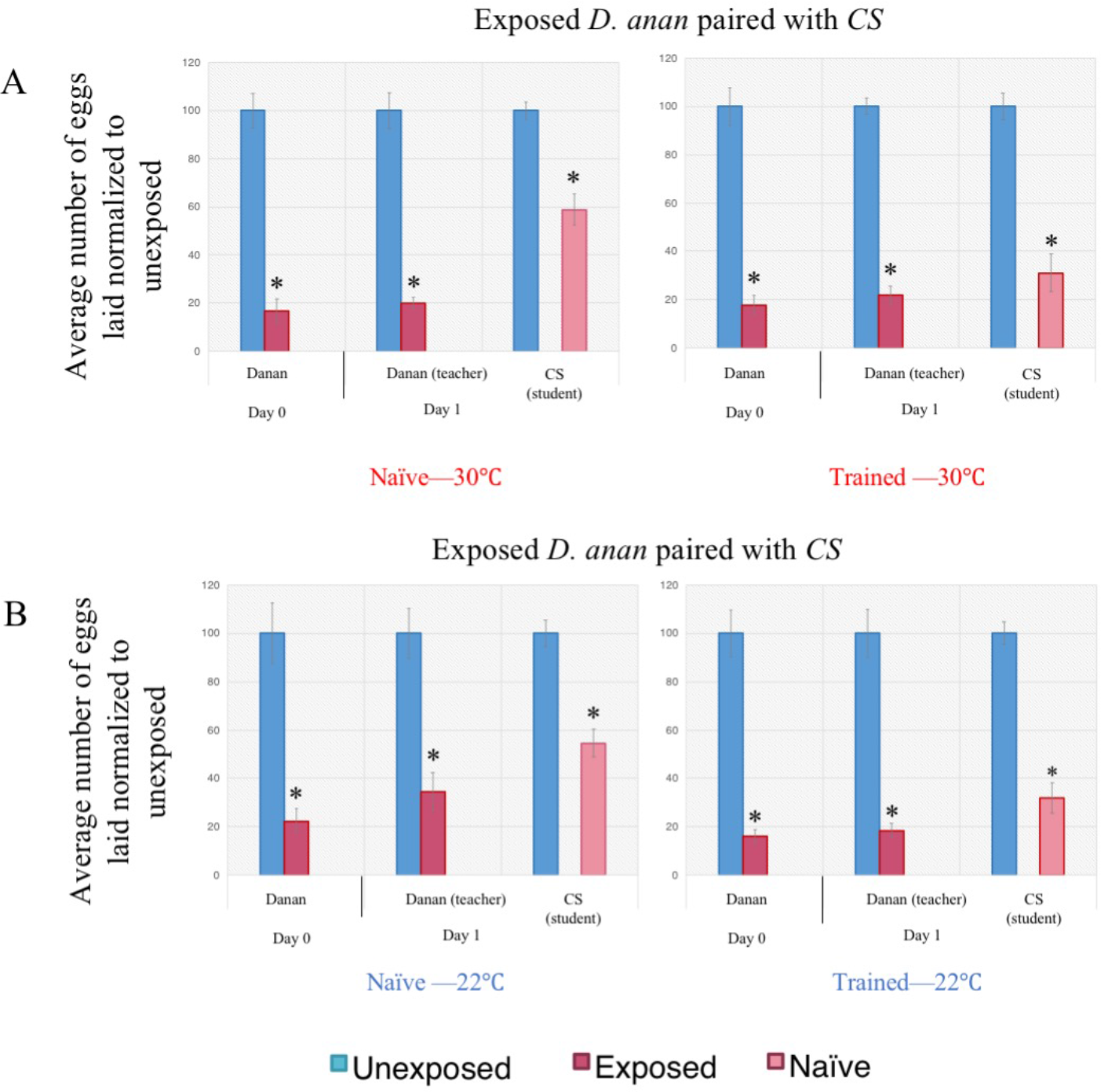
Wild-type *D. melanogaster* can be trained by *D. ananassae* at either 22°C or 30°C. Dialect learning is performed at either the permissive (22°C) or restrictive (30°C) temperature, while the wasp exposure and social learning period is performed exclusively at the permissive temperature. Percentage of eggs laid by exposed flies normalized to eggs laid by unexposed flies is shown. *Canton S* naïve and trained states by *D. ananassae* at either 30°C (A) or 22°C (B) show wild-type dialect acquisition. Error bars represent standard error (n = 12 biological replicates) (*p < 0.05).

**Supplementary Figure 2.**
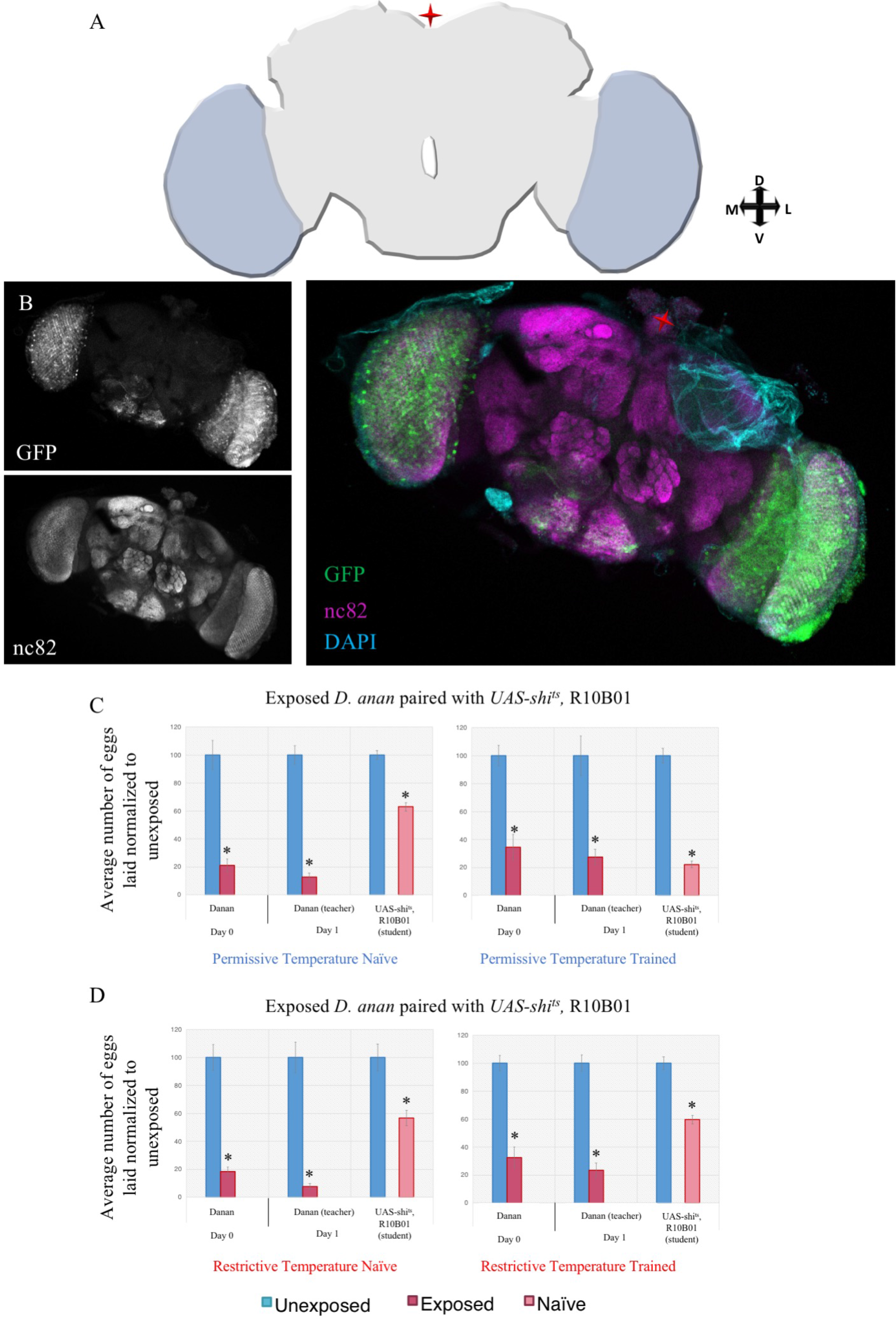
The optic lobe is required for dialect learning shown with R10B01 driver. (A) Cartoon schematic of the region of interest. (B) Confocal image of adult brain where R10B01-GAL4 is driving UAS-CD8-GFP, stained with nc82 (magenta) and DAPI (teal). Fly light intensity/distribution score is 5/2 and was the only region of the brain identified. Percentage of eggs laid by exposed flies normalized to eggs laid by unexposed flies is shown. UAS-shi^ts^ crossed to R10B01-GAL4 trained by *D. ananassae* at the permissive temperature shows wild-type trained state (C), but at the restrictive temperature shows defective acquisition in the trained state (D). Error bars represent standard error (n = 12 biological replicates) (*p < 0.05).

**Supplementary Figure 3.**
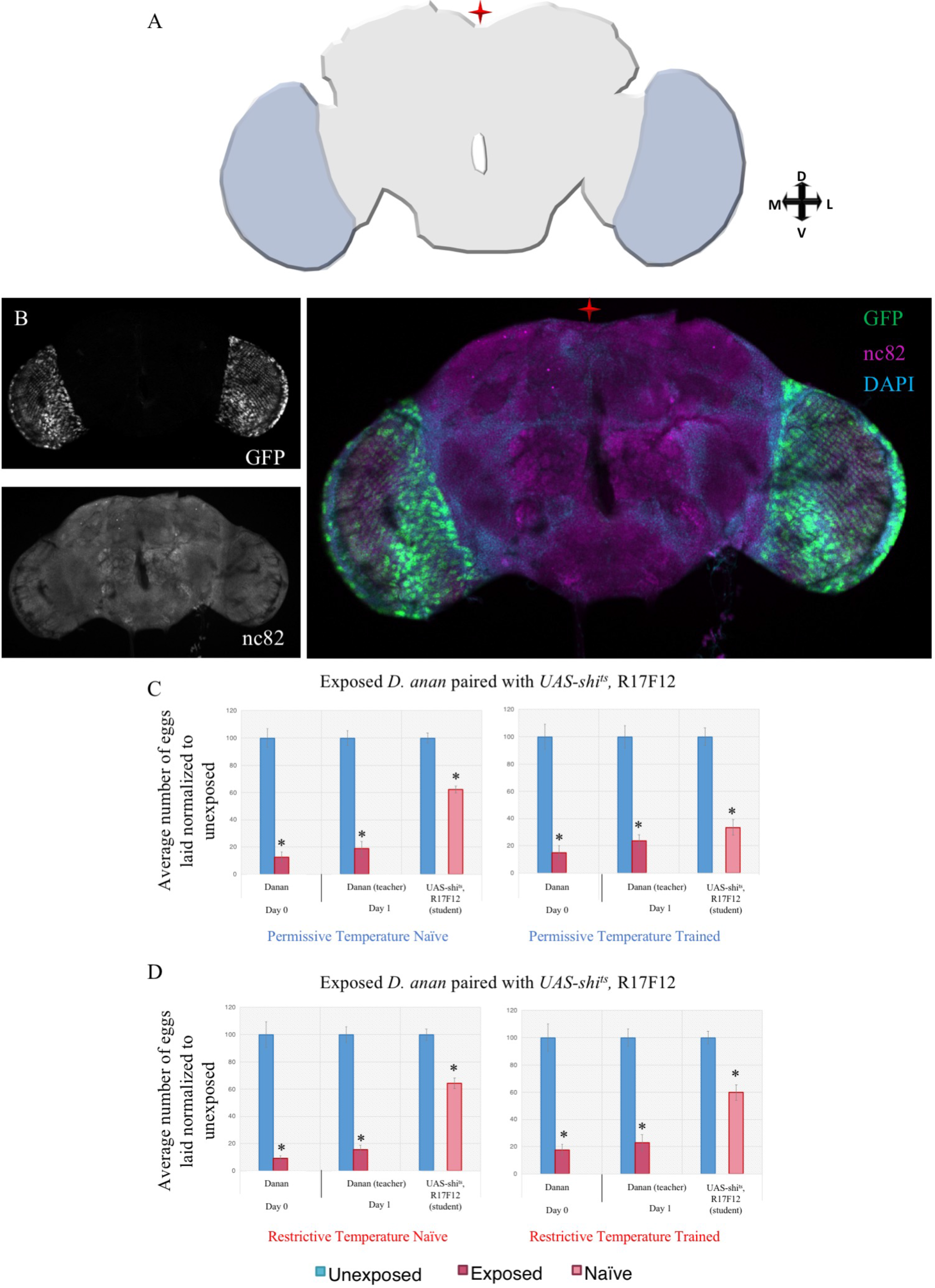
The optic lobe is required for dialect learning shown with R17F12 driver. (A) Cartoon schematic of the region of interest. (B) Confocal image of adult brain where R17F12-GAL4 is driving UAS-CD8-GFP, stained with nc82 (magenta) and DAPI (teal). Fly light intensity/distribution score is 5/2 and was the only region of the brain identified. Percentage of eggs laid by exposed flies normalized to eggs laid by unexposed flies is shown. UAS-shi^ts^ crossed to R17F12-GAL4 trained by *D. ananassae* at the permissive temperature shows wild-type trained state (C), but at the restrictive temperature shows defective acquisition in the trained state (D). Error bars represent standard error (n = 12 biological replicates) (*p < 0.05).

**Supplementary Figure 4.**
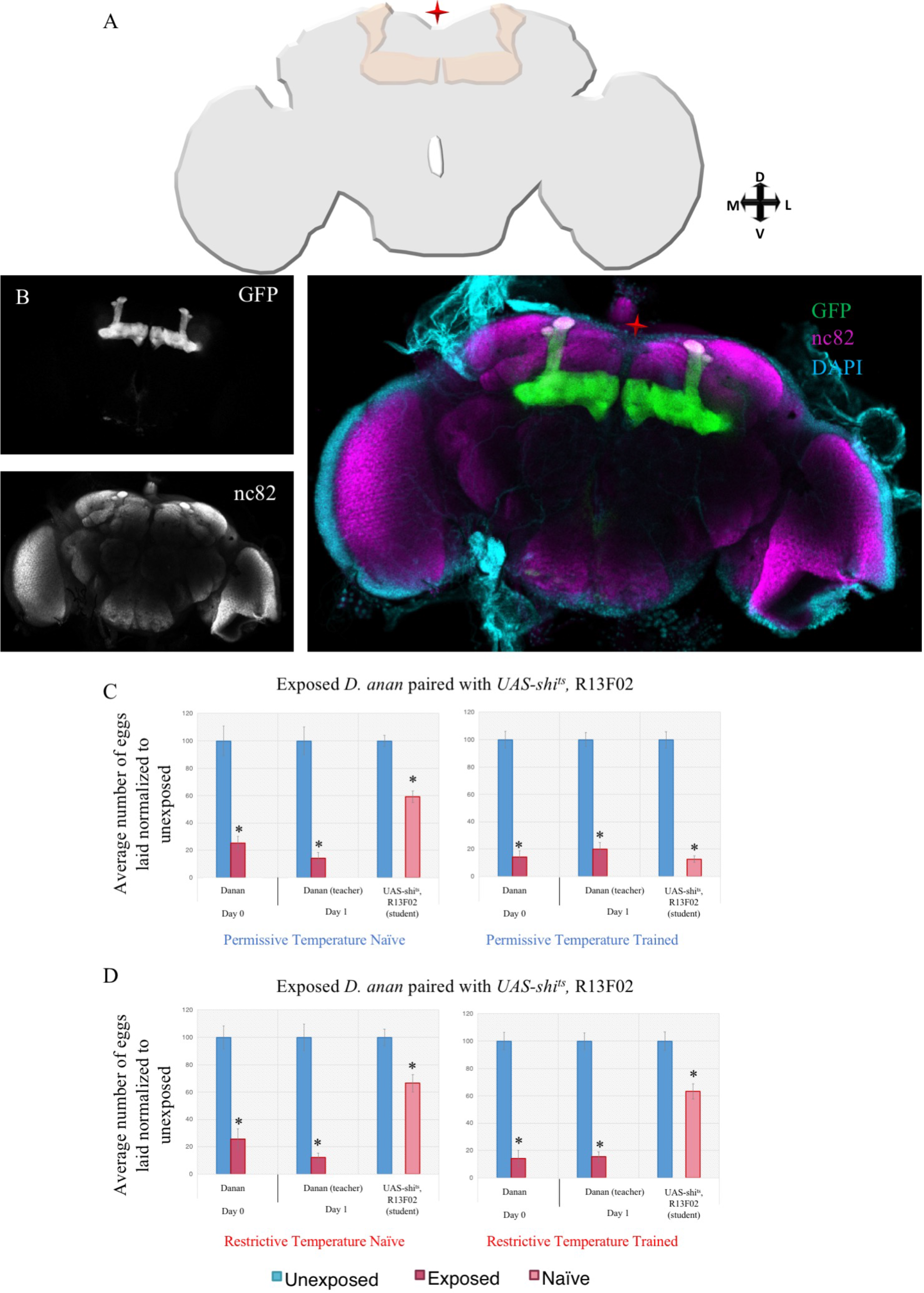
The mushroom body is required for dialect learning shown with R13F02 driver. (A) Cartoon schematic of the region of interest. (B) Confocal image of adult brain where R13F02-GAL4 is driving UAS-CD8-GFP, stained with nc82 (magenta) and DAPI (teal). Fly light intensity/distribution score is 5/4 and was the only region of the brain identified. Percentage of eggs laid by exposed flies normalized to eggs laid by unexposed flies is shown. UAS-shi^ts^ crossed to R13F02-GAL4 trained by *D. ananassae* at the permissive temperature shows wild-type trained state (C), but at the restrictive temperature shows defective acquisition in the trained state (D). Error bars represent standard error (n = 12 biological replicates) (*p < 0.05).

**Supplementary Figure 5.**
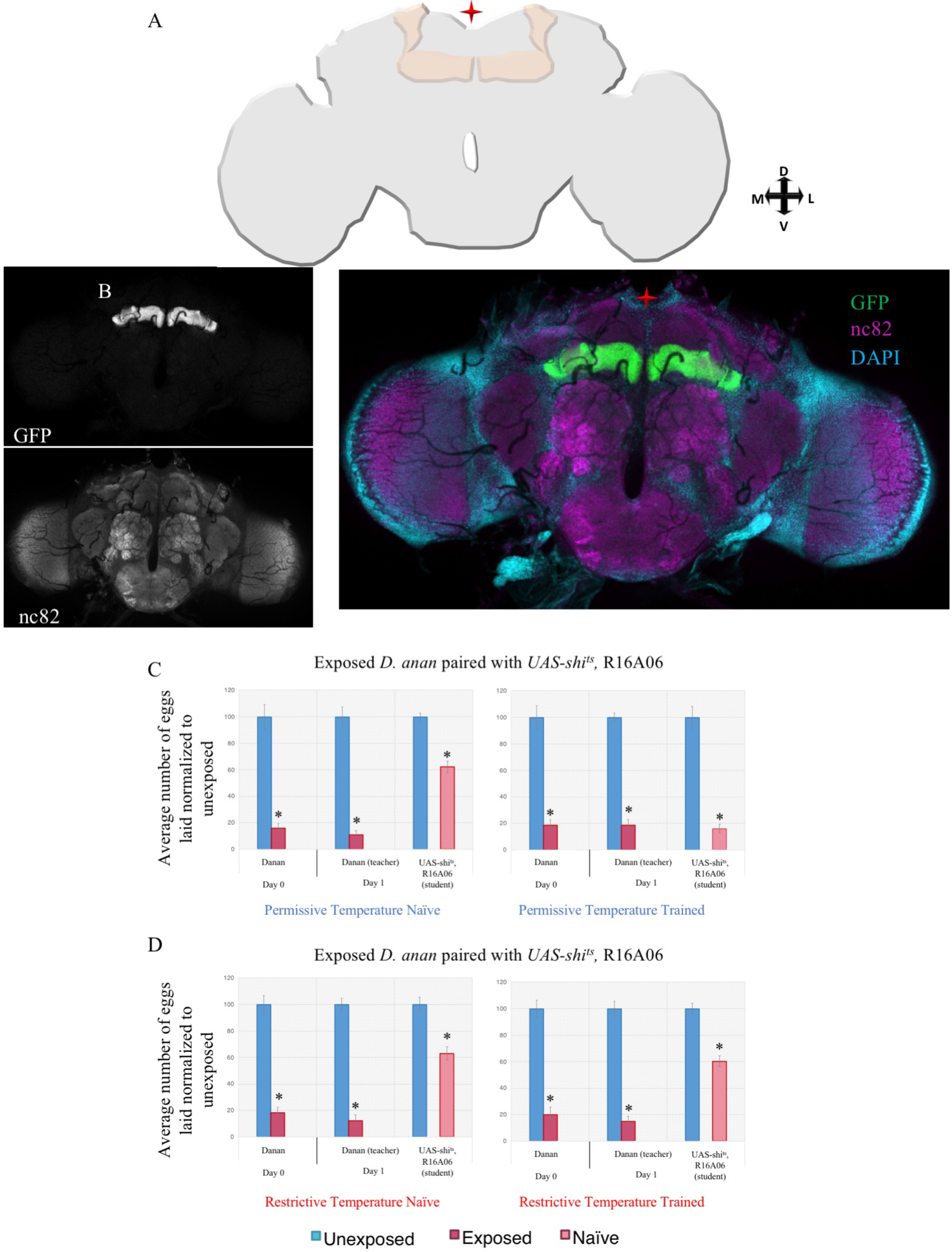
The mushroom body is required for dialect learning shown with R16A06 driver. (A) Cartoon schematic of the region of interest. (B) Confocal image of adult brain where R16A06-GAL4 is driving UAS-CD8-GFP, stained with nc82 (magenta) and DAPI (teal). Percentage of eggs laid by exposed flies normalized to eggs laid by unexposed flies is shown. UAS-shi^ts^ crossed to R16A06-GAL4 trained by *D. ananassae* at the permissive temperature shows wild-type trained state (C), but at the restrictive temperature shows defective acquisition in the trained state (D). Error bars represent standard error (n = 12 biological replicates) (*p < 0.05).

**Supplementary Figure 6.**
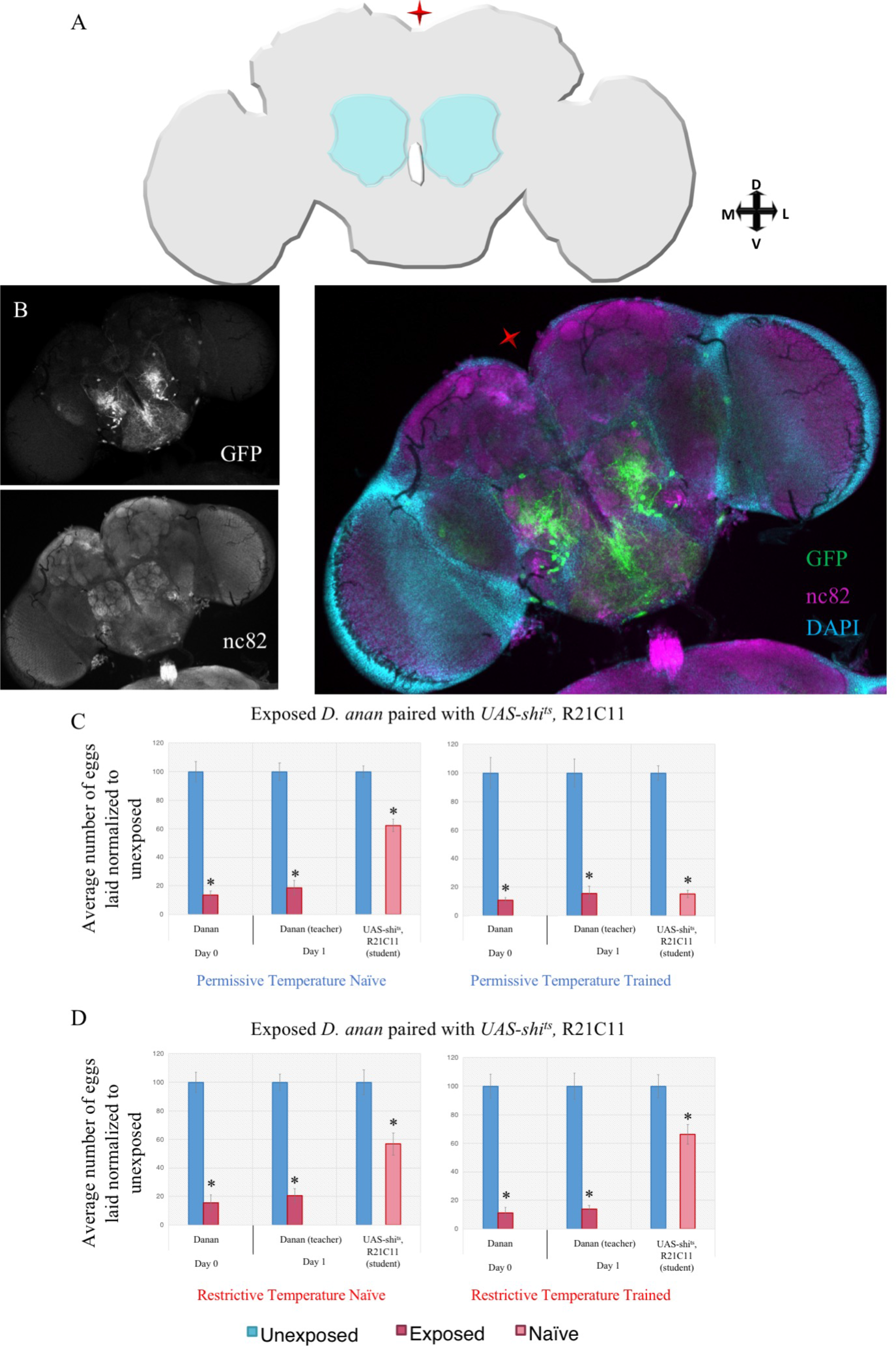
The antennal lobe is required for dialect learning shown with R21C11 driver. (A) Cartoon schematic of the region of interest. (B) Confocal image of adult brain where R21C11-GAL4 is driving UAS-CD8-GFP, stained with nc82 (magenta) and DAPI (teal). Fly light intensity/distribution score is 4/4 and was the only region of the brain identified. Percentage of eggs laid by exposed flies normalized to eggs laid by unexposed flies is shown. UAS-shi^ts^ crossed to R21C11-GAL4 trained by *D. ananassae* at the permissive temperature shows wild-type trained state (C), but at the restrictive temperature shows defective acquisition in the trained state (D). Error bars represent standard error (n = 12 biological replicates) (*p < 0.05).

**Supplementary Figure 7.**
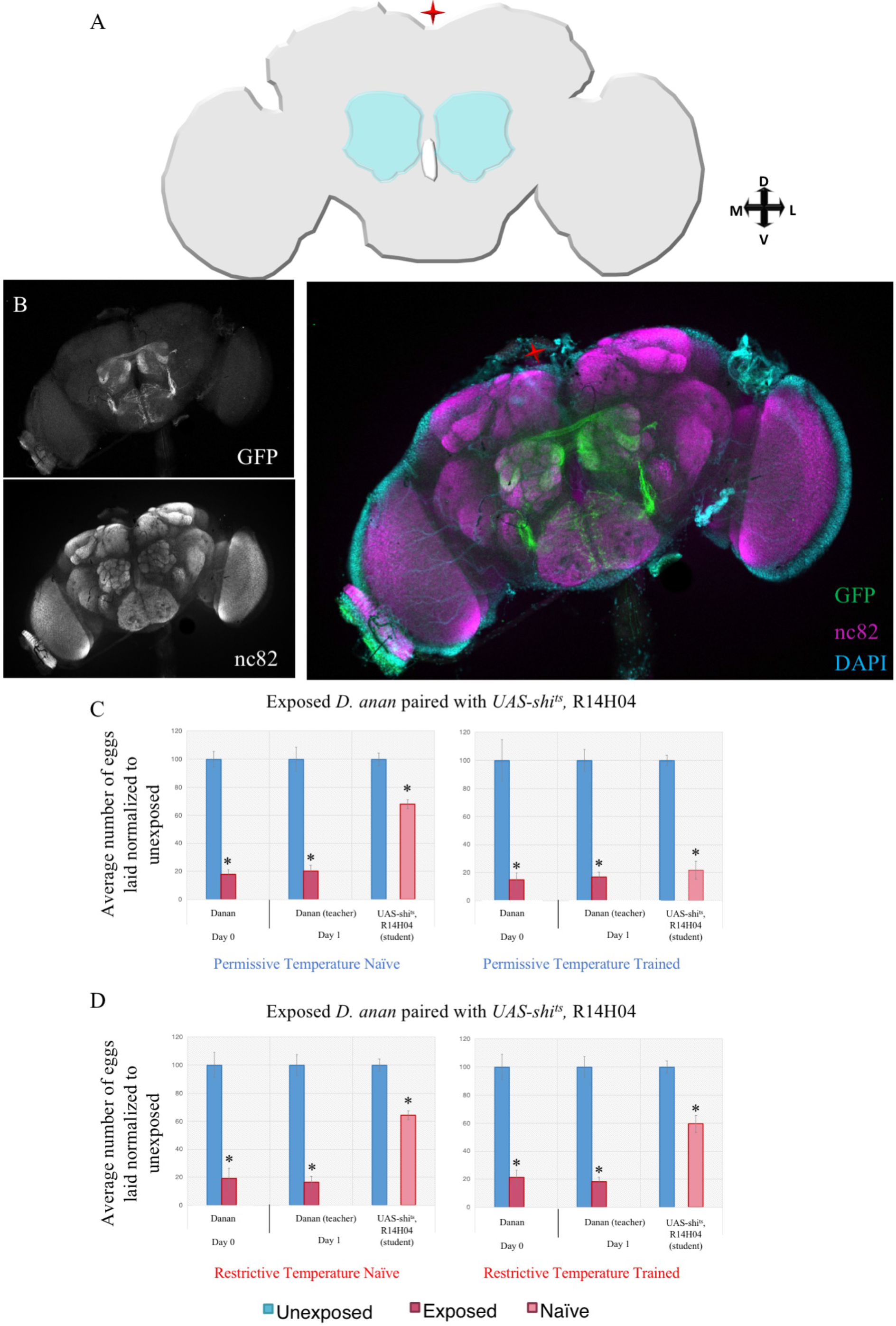
The antennal lobe is required for dialect learning shown with R14H04 driver. (A) Cartoon schematic of the region of interest. (B) Confocal image of adult brain where R14H04-GAL4 is driving UAS-CD8-GFP, stained with nc82 (magenta) and DAPI (teal). Fly light intensity/distribution score is 5/5 and was the only region of the brain identified. Percentage of eggs laid by exposed flies normalized to eggs laid by unexposed flies is shown. UAS-shi^ts^ crossed to R14H04-GAL4 trained by *D. ananassae* at the permissive temperature shows wild-type trained state (C), but at the restrictive temperature shows defective acquisition in the trained state (D). Error bars represent standard error (n = 12 biological replicates) (*p < 0.05).

**Supplementary Figure 8.**
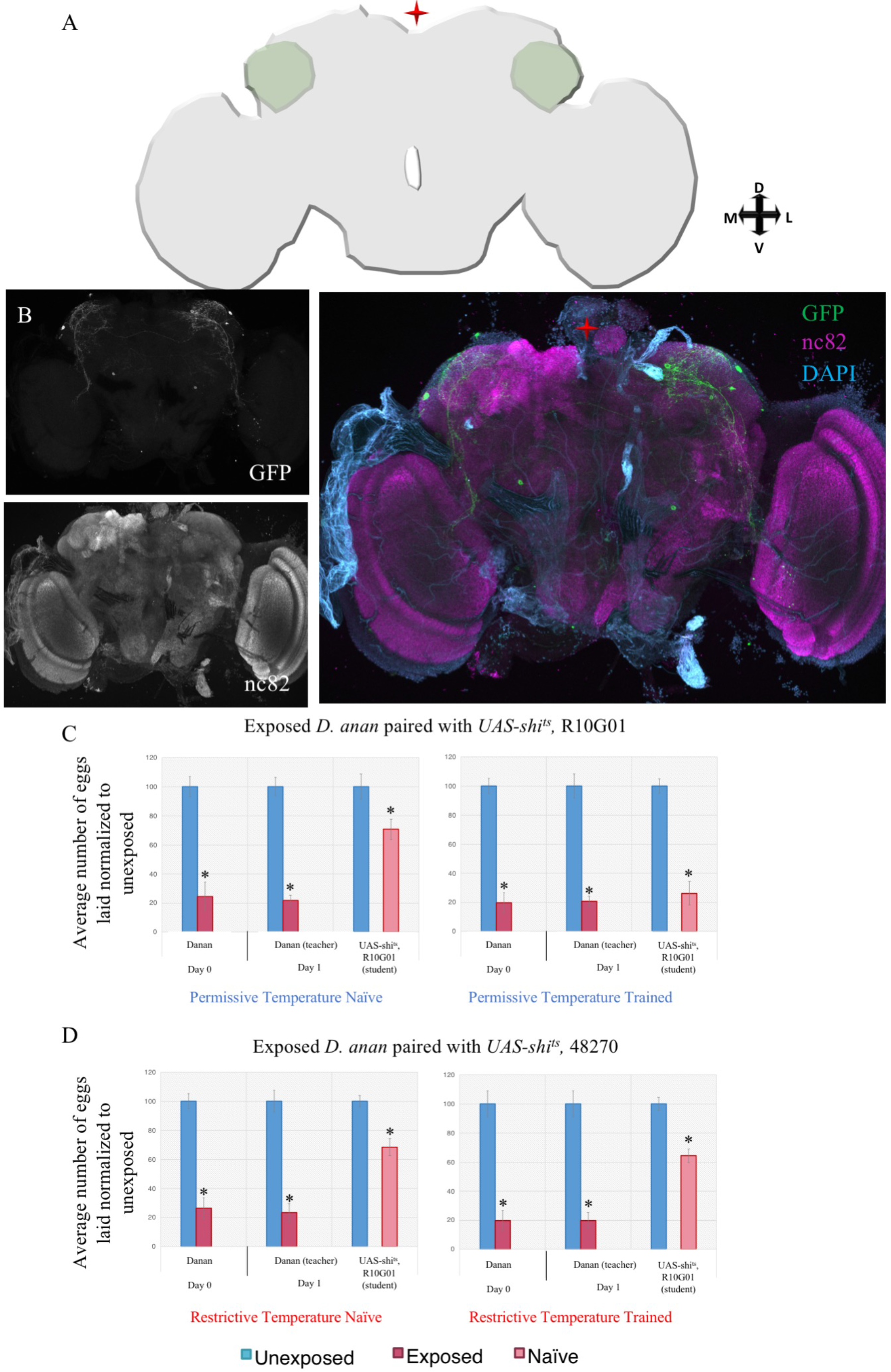
The lateral horn is required for dialect learning shown with R10G01 driver. (A) Cartoon schematic of the region of interest. (B) Confocal image of adult brain where R10G01-GAL4 is driving UAS-CD8-GFP, stained with nc82 (magenta) and DAPI (teal). Fly light intensity/distribution score is 2/2 and was the only region of the brain identified. Percentage of eggs laid by exposed flies normalized to eggs laid by unexposed flies is shown. UAS-shi^ts^ crossed to R10G01-GAL4 trained by *D. ananassae* at the permissive temperature shows wild-type trained state (C), but at the restrictive temperature shows defective acquisition in the trained state (D). Error bars represent standard error (n = 12 biological replicates) (*p < 0.05).

**Supplementary Figure 9.**
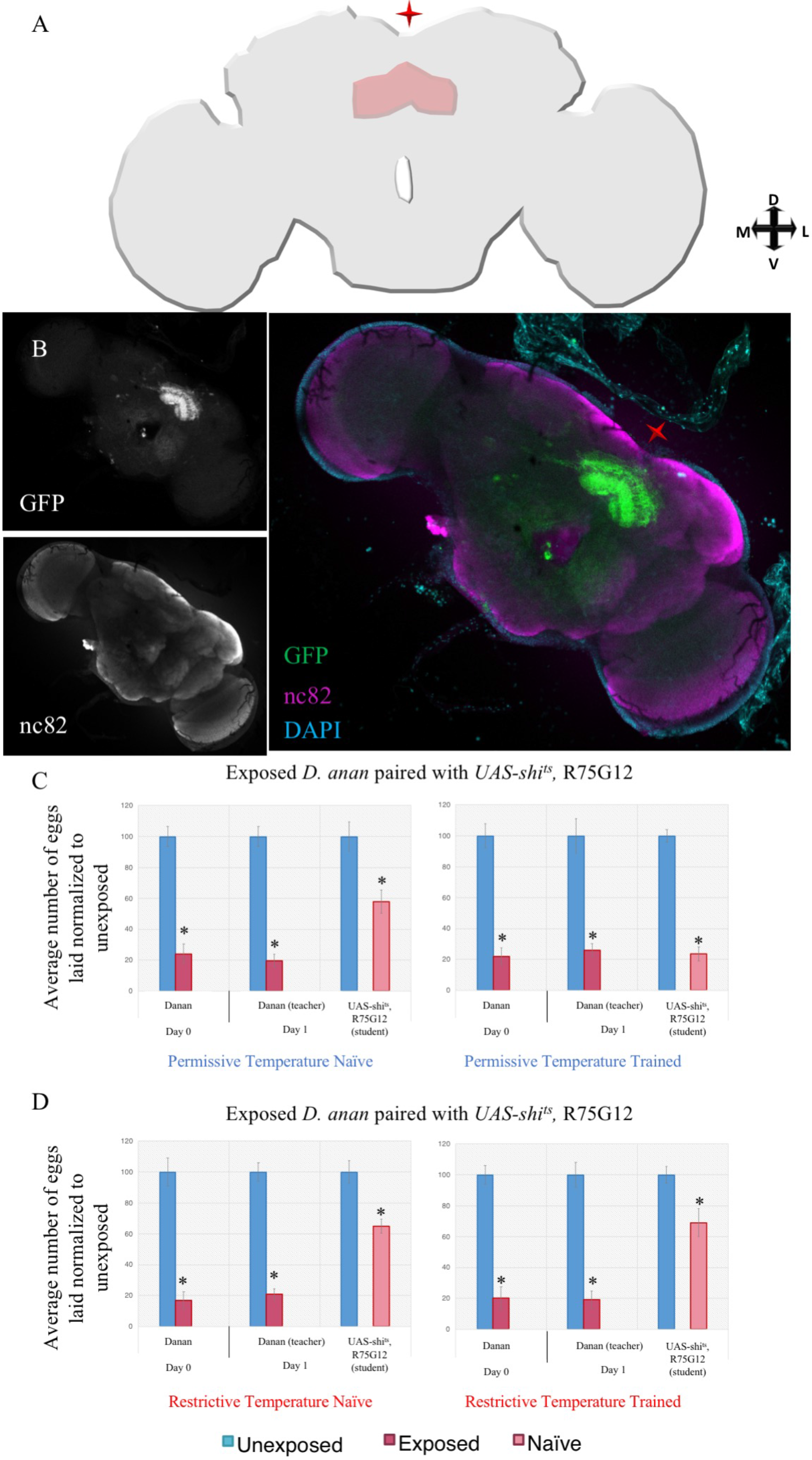
The fan-shaped body is required for dialect learning shown with R75G12 driver. (A) Cartoon schematic of the region of interest. (B) Confocal image of adult brain where R75G12-GAL4 is driving UAS-CD8-GFP, stained with nc82 (magenta) and DAPI (teal). Fly light intensity/distribution score is 5/3 and was the only region of the brain identified. Percentage of eggs laid by exposed flies normalized to eggs laid by unexposed flies is shown. UAS-shi^ts^ crossed to R75G12-GAL4 trained by *D. ananassae* at the permissive temperature shows wild-type trained state (C), but at the restrictive temperature shows defective acquisition in the trained state (D). Error bars represent standard error (n = 12 biological replicates) (*p < 0.05).

**Supplementary Figure 10.**
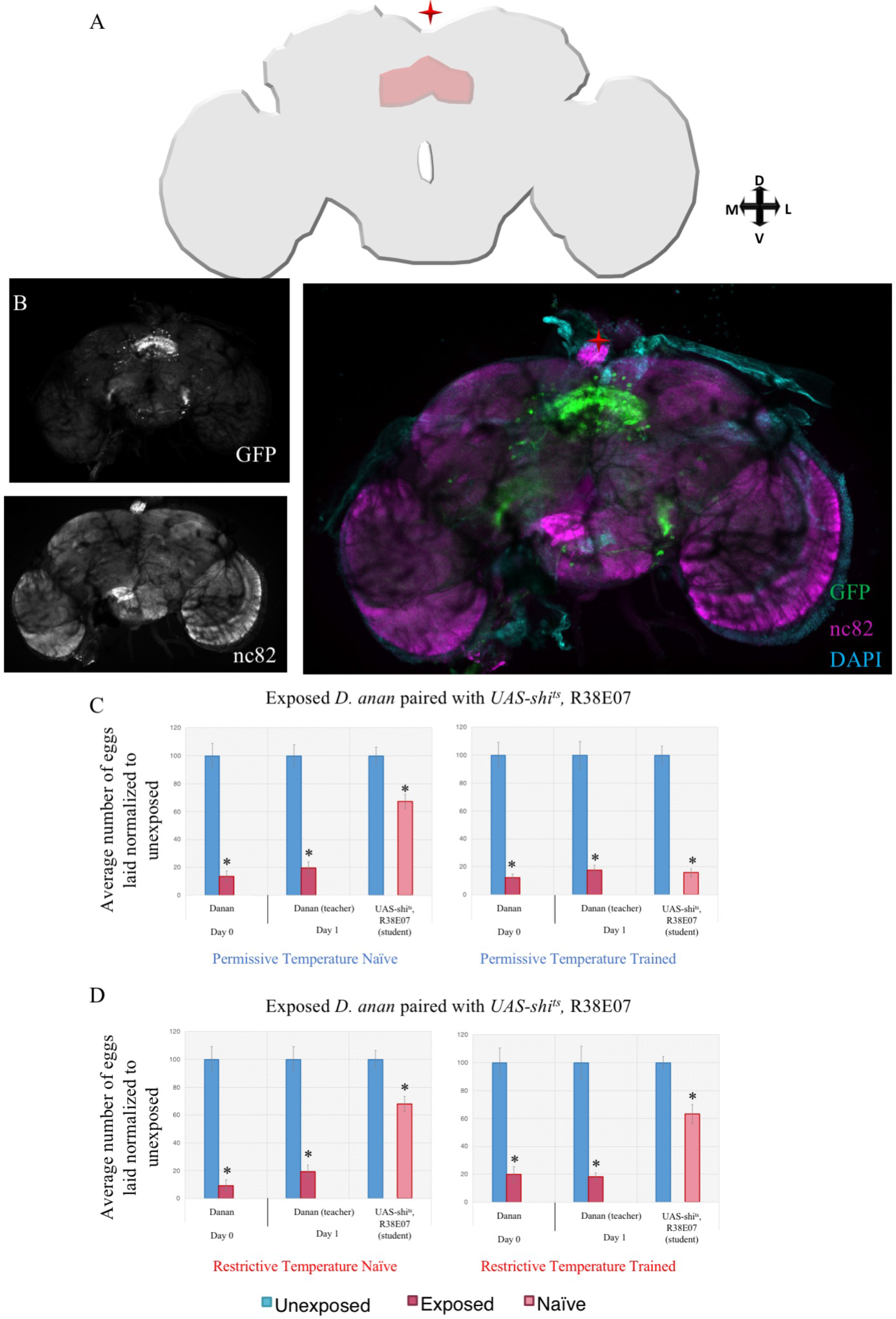
The fan-shaped body is required for dialect learning shown with R38E07 driver. (A) Cartoon schematic of the region of interest. (B) Confocal image of adult brain where R38E07-GAL4 is driving UAS-CD8-GFP, stained with nc82 (magenta) and DAPI (teal). Fly light intensity/distribution score is 3/3 and was the only region of the brain identified. Percentage of eggs laid by exposed flies normalized to eggs laid by unexposed flies is shown. UAS-shi^ts^ crossed to R38E07-GAL4 trained by *D. ananassae* at the permissive temperature shows wild-type trained state (C), but at the restrictive temperature shows defective acquisition in the trained state (D). Error bars represent standard error (n = 12 biological replicates) (*p < 0.05).

**Supplementary Figure 11.**
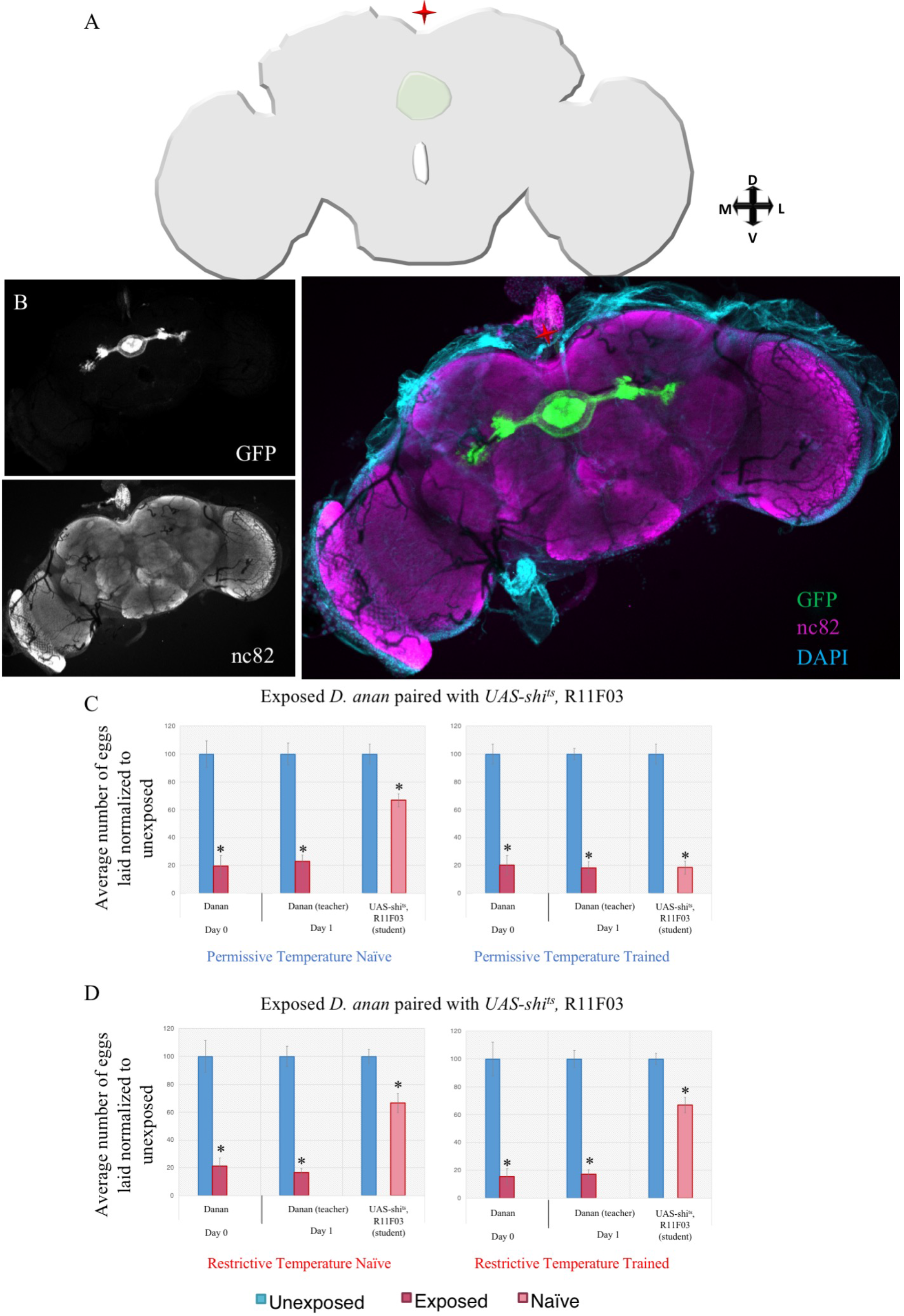
The ellipsoid body is required for dialect learning shown with R11F03 driver. (A) Cartoon schematic of the region of interest. (B) Confocal image of adult brain where R11F03-GAL4 is driving UAS-CD8-GFP, stained with nc82 (magenta) and DAPI (teal). Fly light intensity/distribution score is 3/5 and was the only region of the brain identified. Percentage of eggs laid by exposed flies normalized to eggs laid by unexposed flies is shown. UAS-shi^ts^ crossed to R11F03-GAL4 trained by *D. ananassae* at the permissive temperature shows wild-type trained state (C), but at the restrictive temperature shows defective acquisition in the trained state (D). Error bars represent standard error (n = 12 biological replicates) (*p < 0.05).

**Supplementary Figure 12.**
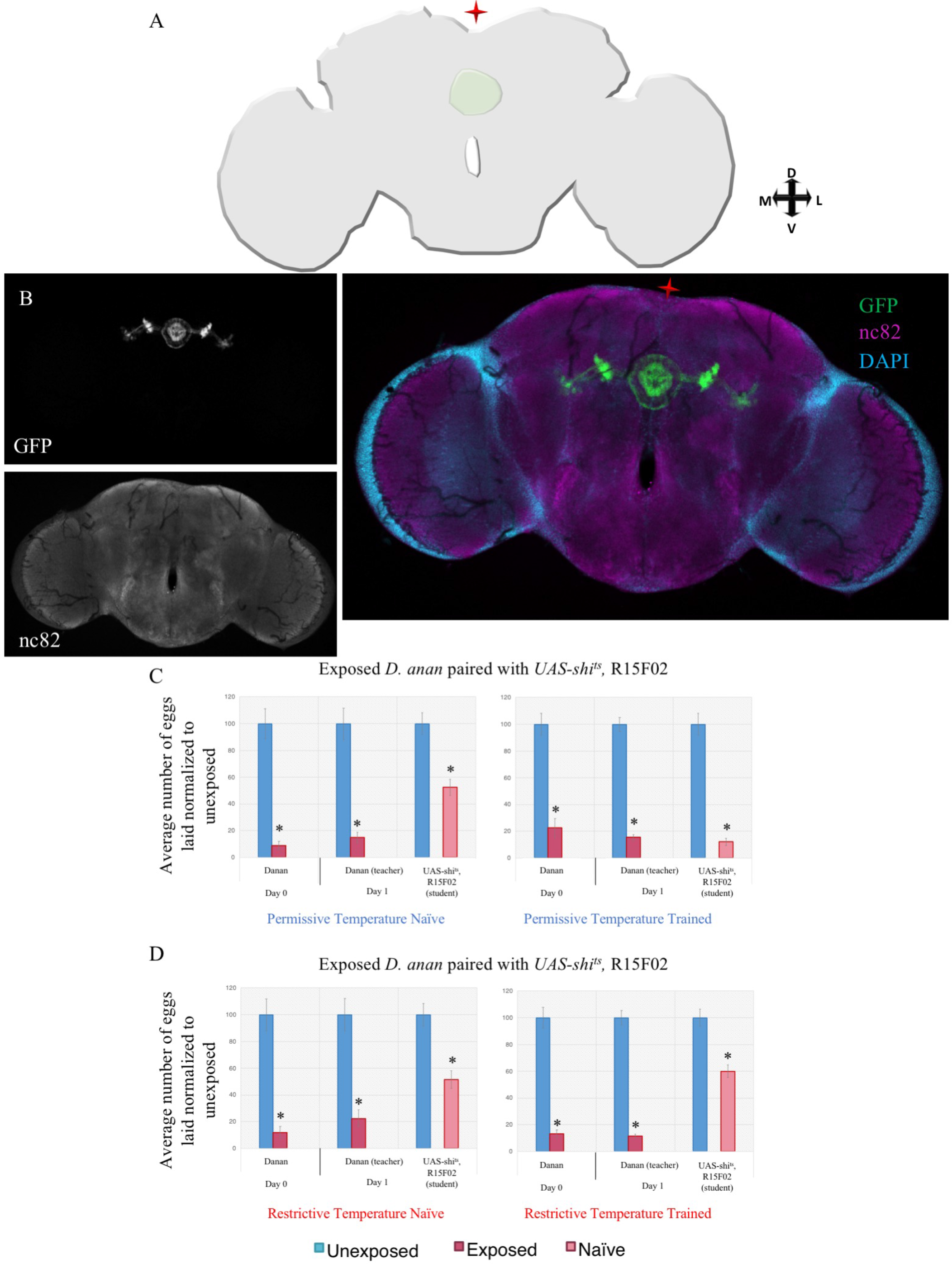
The ellipsoid body is required for dialect learning shown with R15F02 driver. (A) Cartoon schematic of the region of interest. (B) Confocal image of adult brain where R15F02-GAL4 is driving UAS-CD8-GFP, stained with nc82 (magenta) and DAPI (teal). Fly light intensity/distribution score is 5/5 and was the only region of the brain identified. Percentage of eggs laid by exposed flies normalized to eggs laid by unexposed flies is shown. UAS-shi^ts^ crossed to R15F02-GAL4 trained by *D. ananassae* at the permissive temperature shows wild-type trained state (C), but at the restrictive temperature shows defective acquisition in the trained state (D). Error bars represent standard error (n = 12 biological replicates) (*p < 0.05).

**Supplementary Figure 13.**
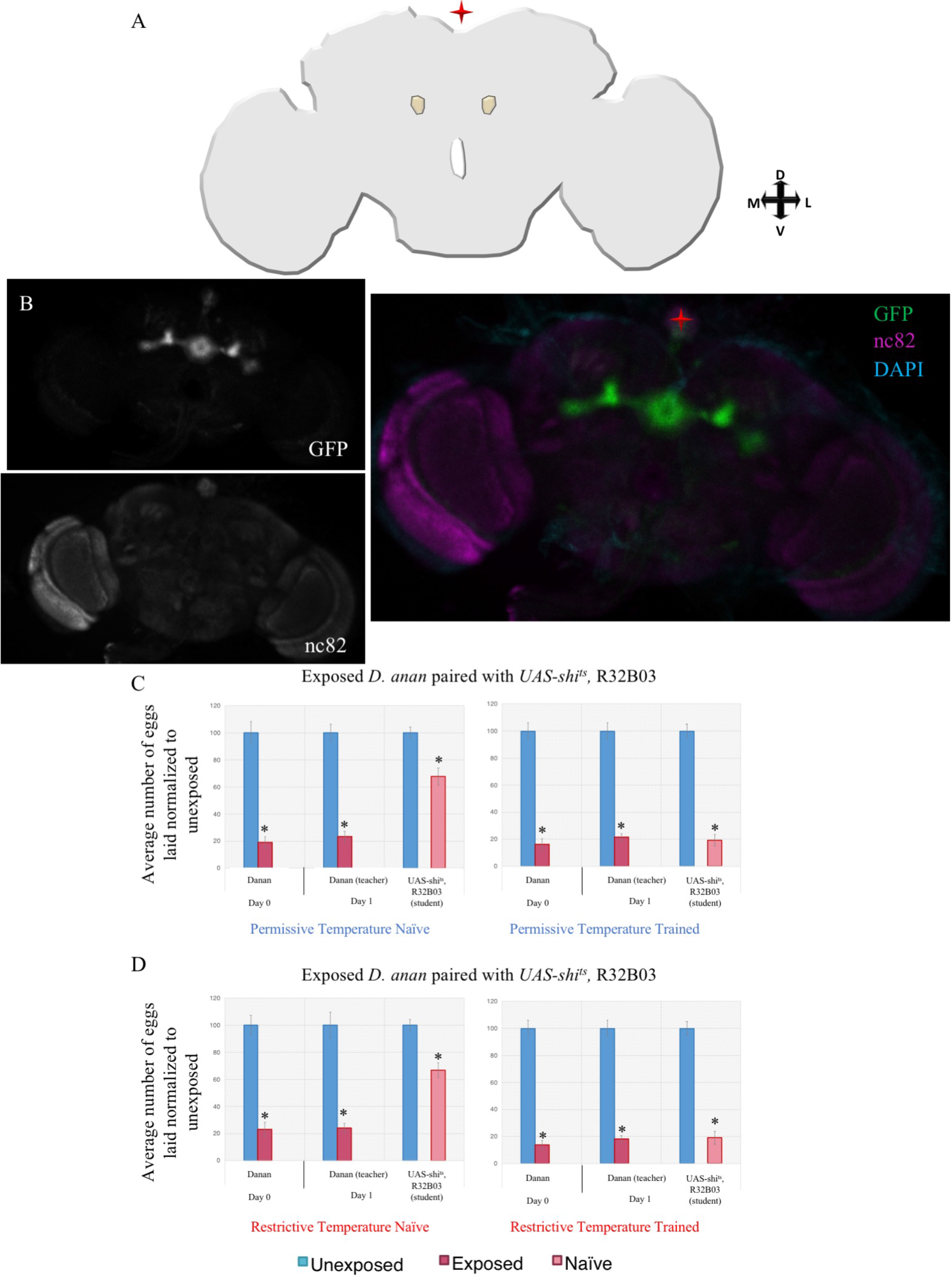
The bulb is dispensable for dialect learning shown with R32B03 driver. (A) Cartoon schematic of the region of interest. (B) Confocal image of adult brain where R32B03-GAL4 is driving UAS-CD8-GFP, stained with nc82 (magenta) and DAPI (teal). Fly light intensity/distribution score is 5/4 and was the only region of the brain identified. Percentage of eggs laid by exposed flies normalized to eggs laid by unexposed flies is shown. Percentage of eggs laid by exposed flies normalized to eggs laid by unexposed flies is shown. UAS-shi^ts^ crossed to R32B03-GAL4 trained by *D. ananassae* at both the permissive temperature (C) and restrictive temperature (D) shows wild-type trained state. Error bars represent standard error (n = 12 biological replicates) (*p < 0.05).

**Supplementary Figure 14.**
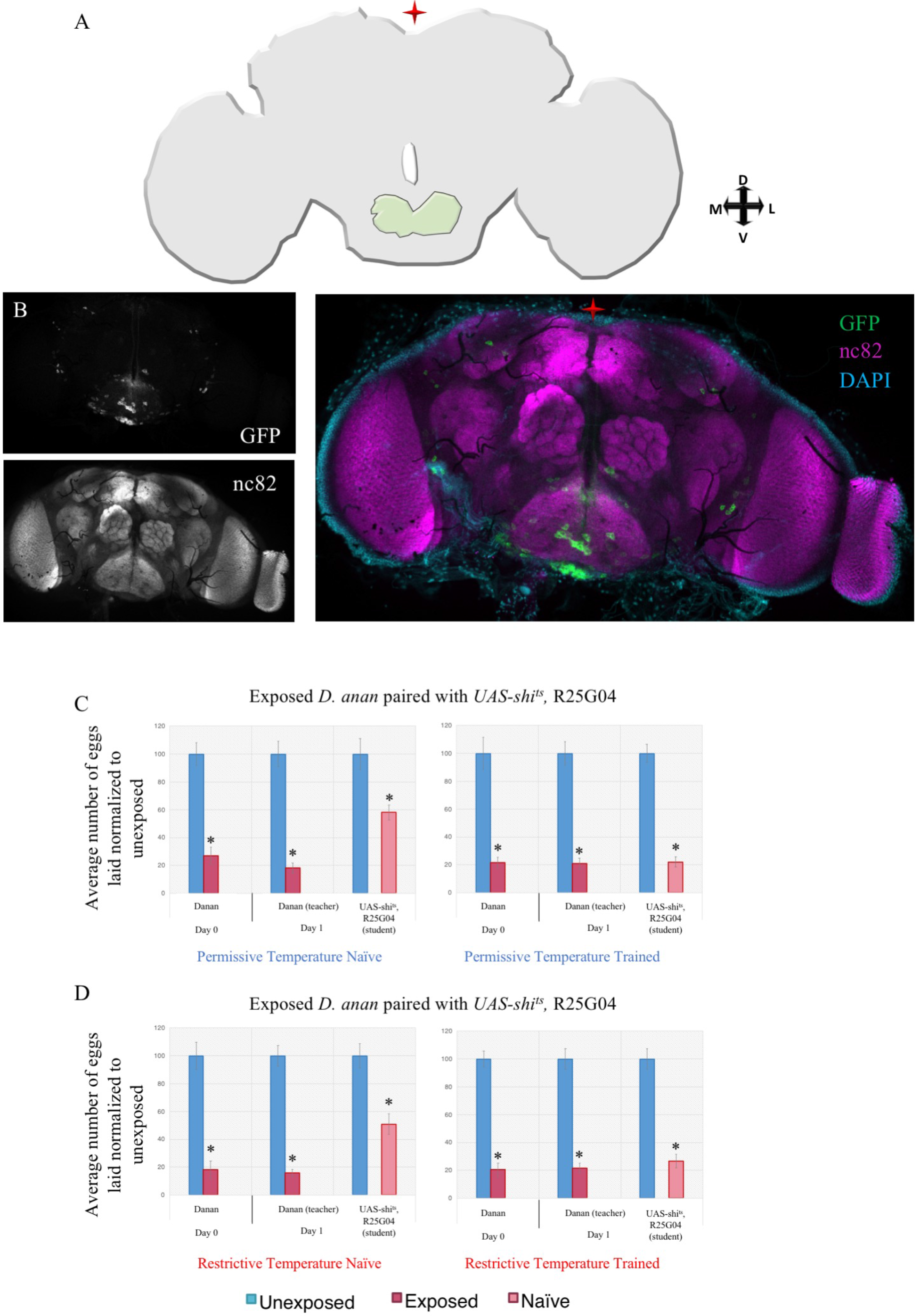
The prow is dispensable for dialect learning shown with R25G04 driver. (A) Cartoon schematic of the region of interest. (B) Confocal image of adult brain where R25G04-GAL4 is driving UAS-CD8-GFP, stained with nc82 (magenta) and DAPI (teal). Fly light intensity/distribution score is 4/4 and was the only region of the brain identified. Percentage of eggs laid by exposed flies normalized to eggs laid by unexposed flies is shown. UAS-shi^ts^ crossed to R25G04-GAL4 trained by *D. ananassae* at both the permissive temperature (C) and restrictive temperature (D) shows wild-type trained state. Error bars represent standard error (n = 12 biological replicates) (*p < 0.05).

**Supplementary Figure 15.**
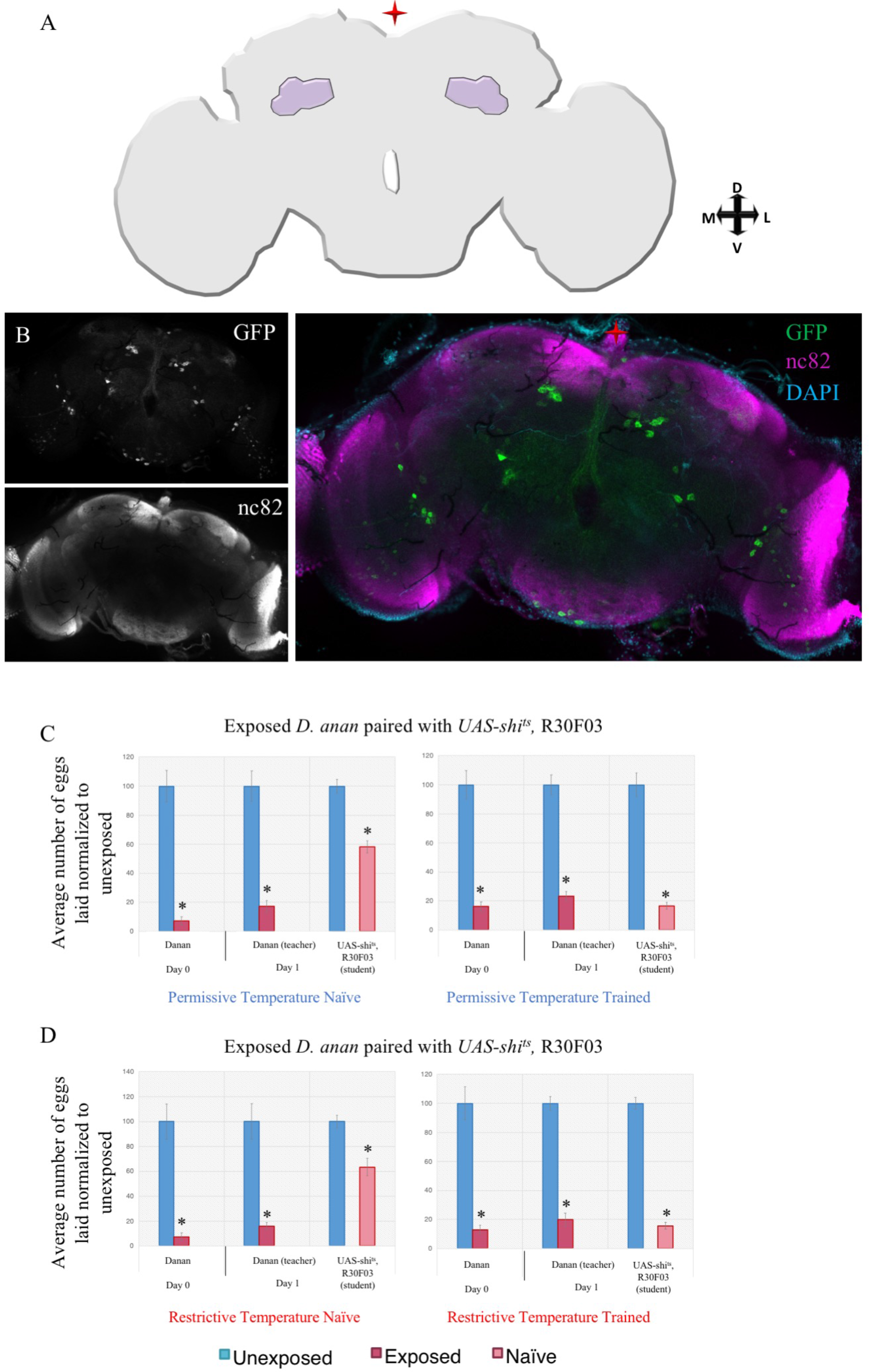
The superior clamp is dispensable for dialect learning shown with R30F03 driver. (A) Cartoon schematic of the region of interest. (B) Confocal image of adult brain where R30F03-GAL4 is driving UAS-CD8-GFP, stained with nc82 (magenta) and DAPI (teal). Fly light intensity/distribution score is 3/3 and was the only region of the brain identified. Percentage of eggs laid by exposed flies normalized to eggs laid by unexposed flies is shown. UAS-shi^ts^ crossed to R30F03-GAL4 trained by *D. ananassae* at both the permissive temperature (C) and restrictive temperature (D) shows wild-type trained state. Error bars represent standard error (n = 12 biological replicates) (*p < 0.05).

**Supplementary Figure 16.**
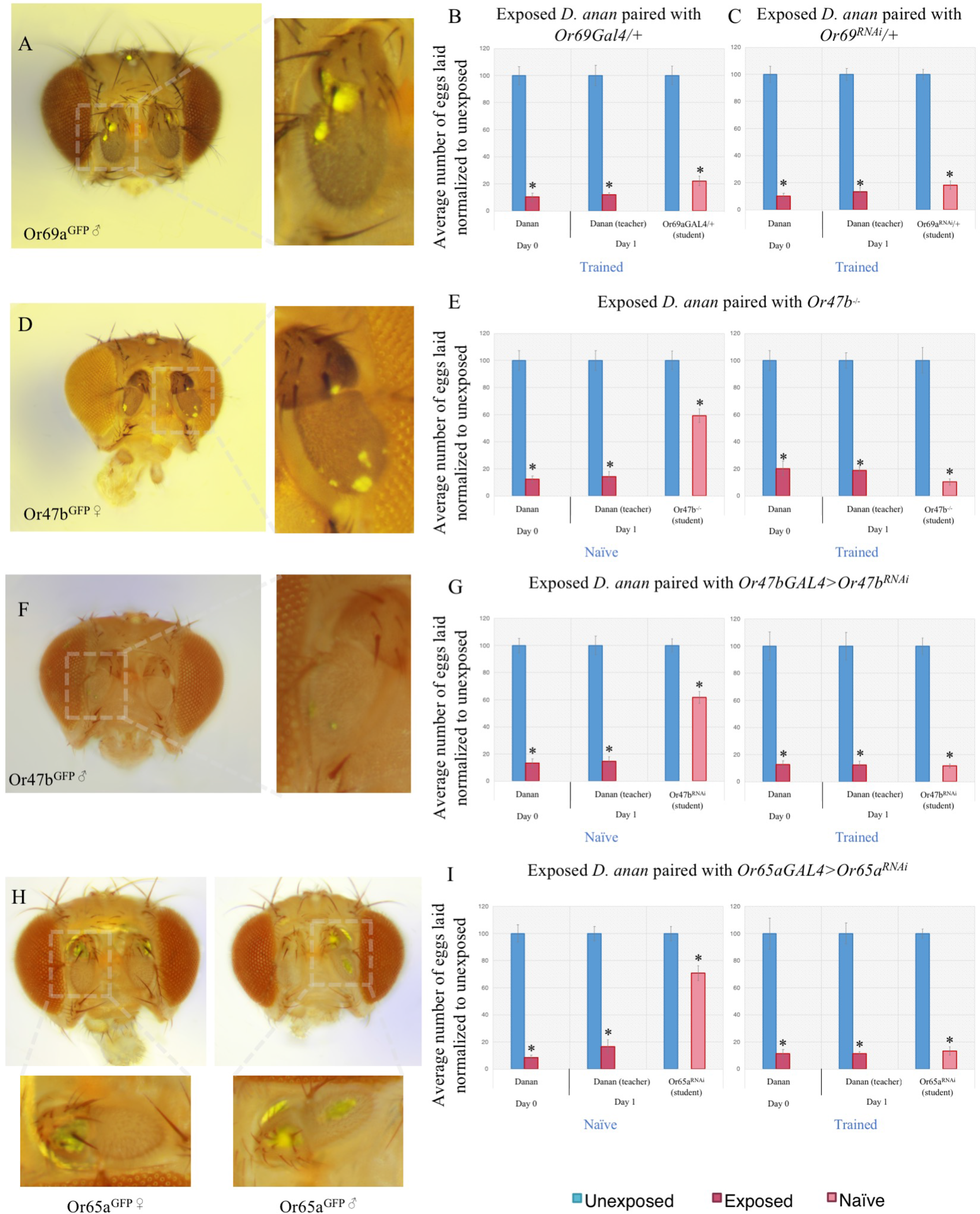
Further evidence implicating the role of *Or69a* in dialect learning. (A) Expression of Or69a using the Or69a^GFP^ construct highlighting expression in the antenna in males. Dotted boxes indicate regions of magnification. Percentage of eggs laid by exposed flies normalized to eggs laid by unexposed flies is shown. Or69aGAL4/+ and Or69a^RNAi^/+ show wild-type trained behavior (B-C). (D) Female expression of Or49b using the Or49b ^GFP^ construct highlighting expression in the antenna. Or49b^−/−^ shows wild-type naïve behavior, and are able to learn the dialect from *D. ananassae* following training (E). Female expression of Or49b using the Or49b ^GFP^ construct highlighting expression in the antenna (F). Or49b^GAL4^ driving Or49b^RNAi^ shows wild-type naïve behavior and can learn the dialect from *D. ananassae* following training (G). Male and female expression of Or65a using the Or65a ^GFP^ construct highlighting expression in the antenna (H). Or65a^GAL4^ driving Or65a^RNAi^ shows wild-type naïve behavior and can learn the dialect from *D. ananassae* following training (I) Error bars represent standard error (n = 12 biological replicates) (*p < 0.05).

**Supplementary Figure 17.**
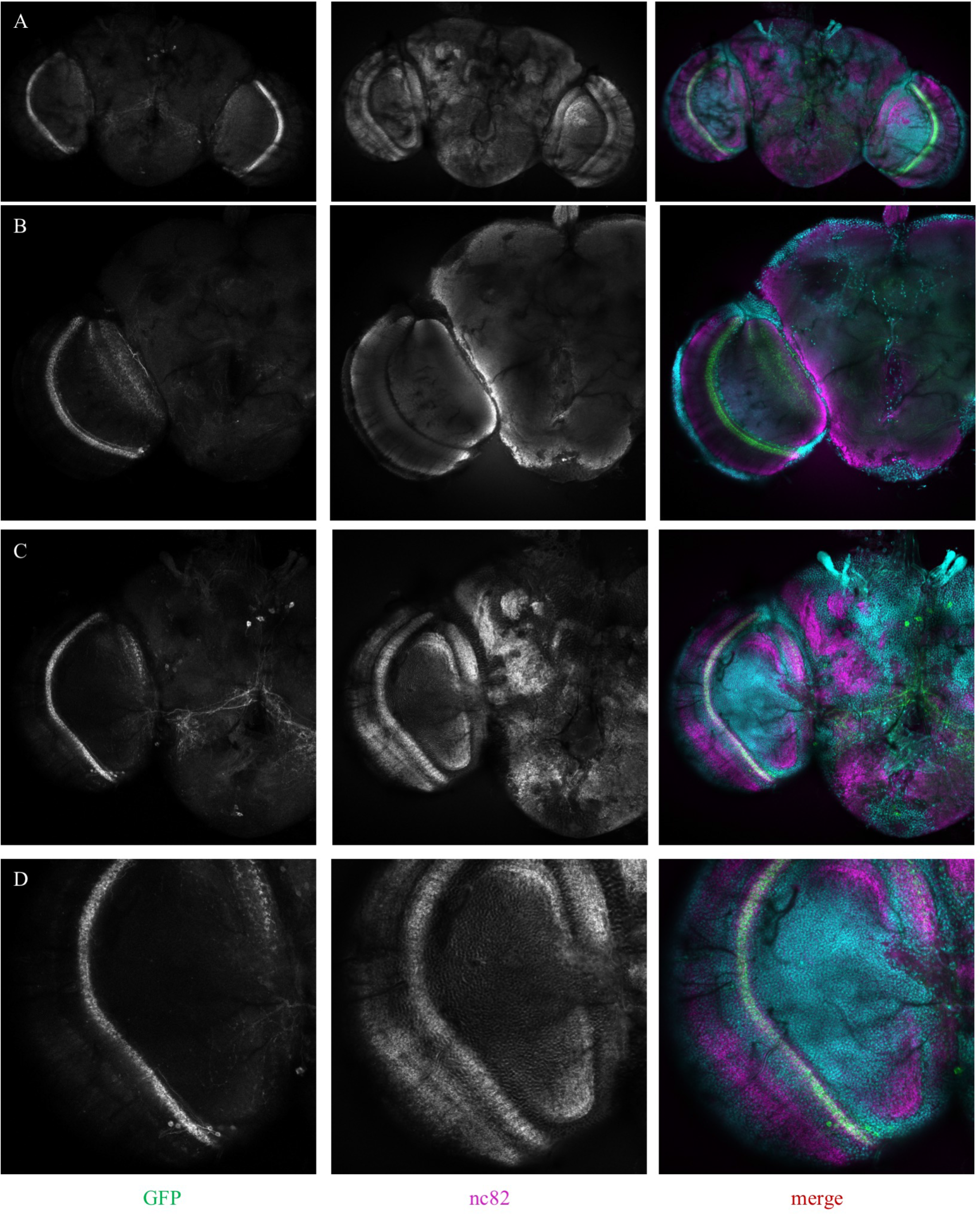
Expression pattern of the L1 line used. (A-D) Confocal image of adult brain where L1-GAL4 is driving UAS-CD8-GFP, stained with nc82 (magenta) and DAPI (teal).

**Supplementary Figure 18.**
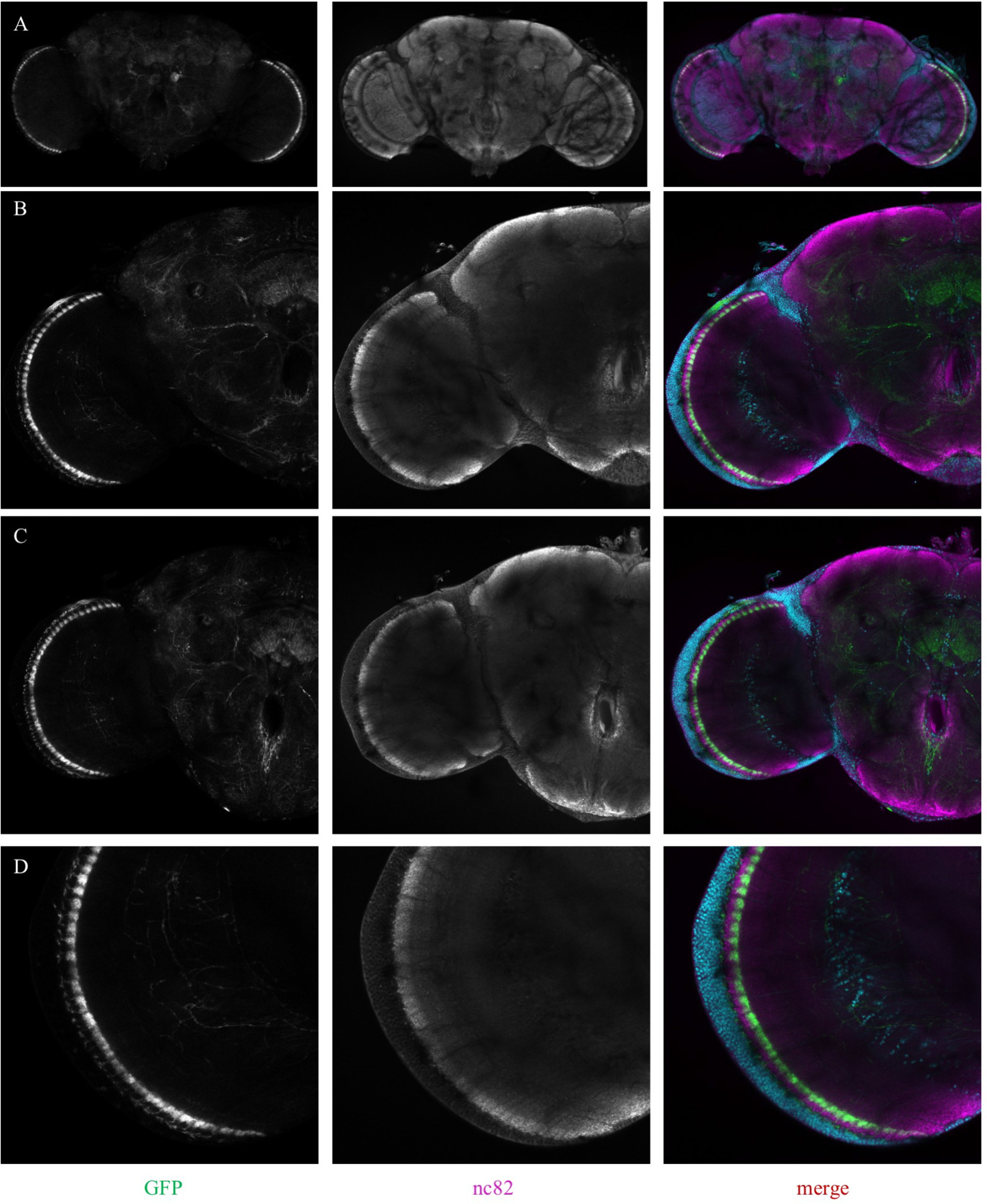
Expression pattern of the L2 line used. (A-D) Confocal image of adult brain where L2-GAL4 is driving UAS-CD8-GFP, stained with nc82 (magenta) and DAPI (teal).

**Supplementary Figure 19.**
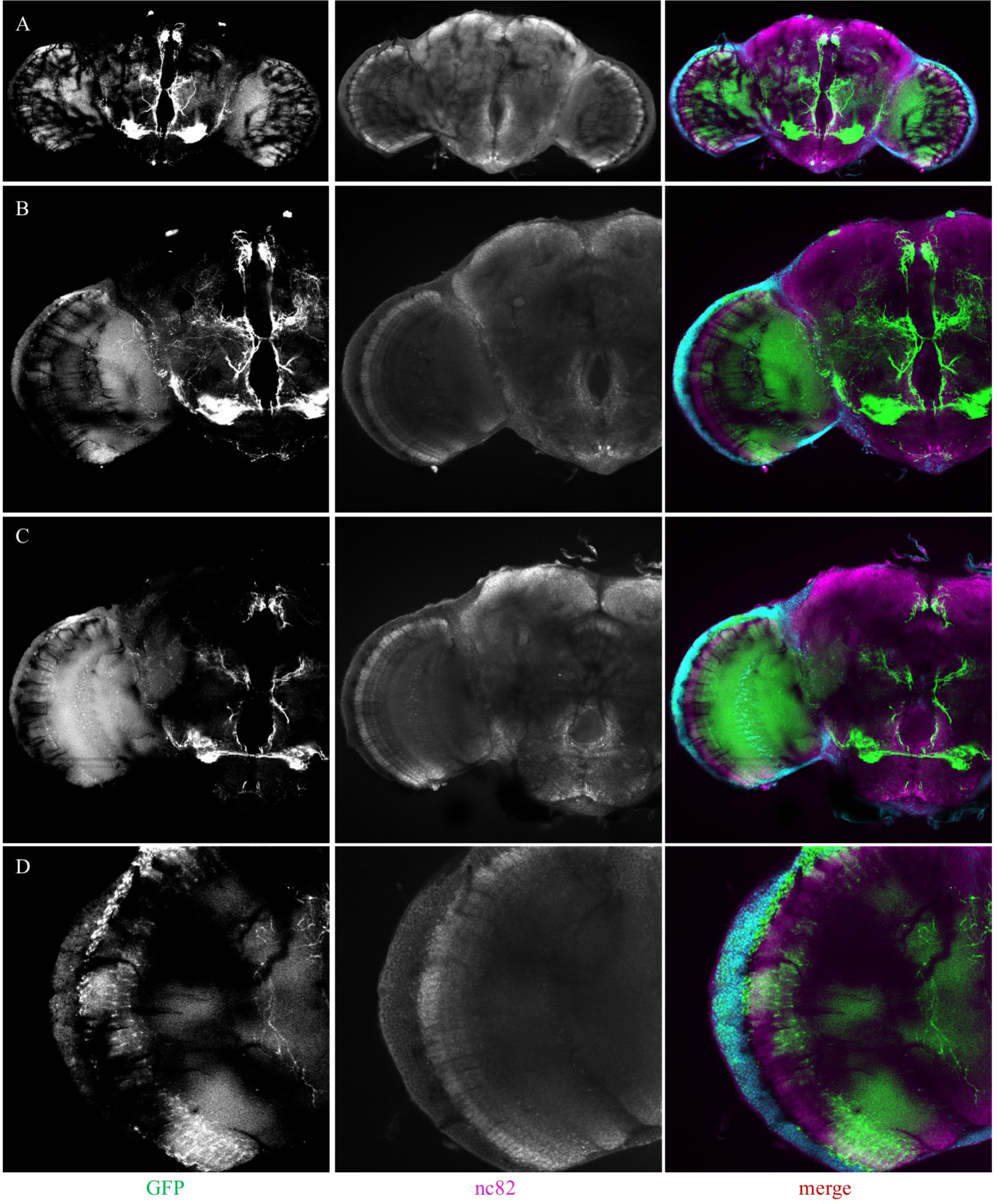
Expression pattern of the L3 line used. (A-D) Confocal image of adult brain where L3-GAL4 is driving UAS-CD8-GFP, stained with nc82 (magenta) and DAPI (teal).

**Supplementary Figure 20.**
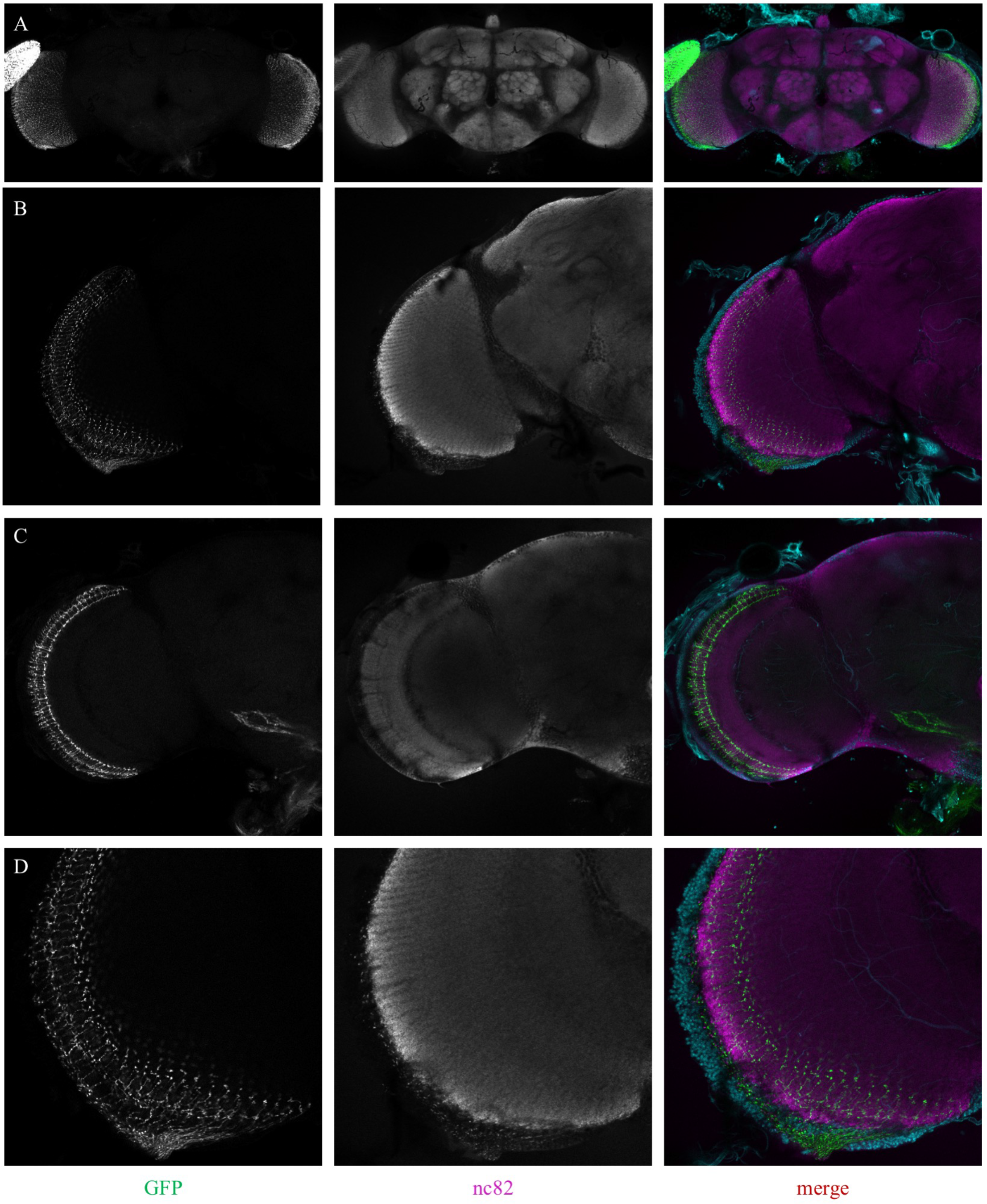
Expression pattern of the L4^0987^ line used. (A-D) Confocal image of adult brain where L4^0987^-GAL4 is driving UAS-CD8-GFP, stained with nc82 (magenta) and DAPI (teal).

**Supplementary Figure 21.**
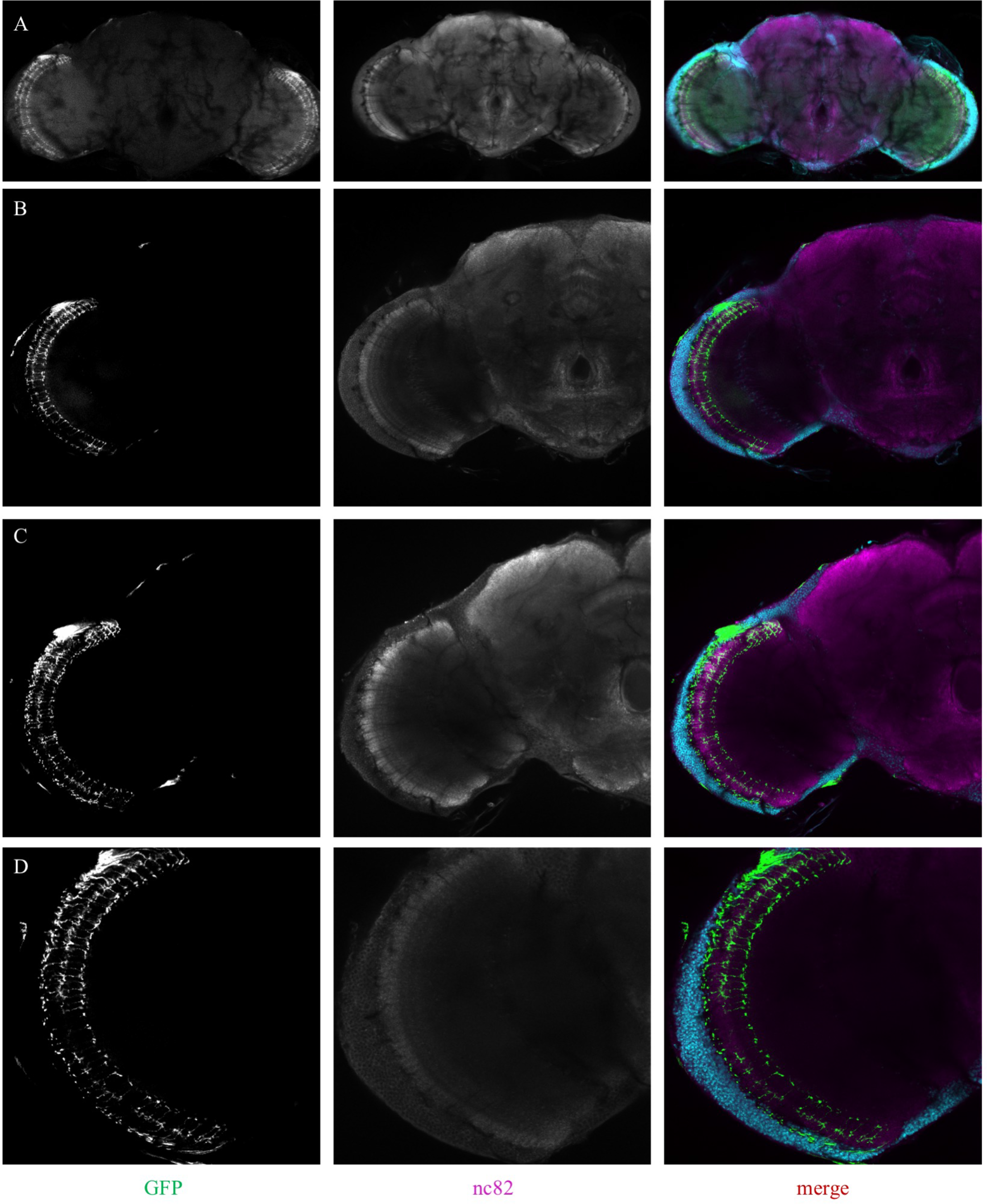
Expression pattern of the splitL4 line used. (A-D) Confocal image of adult brain where splitL4-GAL4 is driving UAS-CD8-GFP, stained with nc82 (magenta) and DAPI (teal).

**Supplementary Figure 22.**
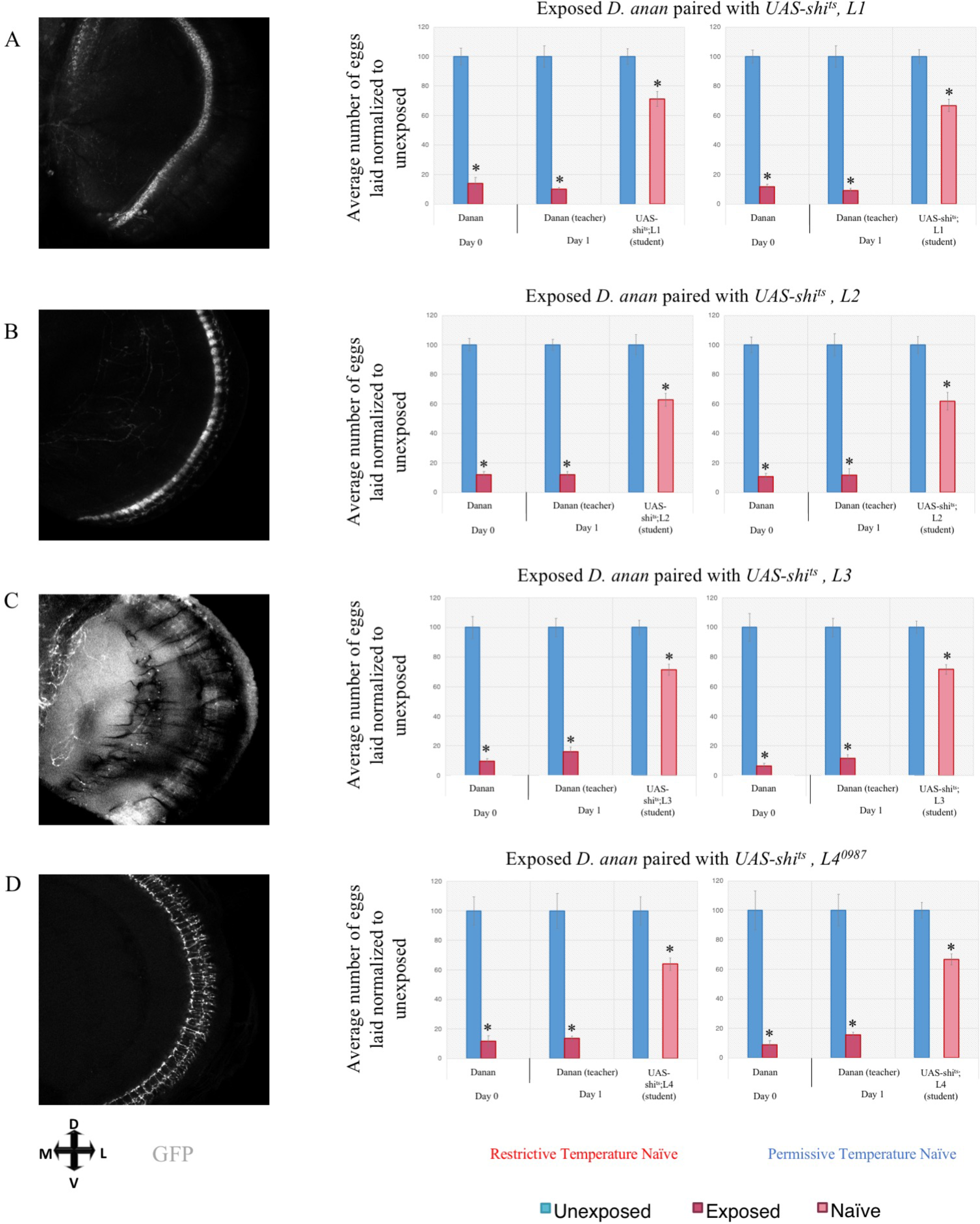
Further evidence implicating the role of motion-detecting circuitry in dialect learning using UAS Shi^ts^. Dialect learning is performed at either the permissive (22°C) or restrictive (30°C) temperature, while the wasp exposure and social learning period is performed exclusively at the restrictive temperature. Percentage of eggs laid by exposed flies normalized to eggs laid by unexposed flies is shown. (A) UAS-shi^ts^ crossed to *L1-GAL4* in the naïve state at restrictive and permissive temperatures shows wild-type naïve state. Expression pattern of *L1-GAL4* also shown. (B) UAS-shi^ts^ crossed to *L2-GAL4* in the naïve state at restrictive and permissive temperatures shows wild-type naïve state. Expression pattern of *L2-GAL4* also shown. (C) UAS-shi^ts^ crossed to *L3-GAL4* in the naïve state at restrictive and permissive temperatures shows wild-type naïve state. Expression pattern of *L3-GAL4* also shown. (D) UAS-shi^ts^ crossed to *L4^0987^-GAL4* in the naïve state at restrictive and permissive temperatures shows wild-type naïve state. Expression pattern of *L4^0987^-GAL4* also shown. Error bars represent standard error (n = 12 biological replicates) (*p < 0.05).

**Supplementary Figure 23.**
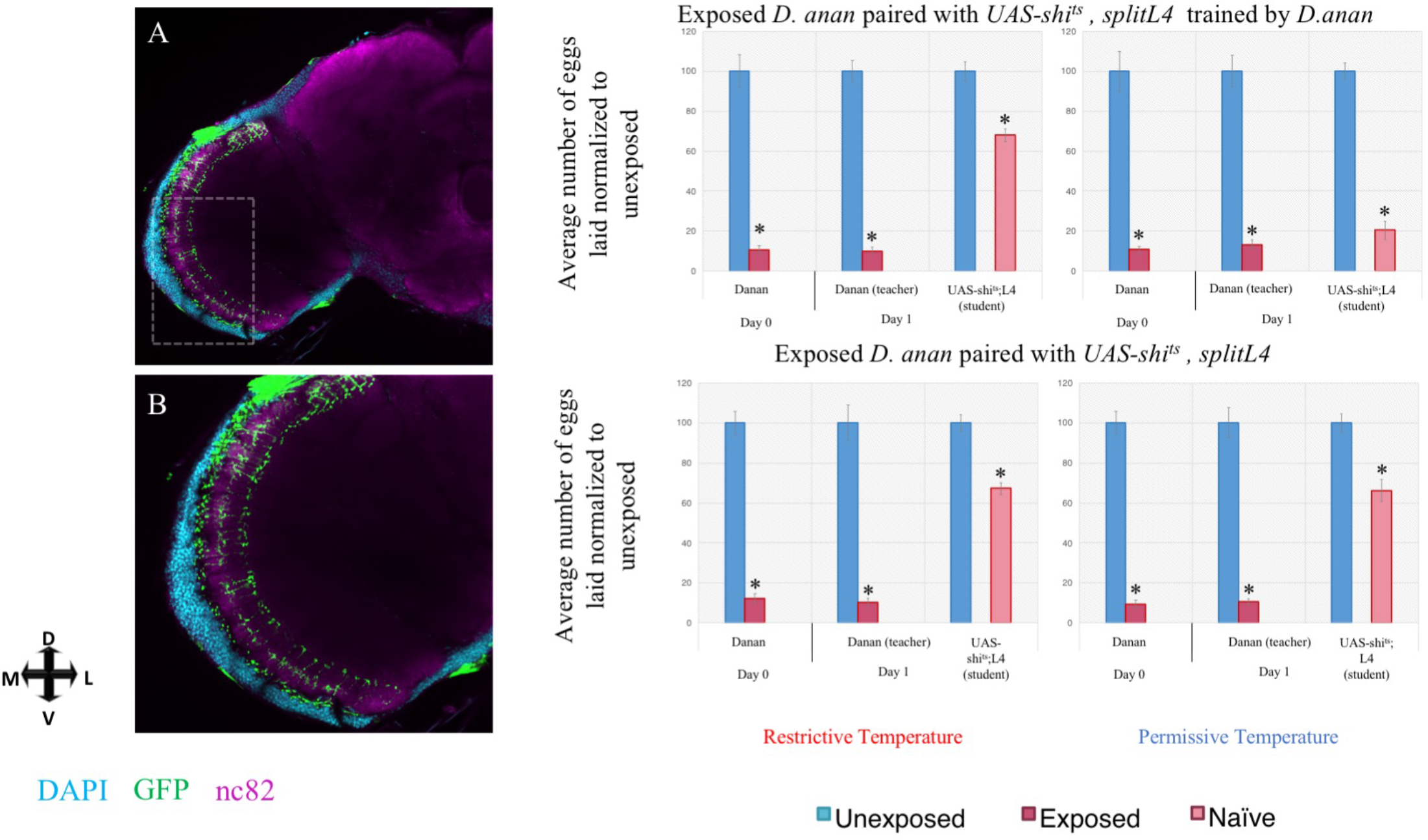
L4 motion sensing neurons in the optic lobe are required for dialect learning. Dialect learning is performed at either the permissive (22°C) or restrictive (30°C) temperature, while the wasp exposure and social learning period is performed exclusively at the permissive temperature. (A) Confocal image of adult brain where splitL4-GAL4 is driving UAS-CD8-GFP, stained with nc82 (magenta) and DAPI (teal). A magnification is shown of boxed area. See also, figure S 21. UAS-shits crossed to splitL4-GAL4 trained by *D. ananassae* at the permissive temperature shows wild-type trained state, but at the restrictive temperature shows defective acquisition in the trained state. (B) UAS-shi^ts^ crossed to split*L4-GAL4* in the naïve state at restrictive and permissive temperatures shows wild-type naïve state. Expression pattern of split*L4-GAL4* also shown. Error bars represent standard error (n = 12 biological replicates) (*p < 0.05).

**Supplementary Figure 24.**
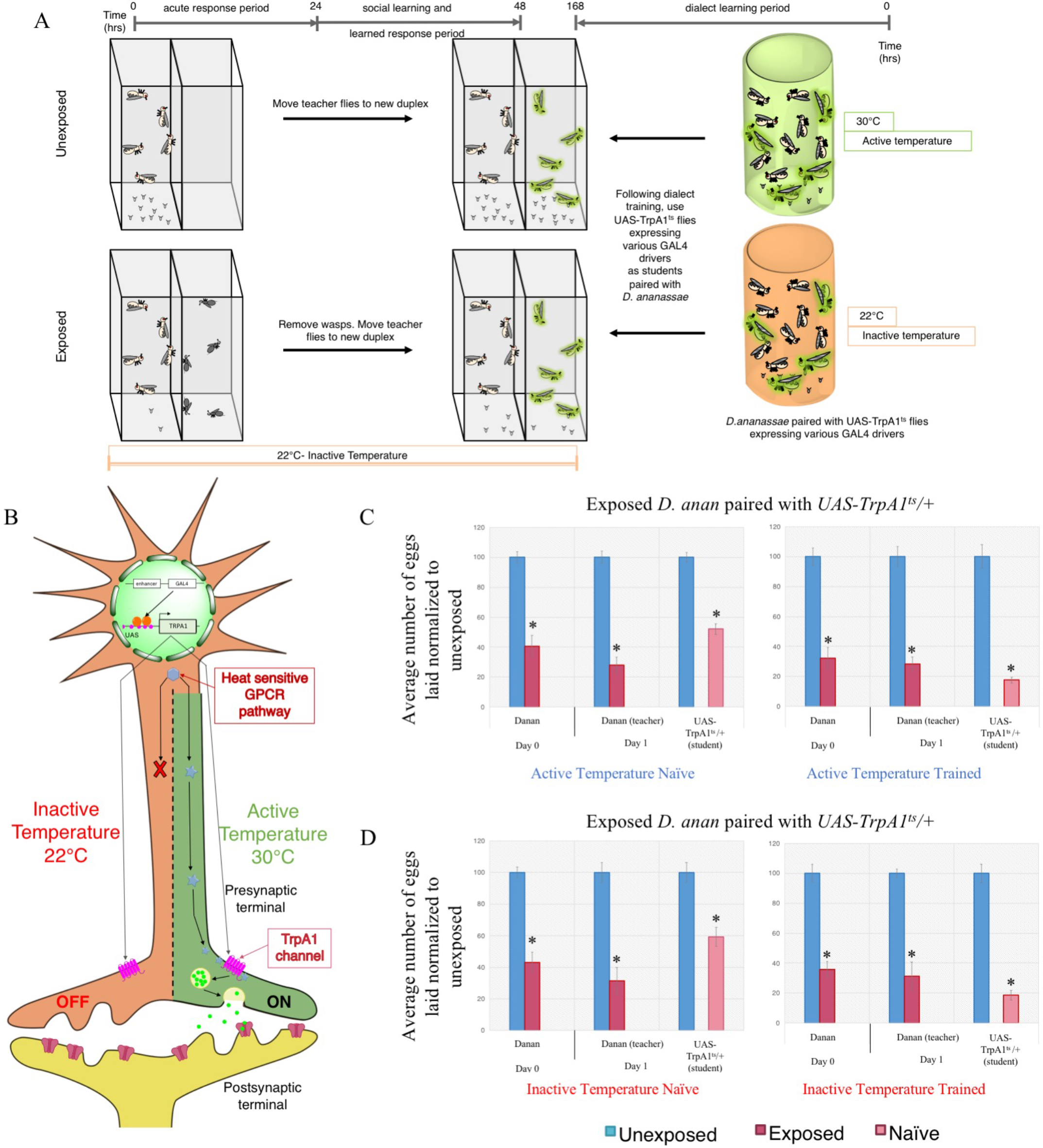
Dialect learning can be analyzed using UAS-TrpA1^ts^. Standard experimental design using UAS-TrpA1^ts^ in conjunction with various GAL4 drivers. Dialect learning is performed at either the inactive (22°C) or active (30°C) temperature, while the wasp exposure and social learning period is performed exclusively at the restrictive temperature. Schematic of the UAS-TrpA1^ts^ expression in neurons, where the restrictive temperature turns on the neurons of interest in the absence of a given stimulus and the restrictive temperature does not affect these neurons. (C) Percentage of eggs laid by exposed flies normalized to eggs laid by unexposed flies is shown. UAS-TrpA1^ts^ outcrossed to *Canton S* trained by *D. ananassae* at either the permissive temperature (C) or the restrictive temperature (D) show wild-type dialect acquisition in both the naïve and trained states. Error bars represent standard error (n = 12 biological replicates) (*p < 0.05).

**Supplementary Figure 25.**
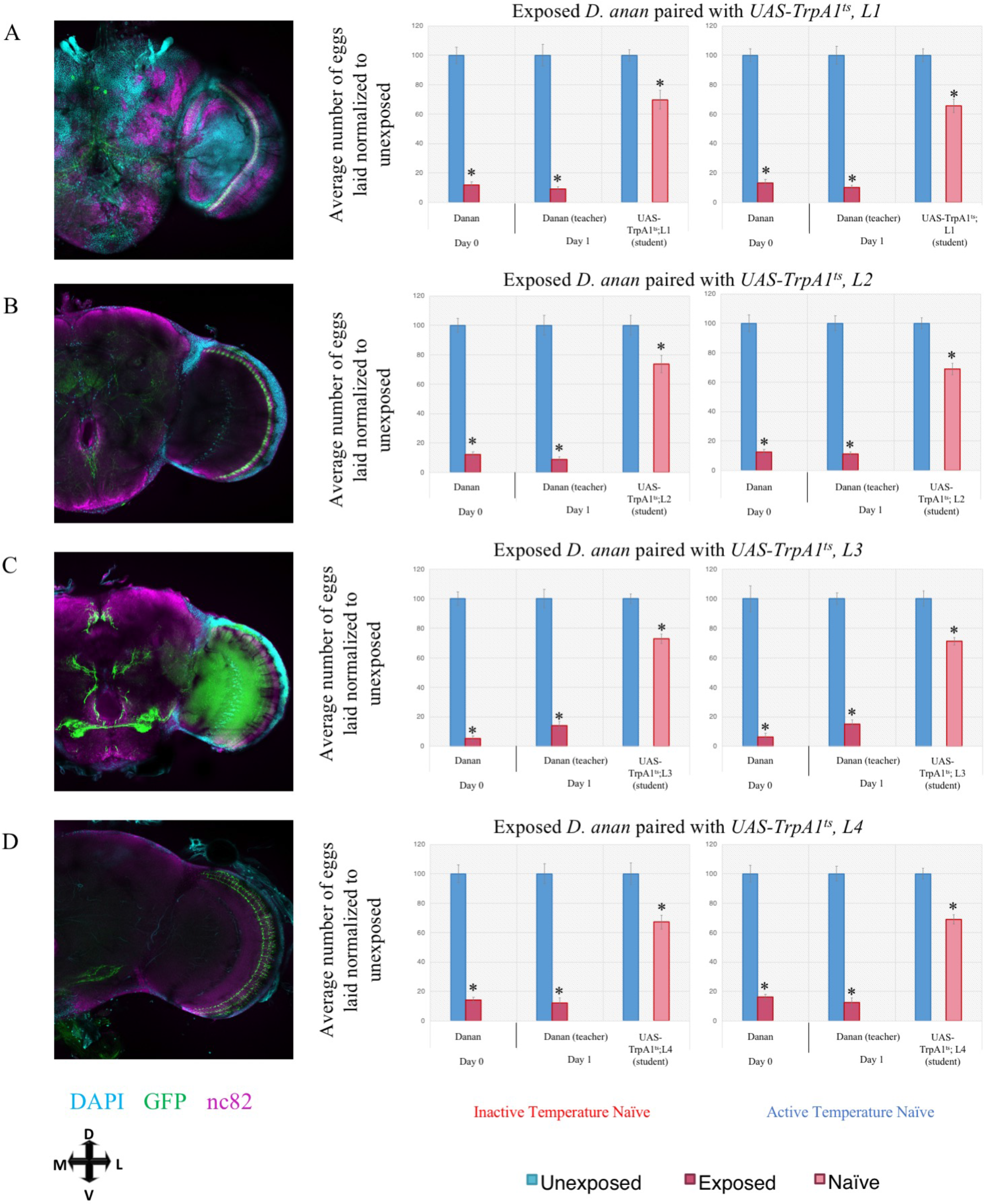
Further evidence implicating the role of motion-detecting circuitry in dialect learning using UAS TrpA1. Dialect learning is performed at either the active (30°C) or inactive (22°C) temperature exclusively in the dark, while the wasp exposure and social learning period is performed exclusively at the restrictive temperature. Percentage of eggs laid by exposed flies normalized to eggs laid by unexposed flies is shown. (A) UAS-TrpA1^ts^ crossed to *L1-GAL4* in the naïve state at restrictive and permissive temperatures shows wild-type naïve state. Expression pattern of *L1-GAL4* also shown. (B) UAS-TrpA1^ts^ crossed to *L2-GAL4* in the naïve state at restrictive and permissive temperatures shows wild-type naïve state. Expression pattern of *L2-GAL4* also shown. (C) UAS-TrpA1^ts^ crossed to *L3-GAL4* in the naïve state at active and inactive temperatures shows wild-type naïve state. Expression pattern of *L3-GAL4* also shown. (D) UAS-TrpA1^ts^ crossed to *L4*^0987^*-GAL4* in the naïve state at active and inactive temperatures shows wild-type naïve state. Expression pattern of *L4*^0987^*-GAL4* also shown. Error bars represent standard error (n = 12 biological replicates) (*p < 0.05).

**Supplementary Figure 26.**
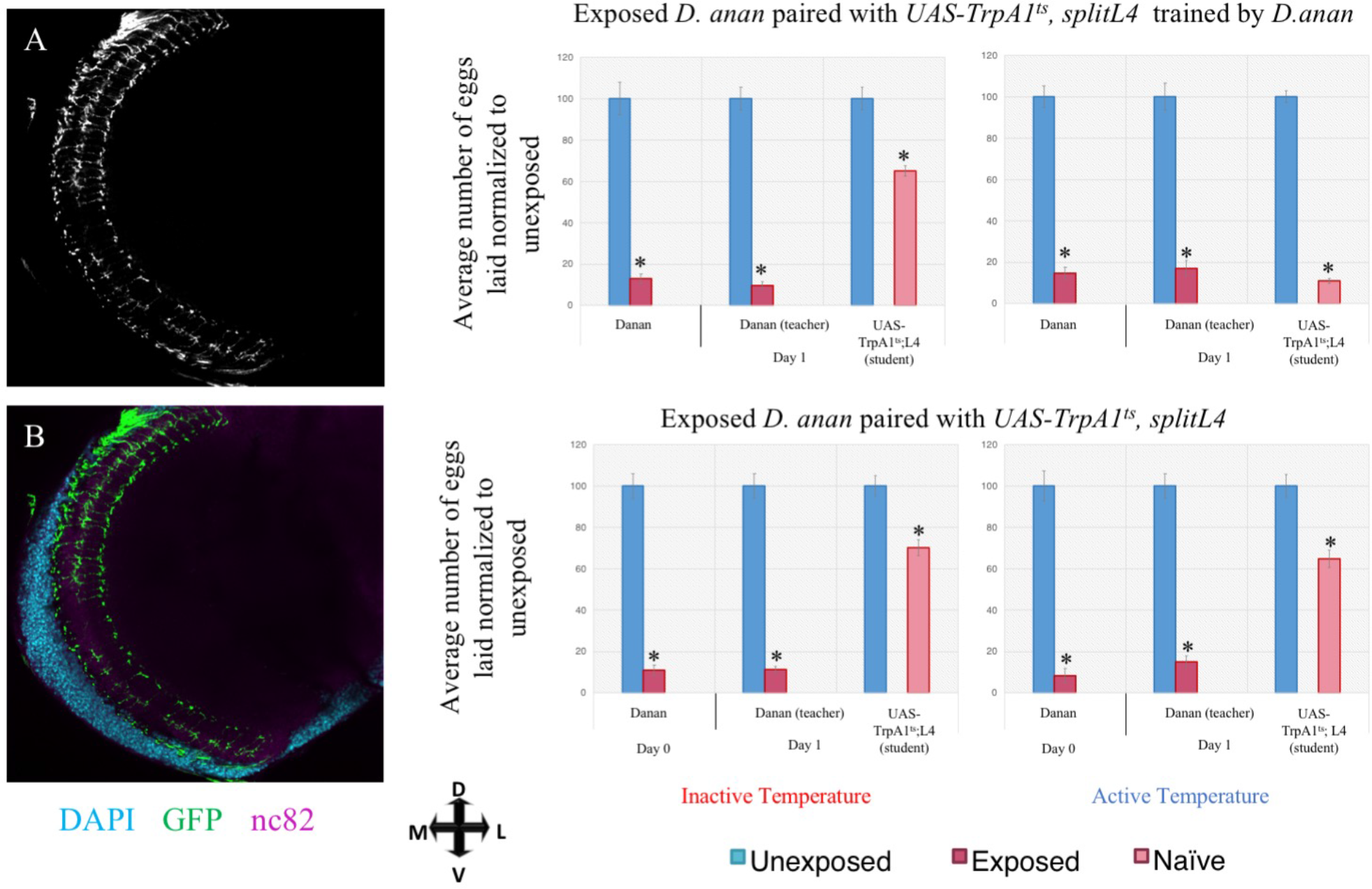
L4 motion detecting neurons in the optic lobe are sufficient for dialect learning. Dialect learning is performed at either the active (30°C) or inactive (22°C) temperature exclusively in the dark, while the wasp exposure and social learning period is performed exclusively at the restrictive temperature. (A) Confocal image of adult brain where splitL4-GAL4 is driving UAS-CD8-GFP, stained with nc82 (magenta) and DAPI (teal). UAS-TrpA1^ts^ crossed to splitL4-GAL4 trained by *D. ananassae* at the activation temperature shows acquisition of the dialect, but not at the inactive temperature, which demonstrates the need for full spectrum light. (B) UAS-TrpA1^ts^ crossed to *splitL4-GAL4* in the naïve state at active and inactive temperatures shows wild-type naïve state. Expression pattern of *splitL4-GAL4* shown. Error bars represent standard error (n = 12 biological replicates) (*p < 0.05).

**Supplementary Figure 27.**
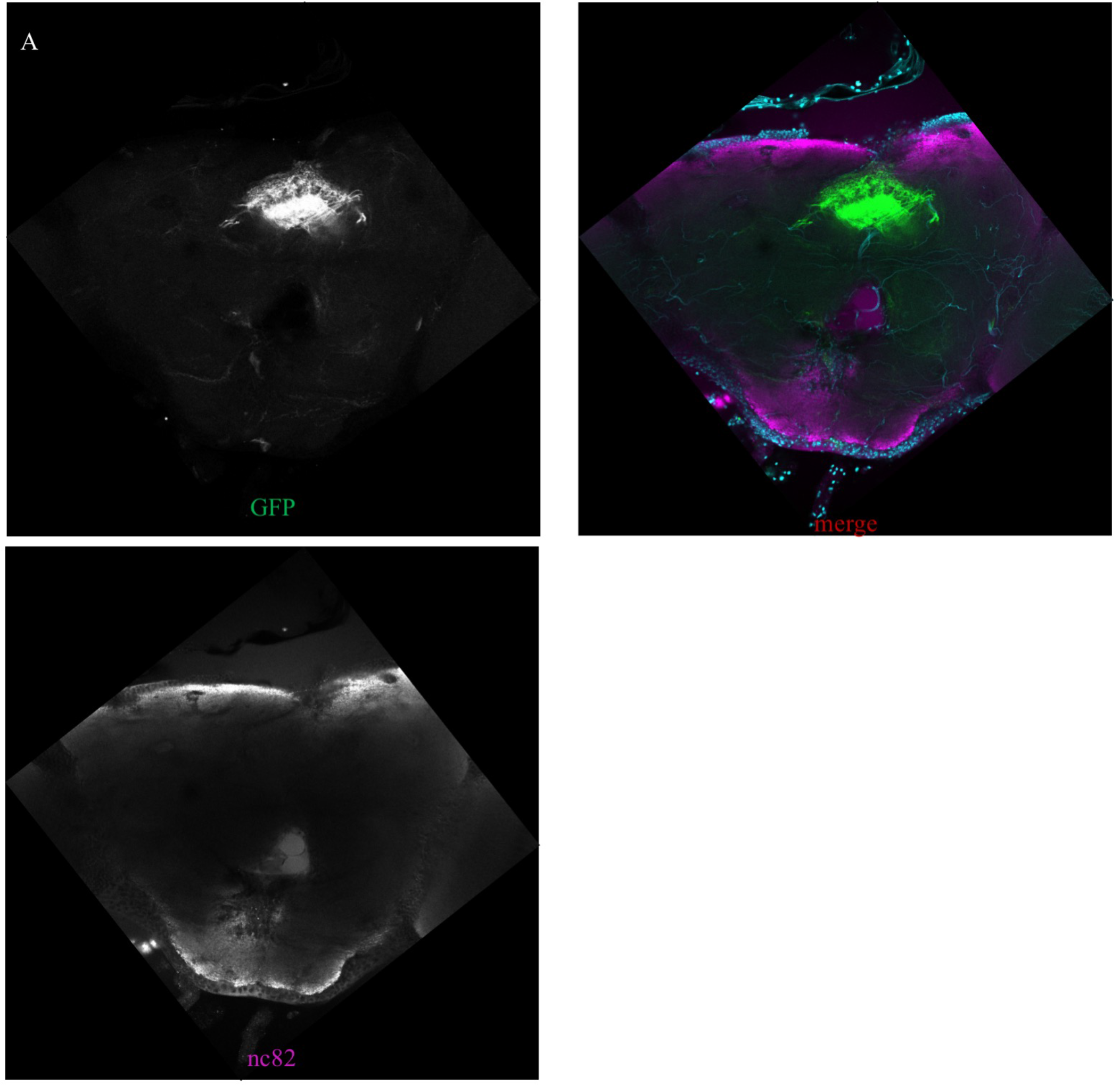
Expression pattern of the fan-shaped body driver line R75G12. (A) Confocal image of adult brain where R75G12 is driving UAS-CD8-GFP, stained with nc82 (magenta) and DAPI (teal).

**Supplementary Figure 28.**
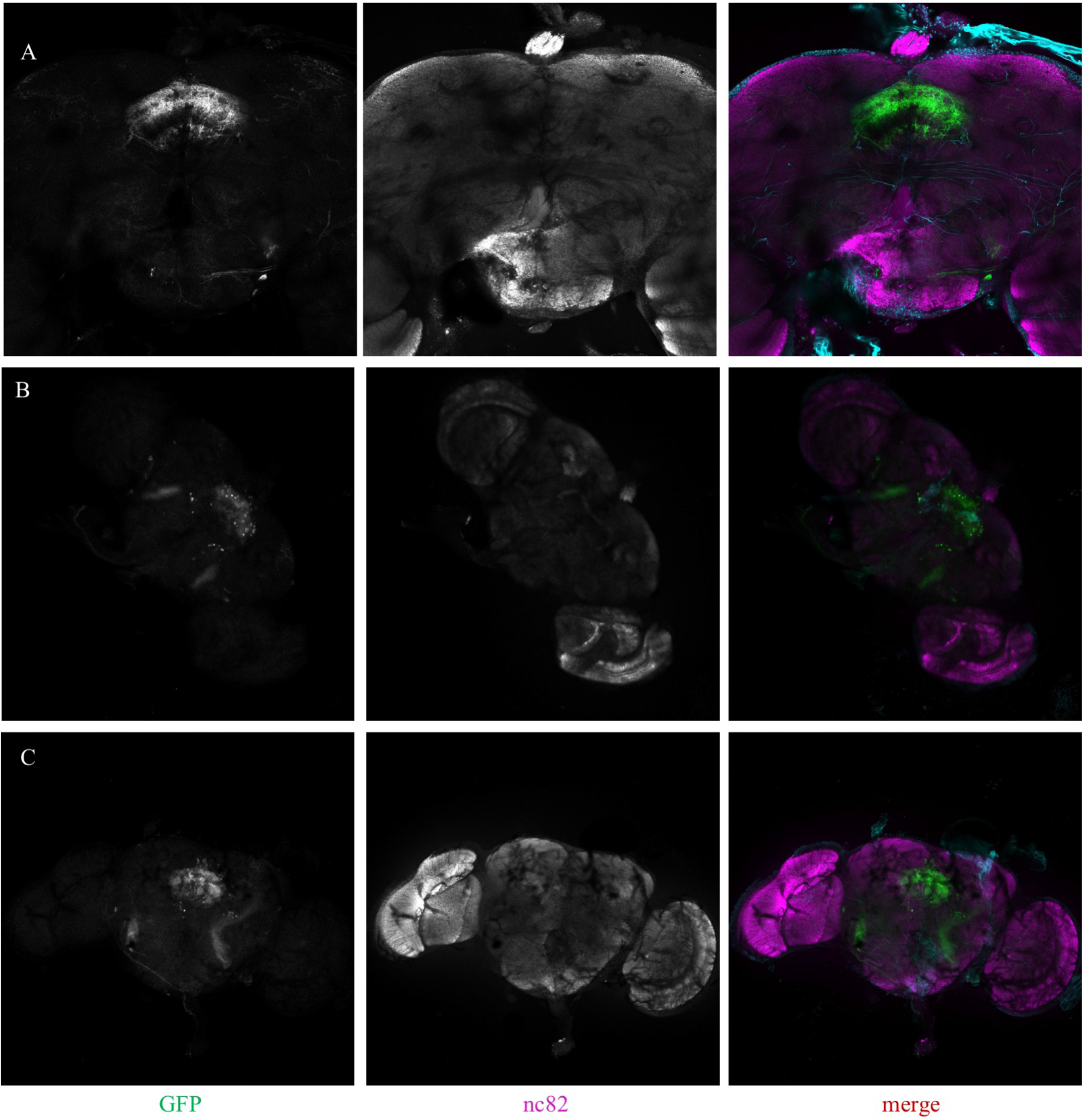
Expression pattern of the fan-shaped body driver line R38E07. (A-C) Confocal image of adult brain where R38E07 is driving UAS-CD8-GFP, stained with nc82 (magenta) and DAPI (teal).

**Supplementary Figure 29.**
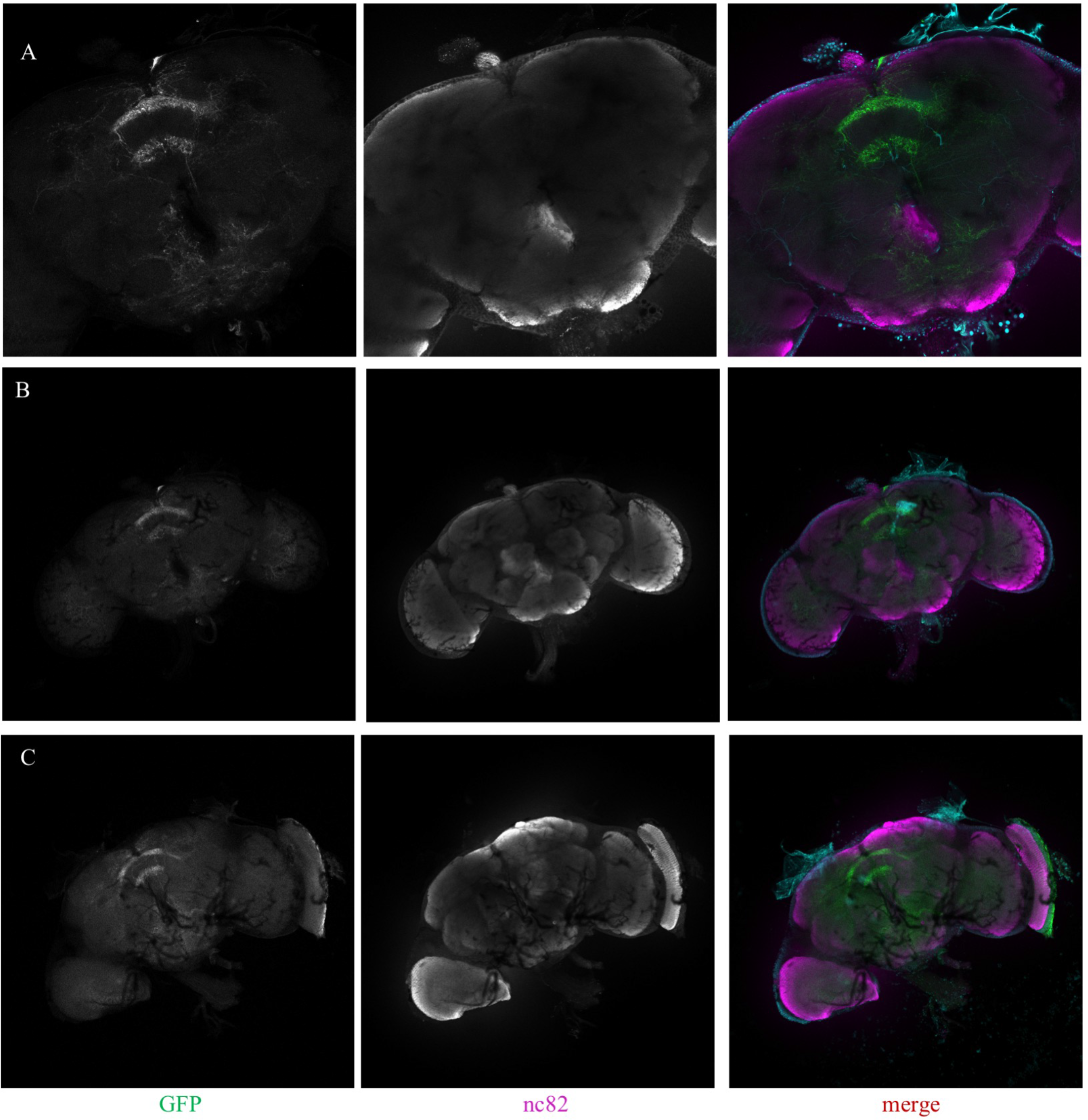
Expression pattern of the fan-shaped body driver line R89E07. (A-C) Confocal image of adult brain where R89E07 is driving UAS-CD8-GFP, stained with nc82 (magenta) and DAPI (teal).

**Supplementary Figure 30.**
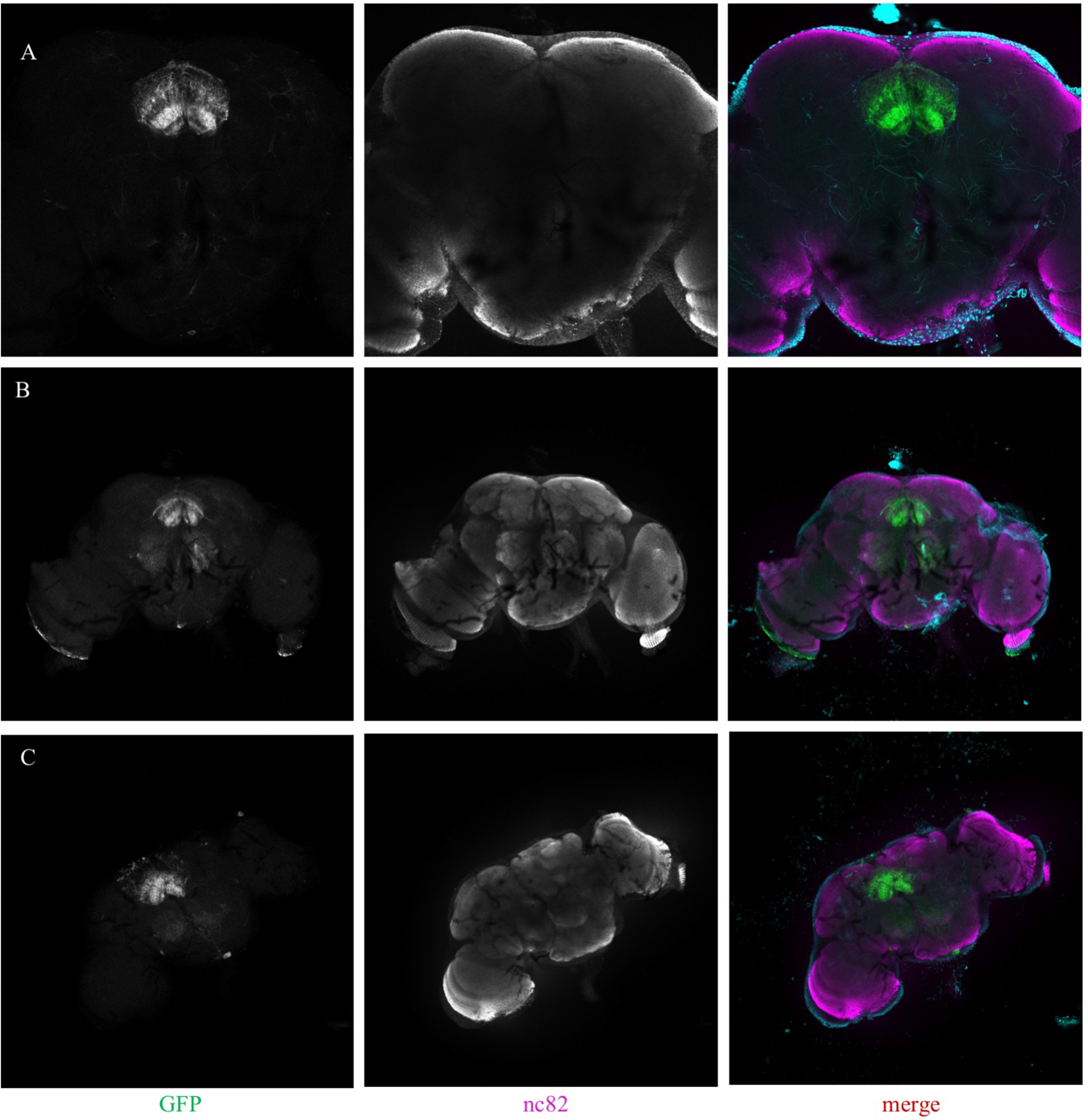
Expression pattern of the fan-shaped body driver line R49H02. (A-C) Confocal image of adult brain where R49H02 is driving UAS-CD8-GFP, stained with nc82 (magenta) and DAPI (teal).

**Supplementary Figure 31.**
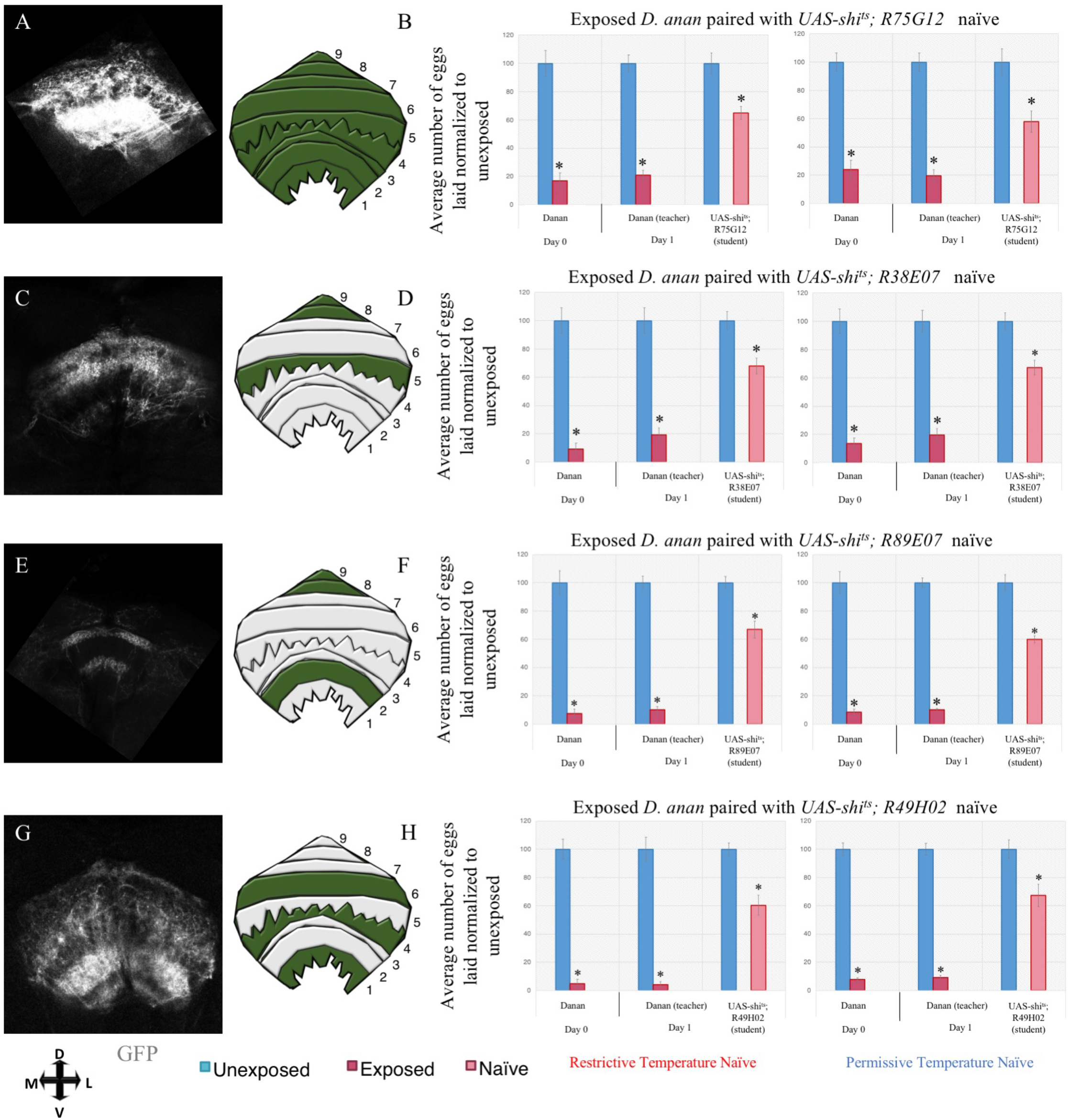
Further evidence indicating that Region 5 of the fan-shaped body is necessary for dialect learning. Dialect learning is performed at either the permissive (22°C) or restrictive (30°C) temperature, while the wasp exposure and social learning period is performed exclusively at the restrictive temperature. Percentage of eggs laid by exposed flies normalized to eggs laid by unexposed flies is shown. (A) Expression and schematic of expression pattern of the R75G12 FSB driver shows pan-FSB marking. (B) UAS-shi^ts^ crossed to R75G12 trained by *D. ananassae* at the permissive and restrictive temperature shows wild-type untrained state. (C) Expression and schematic of expression pattern of the R38E07 FSB driver shows expression in regions 5, 8, and 9. (D) UAS-shi^ts^ crossed to R38E07 trained by *D. ananassae* at the permissive and restrictive temperature shows wild-type naïve state. (E) Expression and schematic of expression pattern of the R89E07 FSB driver shows expression in regions 2, 8, and 9. (F) UAS-shi^ts^ crossed to R89E07 trained by *D. ananassae* at the permissive and restrictive temperature shows wild-type untrained state. (G) Expression and schematic of expression pattern of the R49H02 FSB driver shows expression in regions 1, 4, and 6. (H) UAS-shi^ts^ crossed to R49H02 trained by *D. ananassae* at the permissive and restrictive temperature shows wild-type naïve state. Error bars represent standard error (n = 12 biological replicates) (*p < 0.05).

**Supplementary Figure 32.**
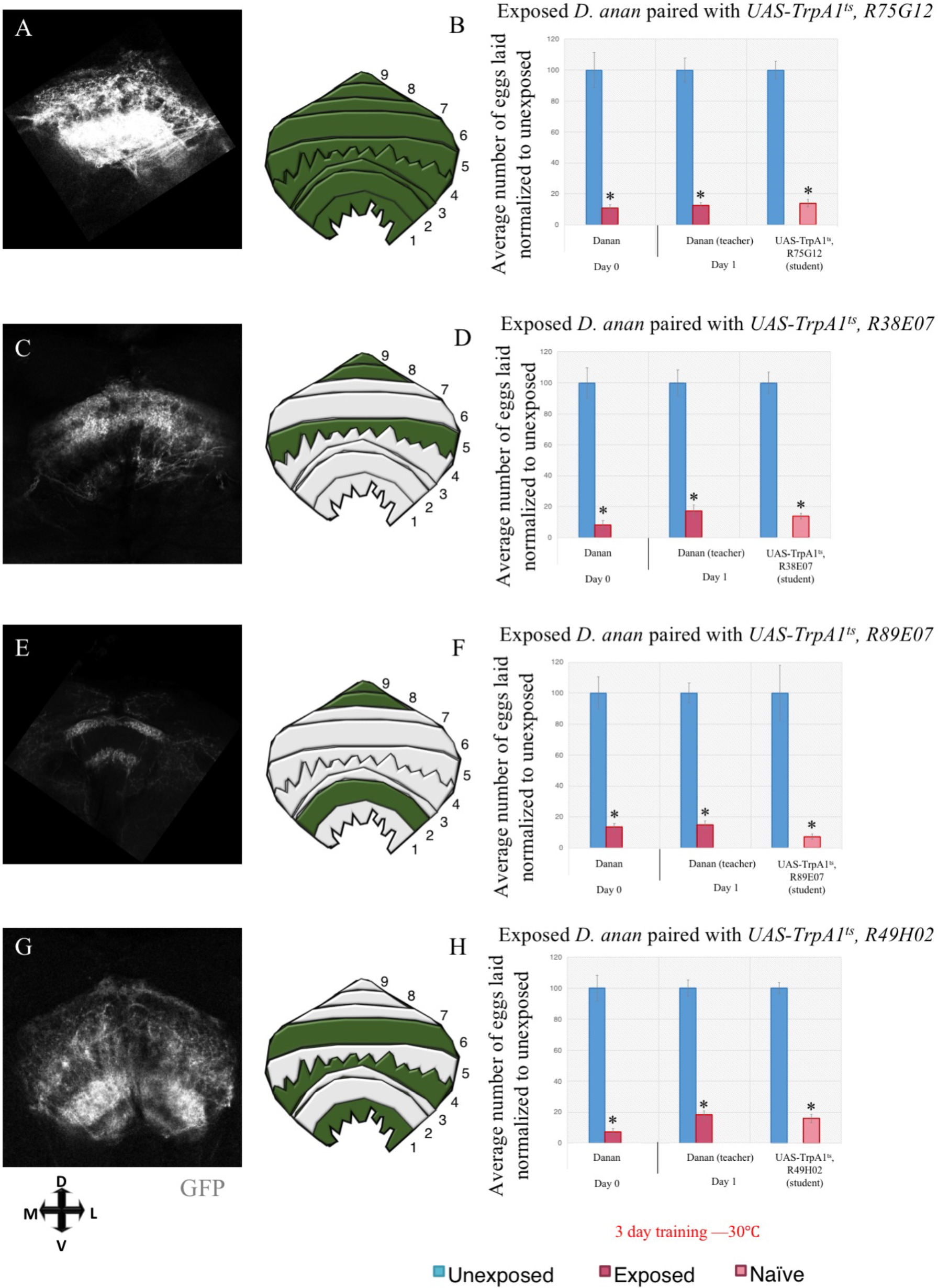
Further evidence indicating that Region 5 of the fan-shaped body is sufficient for dialect learning. Dialect learning for this experiment series is performed for 3 days at the active (30°C) temperature, while the wasp exposure and social learning period is performed exclusively at the restrictive temperature. Percentage of eggs laid by exposed flies normalized to eggs laid by unexposed flies is shown. (A) Expression and schematic of expression pattern of the R75G12 FSB driver shows pan-FSB marking. (B) UAS-shi^ts^ crossed to R75G12 trained by *D. ananassae* for 3 days at the permissive temperature shows wild-type trained state. (C) Expression and schematic of expression pattern of the R38E07 FSB driver shows expression in regions 5, 8, and 9. (D) UAS-shi^ts^ crossed to R38E07 trained by *D. ananassae* for 3 days at the permissive temperature shows wild-type trained state. (E) Expression and schematic of expression pattern of the R89E07 FSB driver shows expression in regions 2, 8, and 9. (F) UAS-shi^ts^ crossed to R89E07 trained by *D. ananassae* for 3 days at the permissive temperature shows wild-type trained state. (G) Expression and schematic of expression pattern of the R49H02 FSB driver shows expression in regions 1, 4, and 6. (H) UAS-shi^ts^ crossed to R49H02 trained by *D. ananassae* for 3 days at the permissive temperature shows wild-type trained state. Error bars represent standard error (n = 12 biological replicates) (*p < 0.05).

**Supplementary Figure 33.**
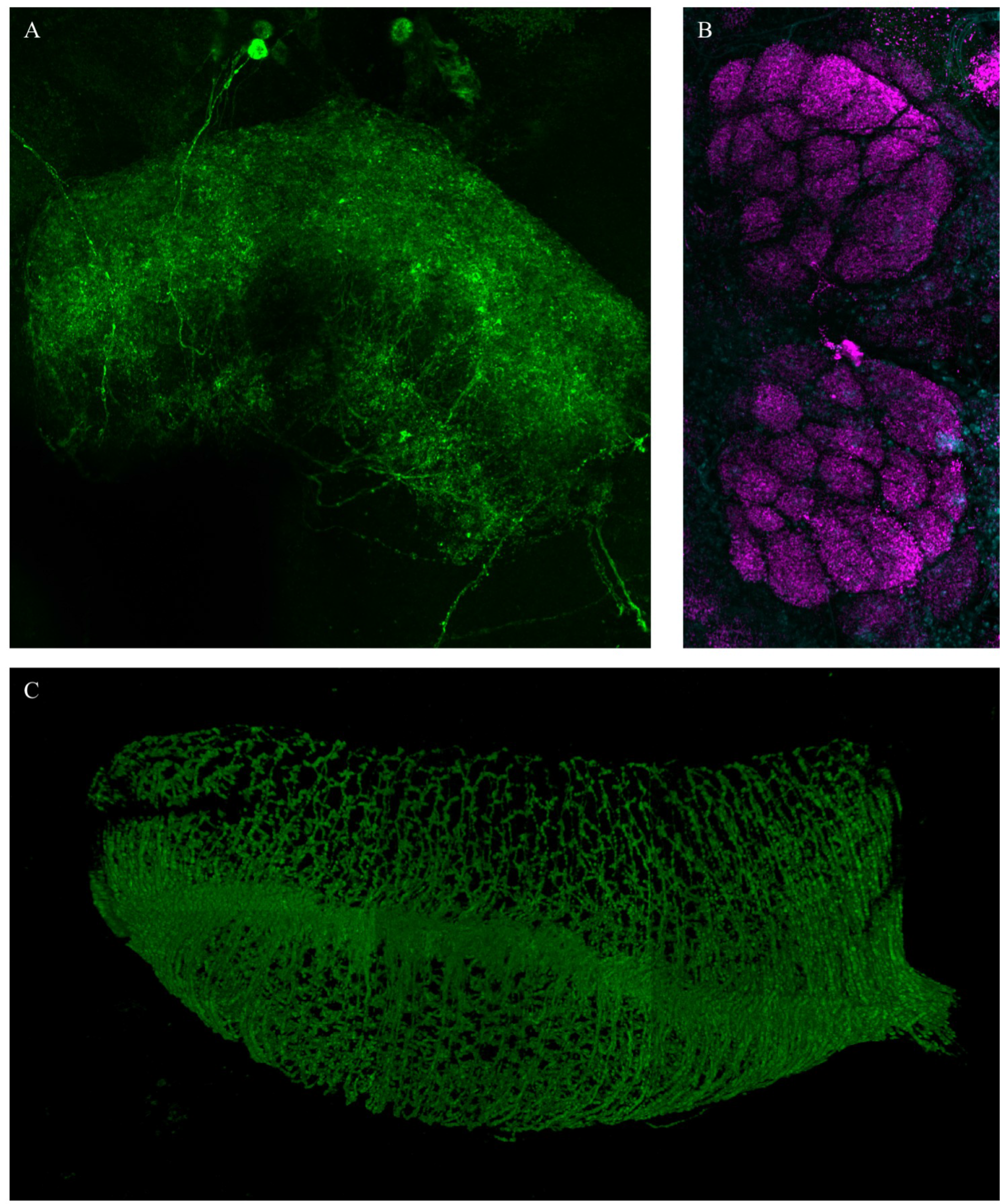
Super resolution microscopy of key brain regions in dialect learning. (A) Z-stack maximum intensity project super resolution image of driver line R38E07 in conjunction with UAS-CD8-GFP. (B) Z-stack maximum intensity project super resolution image of the antennal lobe stained with nc82. (C) Z-stack maximum intensity project super resolution image of driver line L4^0987^ in conjunction with UAS-CD8-GFP.

## SUPPLEMENTARY TABLE LEGENDS

**Supplementary Table 1.**
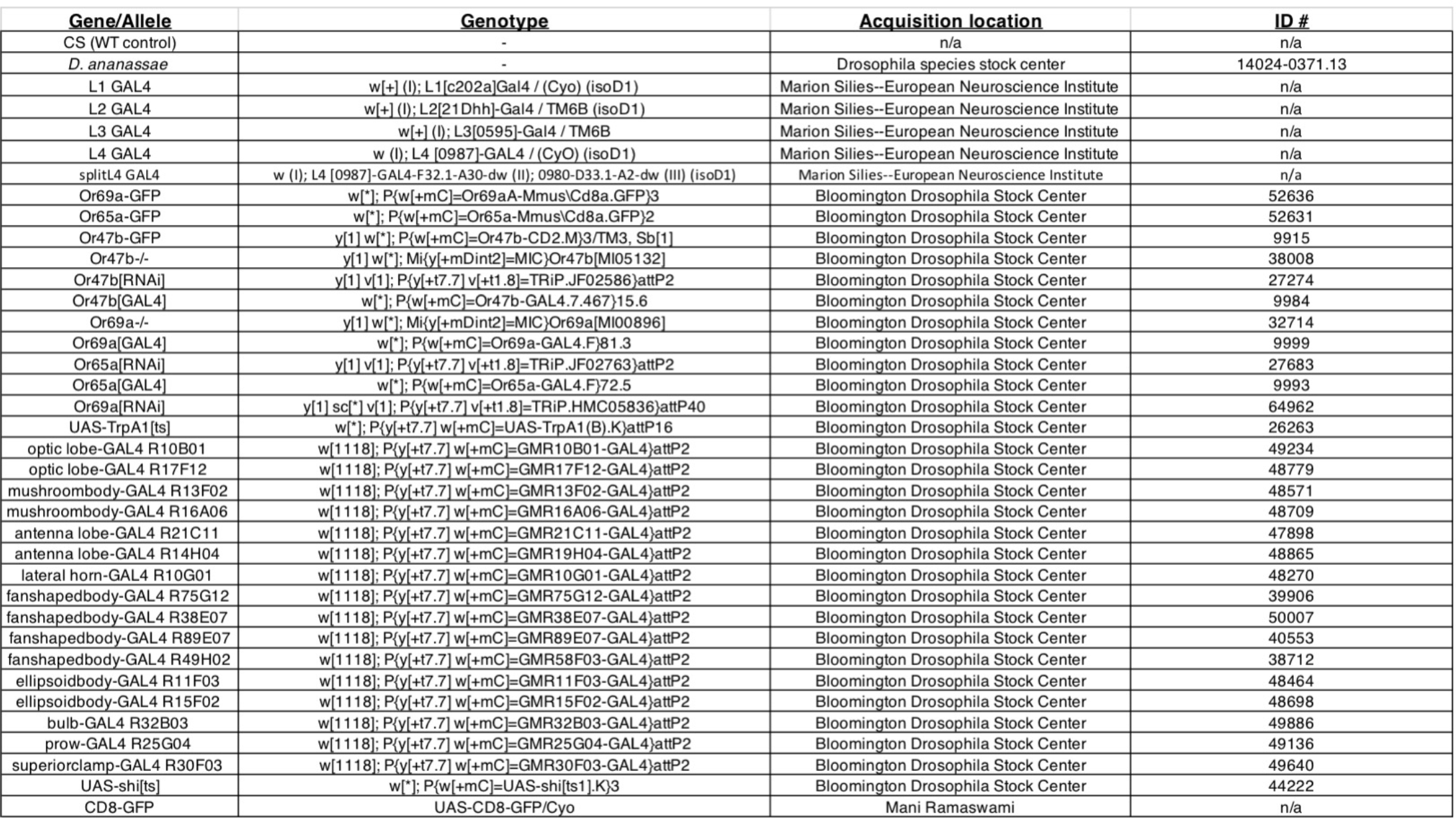
Name, genotype, acquisition location, and stock identification number (if applicable) are shown.

| Gene/Allele | Genotype | Acquisition location | ID # |
| --- | --- | --- | --- |
| CS (WT control) | - | n/a | n/a |
| *D. ananassae* | - | Drosophila species stock center | 14024-0371.13 |
| L1 GAL4 | w[+] (I); L1[c202a]Gal4 / (Cyo) (isoD1) | Marion Silies--European Neuroscience Institute | n/a |
| L2 GAL4 | w[+] (I); L2[21Dhh]-Gal4 / TM6B (isoD1) | Marion Silies--European Neuroscience Institute | n/a |
| L3 GAL4 | w[+] (I); L3[0595]-Gal4 / TM6B | Marion Silies--European Neuroscience Institute | n/a |
| L4 GAL4 | w (I); L4 [0987]-GAL4 / (Cyo) (isoD1) | Marion Silies--European Neuroscience Institute | n/a |
| splitL4 GAL4 | w (I); L4 [0987]-GAL4-F32.1-A30-dw (II); 0980-D33.1-A2-dw (III) (isoD1) | Marion Silies--European Neuroscience Institute | n/a |
| Or69a-GFP | w[*]; P(w[+mC]=Or69aA-Mmus)Cd8a.GFP)3 | Bloomington Drosophila Stock Center | 52636 |
| Or65a-GFP | w[*]; P(w[+mC]=Or65aA-Mmus)Cd8a.GFP)2 | Bloomington Drosophila Stock Center | 52631 |
| Or47b-GFP | y[1] w[*]; P(w[+mC]=Or47b-CD2.M)3/TM3, Sb[1] | Bloomington Drosophila Stock Center | 9915 |
| Or47b-/- | y[1] w[*]; Mi(y[+mDint2]=MIC)Or47b[Mi05132] | Bloomington Drosophila Stock Center | 38008 |
| Or47b[RNAi] | y[1] v[1]; P(y[+t7.7] v[+t1.8]=TRiP.JF02586)attP2 | Bloomington Drosophila Stock Center | 27274 |
| Or47b[GAL4] | w[*]; P(w[+mC]=Or47b-GAL4.7.467)15.6 | Bloomington Drosophila Stock Center | 9984 |
| Or69a-/- | y[1] w[*]; Mi(y[+mDint2]=MIC)Or69a[Mi00896] | Bloomington Drosophila Stock Center | 32714 |
| Or69a[GAL4] | w[*]; P(w[+mC]=Or69a-GAL4.F)81.3 | Bloomington Drosophila Stock Center | 9999 |
| Or65a[RNAi] | y[1] v[1]; P(y[+t7.7] v[+t1.8]=TRiP.JF02763)attP2 | Bloomington Drosophila Stock Center | 27683 |
| Or65a[GAL4] | w[*]; P(w[+mC]=Or65a-GAL4.F)72.5 | Bloomington Drosophila Stock Center | 9993 |
| Or69a[RNAi] | y[1] sc[*] v[1]; P(y[+t7.7] v[+t1.8]=TriP.HMC05836)attP40 | Bloomington Drosophila Stock Center | 64962 |
| UAS-TrpA1[ts] | w[*]; P(y[+t7.7] w[+mC]=UAS-TrpA1(B).K)attP16 | Bloomington Drosophila Stock Center | 26263 |
| optic lobe-GAL4 R10B01 | w[1118]; P(y[+t7.7] w[+mC]=GMR10B01-GAL4)attP2 | Bloomington Drosophila Stock Center | 49234 |
| optic lobe-GAL4 R17F12 | w[1118]; P(y[+t7.7] w[+mC]=GMR17F12-GAL4)attP2 | Bloomington Drosophila Stock Center | 48779 |
| mushroombody-GAL4 R13F02 | w[1118]; P(y[+t7.7] w[+mC]=GMR13F02-GAL4)attP2 | Bloomington Drosophila Stock Center | 48571 |
| mushroombody-GAL4 R16A06 | w[1118]; P(y[+t7.7] w[+mC]=GMR16A06-GAL4)attP2 | Bloomington Drosophila Stock Center | 48709 |
| antenna lobe-GAL4 R21C11 | w[1118]; P(y[+t7.7] w[+mC]=GMR21C11-GAL4)attP2 | Bloomington Drosophila Stock Center | 47898 |
| antenna lobe-GAL4 R14H04 | w[1118]; P(y[+t7.7] w[+mC]=GMR19H04-GAL4)attP2 | Bloomington Drosophila Stock Center | 48865 |
| lateral horn-GAL4 R10G01 | w[1118]; P(y[+t7.7] w[+mC]=GMR10G01-GAL4)attP2 | Bloomington Drosophila Stock Center | 48270 |
| fan-shapedbody-GAL4 R75G12 | w[1118]; P(y[+t7.7] w[+mC]=GMR75G12-GAL4)attP2 | Bloomington Drosophila Stock Center | 39906 |
| fan-shapedbody-GAL4 R38E07 | w[1118]; P(y[+t7.7] w[+mC]=GMR38E07-GAL4)attP2 | Bloomington Drosophila Stock Center | 50007 |
| fan-shapedbody-GAL4 R89E07 | w[1118]; P(y[+t7.7] w[+mC]=GMR89E07-GAL4)attP2 | Bloomington Drosophila Stock Center | 40553 |
| fan-shapedbody-GAL4 R49H02 | w[1118]; P(y[+t7.7] w[+mC]=GMR58F03-GAL4)attP2 | Bloomington Drosophila Stock Center | 38712 |
| ellipsoidbody-GAL4 R11F03 | w[1118]; P(y[+t7.7] w[+mC]=GMR11F03-GAL4)attP2 | Bloomington Drosophila Stock Center | 48464 |
| ellipsoidbody-GAL4 R15F02 | w[1118]; P(y[+t7.7] w[+mC]=GMR15F02-GAL4)attP2 | Bloomington Drosophila Stock Center | 48698 |
| bulb-GAL4 R32B03 | w[1118]; P(y[+t7.7] w[+mC]=GMR32B03-GAL4)attP2 | Bloomington Drosophila Stock Center | 49886 |
| prow-GAL4 R25G04 | w[1118]; P(y[+t7.7] w[+mC]=GMR25G04-GAL4)attP2 | Bloomington Drosophila Stock Center | 49136 |
| superiorclasp-GAL4 R30F03 | w[1118]; P(y[+t7.7] w[+mC]=GMR30F03-GAL4)attP2 | Bloomington Drosophila Stock Center | 49640 |
| UAS-shi[ts] | w[*]; P(w[+mC]=UAS-shi[ts1].K)3 | Bloomington Drosophila Stock Center | 44222 |
| CD8-GFP | UAS-CD8-GFP/Cyo | Mani Ramaswami | n/a |

## SUPPLEMENTARY MOVIE LEGENDS

**Supplementary Movie 1**. Expression pattern of the L4^0987^ line in whole brain.

Super resolution confocal movie of adult Drosophila brain where L4^0987^ GAL4 is driving UAS-CD8-GFP, stained with DAPI (teal) and nc82 (magenta).

**Supplementary Movie 2**. Expression pattern of the L4^0987^ line in optic lobe.

Super resolution confocal movie of adult Drosophila brain where L4^0987^ GAL4 is driving UAS-CD8-GFP, stained with DAPI (teal) and nc82 (magenta).

**Supplementary Movie 3**. Expression pattern of the fan-shaped body driver line R38E07.

Super resolution confocal movie of adult Drosophila brain where R38E07 is driving UAS-CD8-GFP, stained with DAPI (teal).

**Supplementary Movie 4**. Expression pattern of the fan-shaped body driver line R38E07 in a magnified region.

Super resolution confocal movie of adult Drosophila brain where R38E07 is driving UAS-CD8-GFP, stained with DAPI (teal).

## SUPPLEMENTARY FILE LEGENDS

**Supplementary Table 1**. Raw egg counts and p values for figures used in this study. Each tab corresponds to a given panel series. Within tab contains the genotype of construct(s) used, raw egg counts, and p-values.

